# 2′-*O*-Methylated ncRNA condensates form a recombination focus for *IgH* synapsis

**DOI:** 10.64898/2026.11.05.761835

**Authors:** Maki Kobayashi, Jianliang Xu, Afzal Husain, Takahito Moriwaki, Ziwei Yin, Xi Chen, Wencong Wang, Toshitsugu Fujita, Hodaka Fujii, Luthfi Lulul Ulum, Takanori Oyoshi, Kanon Omura, Kei Iida, Hidetaka Kosako, Tasuku Honjo

## Abstract

While FBL-mediated RNA 2′-*O*-methylation has been linked to class switch recombination (CSR), how this epitranscriptomic mark orchestrates higher-order genomic architecture remains unknown. Here, we demonstrate that FBL selectively promotes switch (*S*)–*S* DNA synapsis at the *IgH* locus during CSR, without affecting DNA cleavage. Both RNA-binding and catalytic activities of FBL were essential for CSR. Moreover, MED19 supported DNA cleavage and synapsis by regulating snoRNAs guiding 2′-*O*-methylation. Mechanistically, FBL methylated *α*-germline transcripts (*α*-GLTs), *cis*-acting noncoding RNAs forming nuclear condensates that compartmentalize RNA processing and chromatin-remodeling factors. FBL-mediated 2′-*O*-methylation governed condensate protein composition; FBL suppression aberrantly enriched nucleolar proteins, notably WDR74. Low WDR74 expression in B cells promoted *μ*- and *α*-GLT condensate fusion, enhancing *S*–*S* synapsis, CSR, and *IgH*/*c-Myc* translocation, potentially contributing to poor myeloma prognosis. Altogether, RNA 2′-*O*-methylation dictates GLT condensate composition and fusion dynamics to establish a recombination focus, driving genomic DNA synapsis in B cells.

## INTRODUCTION

In general, alteration of genetic information in somatic cells is strictly suppressed because of its risk of tumorigenesis. However, activated B lymphocytes routinely undergo programmed DNA recombination events, such as somatic hypermutation (SHM) and class switch recombination (CSR), to produce high-affinity antibodies against diverse pathogens.^1–3^ Although cells maintain genome stability by preventing translocation and oncogenesis, the mechanism by which they rapidly and accurately restrict these DNA recombination events specifically to the *IgH* locus remains a fundamental question in adaptive immunity.

The *IgH* locus provides an ideal model for decoding complex recombination systems. This locus contains unique domains of highly repetitive switch (*S*) regions and noncoding transcription units, namely germline transcripts (GLTs), that originate from distinct *I* promoters and span the *S* regions.^4–6^ CSR comprises three sequential steps: DNA cleavage at two *S* regions, spatial juxtaposition of these distant regions (*S*–*S* DNA synapsis), and the final joining of DNA break ends via nonhomologous end-joining (NHEJ).^3,7^ Because the target *S* regions are separated by more than 100 kb, *S*–*S* synapsis relies on higher-order chromatin looping orchestrated by the cohesin complex,^8^ and this process is regulated by the 3′ regulatory region (3′RR) via enhancer RNA (eRNA) and transcriptional cofactors.^9–11^ This dynamic proximity must be strictly oriented. Productive CSR requires deletional recombination, whereas inverted recombination impairs CSR and B-cell receptor expression, ultimately leading to cell death. Furthermore, aberrant synapsis or recombination involving *Sμ* and 3′RR or the *c-Myc* locus can trigger locus suicide recombination or chromosomal translocations, respectively.^12,13^ Indeed, aberrant DNA synapsis leading to *IgH*/*c-Myc* translocation depending on activation-induced cytidine deaminase (AID) drives B lymphomagenesis^14^ and predicts poor clinical outcomes.^15^

AID is essential for CSR,^1,2^ and its C-terminal domain is critically required for *S*–*S* synapsis without affecting DNA cleavage or SHM.^16–19^ However, the function of its C-terminal domain remains incompletely defined. RNA-dependent regulation of *S*–*S* synapsis is highly plausible, as cis-acting long noncoding RNAs regulate gene expression through associations with enhancers and promoters or the recruitment of gene-looping factors.^20^ At the *IgH* locus, cis-acting RNAs, including eRNAs and GLTs, play pivotal roles. Specifically, eRNAs derived from 3′RR facilitate higher-order chromatin looping and synapsis,^10,11^ whereas GLTs form G-quadruplex (G4) structures that recruit AID to *IgH* regions.^21^ Furthermore, GLTs produced from *S* regions form dynamic hubs that recruit AID to *IgH* regions,^22^ and *m*^6^A modification of GLTs by METTL3 is required for RNA exosome-mediated processing and CSR.^23^ Recently, fibrillarin (FBL), a conserved 2′-*O*-methyltransferase involved in rRNA processing and ribosome biogenesis,^24,25^ was reported to influence CSR by promoting local R-loop formation and AID recruitment.^26^

However, the molecular mechanism governing these synaptic GLT-derived RNA hubs,^22^ specifically the processes regulating their physicochemical properties and protein composition during long-range *S*–*S* synapsis, remains largely unknown. To elucidate the molecular machinery operating directly at the *Sα* DNA region, our unbiased insertional chromatin immunoprecipitation (iChIP) screening platform identified FBL as a key locus-specific factor, and its catalytic activity proved essential for CSR. Furthermore, by applying CRISPR-mediated *Sirius-8xMS2* tagging at *Sα*-GLT transcripts, we systematically defined the distinct GLT-bound interactome and its condensate dynamics. We demonstrated that FBL-mediated 2′-*O*-methylation shapes the protein composition of GLT condensates, thereby tuning their fusion dynamics. Collectively, our findings reveal a fundamental mechanism by which RNA 2′-*O*-methylation governs condensate fusion to drive long-range DNA synapsis in B cells.

## RESULTS

### *Sα*-specific proteomics reveals enrichment of nucleolar proteins

To isolate proteins specifically recruited to the *Sα* region, we inserted 8xLexA-binding sequences into the CH12 cell genome, as previously reported^27,28^ (Figures 1A and S1). After expressing LexA protein tagged with 3xFLAG and a nuclear localization signal (FNLDD), cells were designated “FNLDD-8xLexA-*Sα*-core” (control cells expressing FNLDD alone were termed “FNLDD-alone”). Chromatin immunoprecipitation (ChIP) confirmed the specific enrichment of 8xLexA in the *Sα* region (Figure S1H). We compared the FNLDD-recruited protein fractions between FNLDD-8xLexA-*Sα*-core and FNLDD-alone cells using stable isotope labeling by amino acids in cell culture (SILAC) coupled with mass spectrometry (MS; Figure 1B).

**Figure 1.**
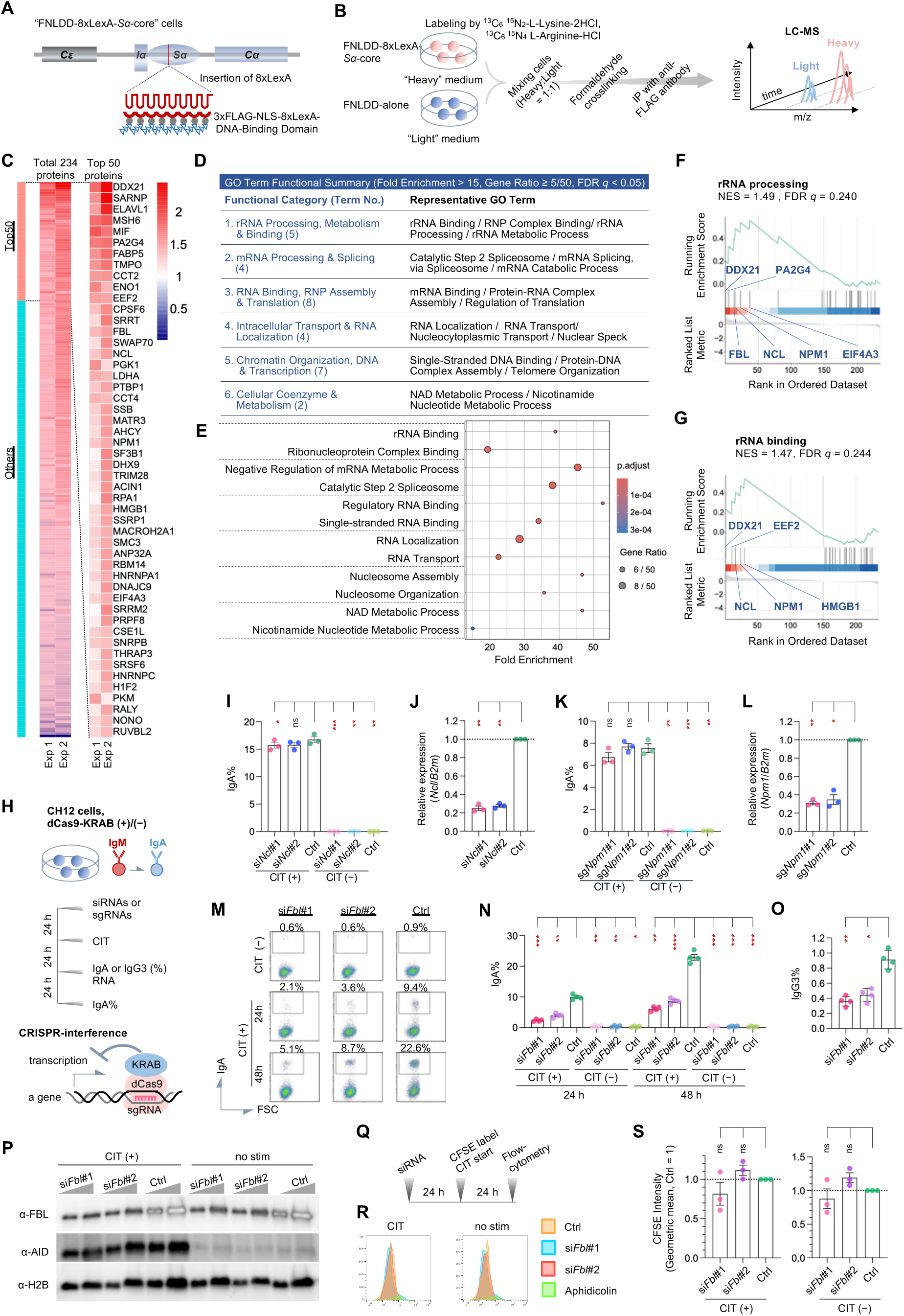
LexA-mediated *Sα-*core DNA interactome proteomics identified multiple nucleolar proteins required for class switch recombination. (A) Schematic of the *IgHα* locus in CH12F3-2A cells carrying an 8xLexA binding element. (B) Workflow of insertional ChIP (iChIP) coupled with SILAC-based mass spectrometry (iChIP-MS). (C–G) Proteomic profiling of the *Sα*-core interactome, displaying heatmaps of the overall and top 50 *Sα*-enriched proteins across two independent experiments (C), a functional summary table (D) and a dot plot (E) from Gene Ontology (GO) enrichment analysis, and GSEA enrichment plots (F and G). False discovery rate (FDR) *q*-value, “Fold Enrichment”, and “Gene Ratio” were used to filter GO terms in enrichGO and GSEA. (H) Timeline for siRNA or CRISPRi knockdown. (I–L) IgA switching efficiency (I, K) and knockdown efficiency (J, L) for *Ncl* and *Npm1* (*n* = 3). (M–P) Flow cytometric profiles (M), quantification of IgA^+^ (N) and IgG3^+^ (O) cells (*n* = 4), and western blot (P) in *Fbl-*knockdown cells. H2B served as a loading control. (Q–S) Cell proliferation timeline (Q), CFSE profiles (R), and quantification (S) in *Fbl*-knockdown cells (*n* = 3). Data represent the mean ± SEM. Dashed horizontal lines indicate the normalized control levels (1.0). \**P*_adj <_ 0.05, \*\**P*_adj <_ 0.01, \*\*\**P*_adj <_ 0.001, \*\*\*\**P*_adj <_ 0.0001; ns, not significant. See STAR Methods for statistical details.

Across two independent MS experiments identifying 234 proteins, we extracted the top 50 factors displaying the highest “Heavy/Light” ratios in FNLDD-8xLexA-*Sα*-core cells (Figure 1C; Data S2). Pathway analysis of these 50 factors using clusterProfiler v4.14.6^29^ was performed. To feature the primary GO terms, threshold criteria of Fold Enrichment > 15, Gene Ratio ≥ 5/50, and false discovery rate (FDR) q-value < 0.05 were applied, highlighting rRNA processing, mRNA processing and splicing, ribonucleoprotein (RNP) assembly and translation, RNA localization, chromatin organization, and cellular coenzymes (Figures 1D and 1E; Data S3).

Gene set enrichment analysis (GSEA) of 234 proteins revealed that the strongly enriched GO terms were predominantly linked to RNA/mRNA metabolic processes and splicing (Data S4). This profile contrasts with our previous findings, in which 8xLexA, inserted downstream of *Sα*, revealed enrichment for DNA repair and chromatin factors, underscoring the distinct RNA-centric microenvironment of the *Sα*-core locus.^28^ Notably, alongside the mRNA/RNA processing terms, nucleolar-related functional categories such as “GOBP_RRNA_PROCESSING” (normalized enrichment score [NES] = 1.49 and FDR *q*-value = 0.240) and “GOMF_RRNA_BINDING” (NES = 1.47 and FDR *q*-value = 0.244) were enriched within the exploratory threshold of FDR *q*-value < 0.25 (Figures 1F-1G).

### The 2′-*O*-methyltransferase Fibrillarin (FBL) is specifically required for CSR

Based on these findings, we selected FBL, nucleolin (NCL), and nucleophosmin 1 (NPM1), all well-established markers for nucleolar enrichment, as candidates for functional evaluation in CH12 cells using siRNA or CRISPR interference (CRISPRi; Figure 1H).^30^ Twenty-four hours post-stimulation with CD40 ligand, IL-4, and TGFβ (CIT), cells were evaluated for CSR to IgA or IgG3 (Figures 1I, 1K, and 1M–1O) and target expression (Figures 1J, 1L, and 1P). Notably, CSR to IgA and IgG3 was significantly impaired exclusively upon *Fbl* knockdown.

Because FBL is essential for rRNA maturation and cell cycle progression,^31,32^ high siRNA concentrations cause proliferation defects. To bypass secondary effects, we titrated the siRNA concentrations to achieve mild knockdown without altering cell cycle kinetics and used *Bcl2*-overexpressing CH12 cells to prevent background apoptosis. Under these conditions, *Fbl* knockdown did not alter AID expression (Figure 1P). Normal cell cycle progression in *Fbl*-knockdown cells was confirmed using carboxyfluorescein succinimidyl ester (CFSE) dilution profiles (Figures 1Q-1S).

Furthermore, FBL exhibited distinct properties from NCL and NPM1 (Figure S2). Unlike NCL and NPM1, which translocated to etoposide-induced γH2AX DNA damage foci, EGFP-FBL remained excluded from the γH2AX regions (Figure S2A). Co-immunoprecipitation (co-IP) demonstrated that FBL specifically interacts with AID via its N- and C-terminal domains, whereas NCL and NPM1 did not interact with AID (Figures S2B–S2H). Crucially, RNase A treatment severely reduced the FBL–AID interaction, demonstrating that FBL associates with AID in an RNA-dependent manner (Figure S2I).

### FBL drives DNA synapsis independently of DNA cleavage

To pinpoint the site at which FBL acts within the multistep cascade of CSR, we systematically evaluated each sequential stage (Figure 2A). *Fbl* knockdown did not alter *Aicda* or GLT expression (Figures 2B-2C). Sequencing of the 5′ *Sμ* region revealed that FBL downregulation did not affect SHM rates (Figures 2D-2F; Tables S1 and S2). Direct DNA 3′-end labeling with biotin-dUTP further confirmed that DNA cleavage at the S regions remained intact (Figures 2G-2H).

**Figure 2.**
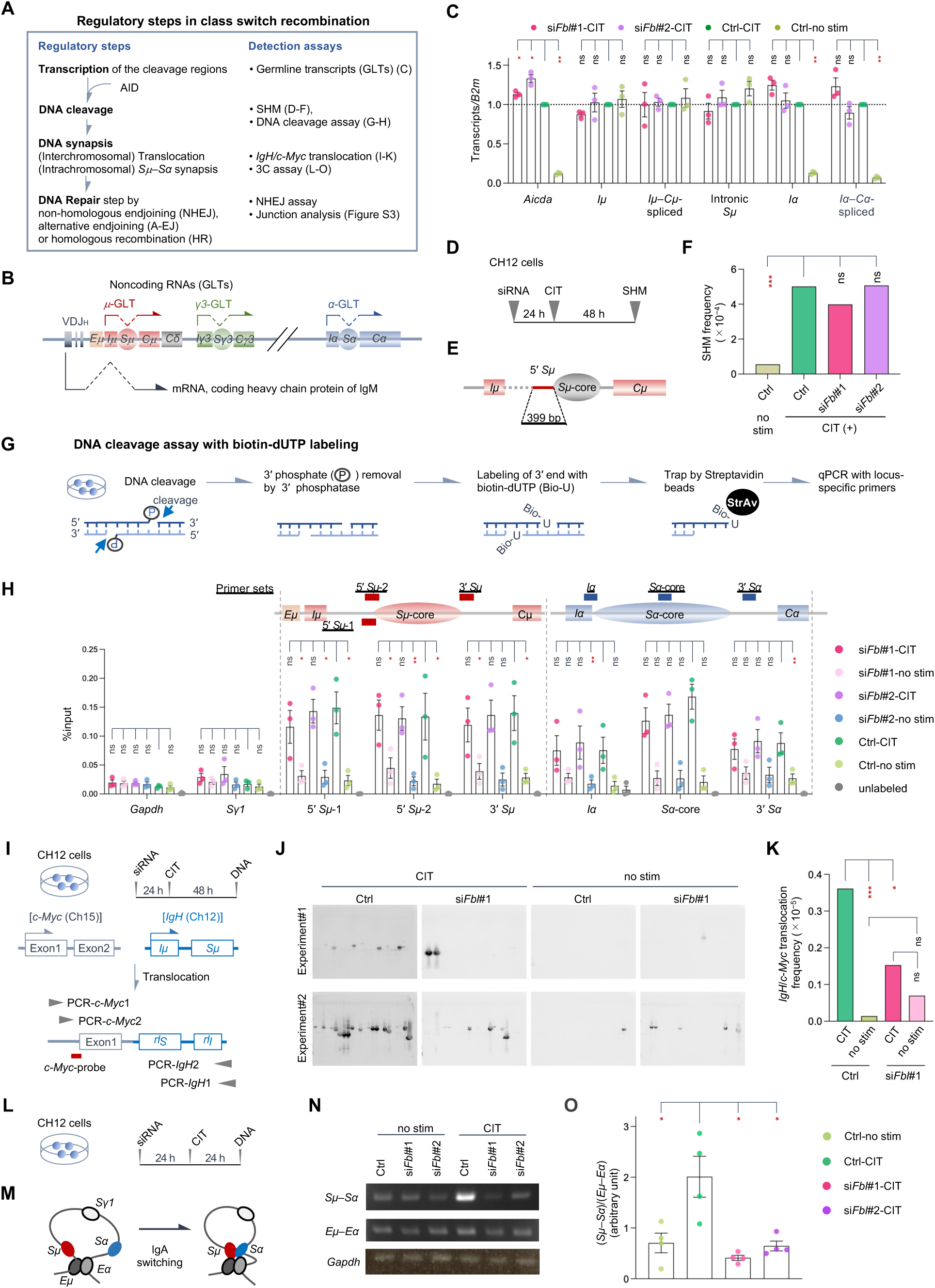
*Fbl* knockdown disrupts long-range *IgH* synapsis and translocation without impairing DNA cleavage frequency. (A) Schematic summary of the regulatory steps in the CSR cascade. (B) Schematic representation of germline transcript (GLT) gene structures, indicating *I* promoters, intronic switch (*S*) regions, and constant (*C*) regions. (C) Reverse transcription-qPCR (RT-qPCR) analysis of *Aicda*, *μ*- and *α-*GLTs in *Fbl-*knockdown cells (*n* = 3). (D–F) Somatic hypermutation (SHM) analysis: timeline (D), 5′ *Sμ*-core locus map (E), and SHM frequencies (F) in *Fbl-*knockdown cells. See Tables S1 and S2. (G and H) Biotin–dUTP (Bio-U) DNA cleavage assay workflow (G) and cleavage-qPCR at the *IgH* locus (H) (*n* = 3) in *Fbl-*knockdown cells. (I–K) *IgH*/*c-Myc* translocation analysis: schematic map (I), Southern blot (J), and translocation frequencies (K) in *Fbl*-knockdown cells. (L–O) 3C assay evaluating *Sμ*–*Sα* proximity: timeline (L), topology scheme (M), representative agarose gels (N), and quantified *Sμ*–*Sα* interaction frequencies (O). *n* = 4. Data represent the mean ± SEM, except in (F) and (K), where values represent calculated frequencies. Dashed horizontal lines indicate the normalized control value (1.0). \**P*_adj <_ 0.05, \*\**P*_adj <_ 0.01, \*\*\**P*_adj <_ 0.001; ns, not significant. See STAR Methods for statistical details.

In contrast, spatial DNA proximity was severely disrupted. Both *IgH*/*c-Myc* translocation frequency (Figures 2I-2K) and *Sμ*–*Sα* DNA synapses, measured by chromosome conformation capture (3C) (Figures 2L-2O), were markedly reduced by *Fbl* knockdown. NHEJ activity remained unimpaired, as demonstrated by artificial reporter assays (H1299dA3-1 cells;^33^ Figures S3A–S3F) and junction microhomology profiling in CH12 cells (Figures S3G-S3I; Data S5). Together, these data established that FBL specifically governs spatial DNA proximity, manifested as *S*–*S* synapses and *IgH*/*c-Myc* translocation, without altering transcription, DNA cleavage, or end-joining repair.

### FBL targets G-quadruplex structures in germline transcripts to facilitate CSR

FBL functions primarily as an RNA 2′-*O*-methyltransferase.^34,35^ Using available structural data (PDB: 2IPX)^36^ and the KYG algorithm,^37^ we generated RNA-binding mutants outside the nucleolar-targeting glycine–arginine-rich (GAR) domain^38^ (Figures 3A and S4A). In knockdown-rescue experiments, all RNA-binding mutants, including the most severely affected ΔGAR91 and ΔGAR195, failed to rescue CSR (Figures 3B, S4B, and S4C). RNA immunoprecipitation (RIP) confirmed reduced GLT binding across these mutants compared with the findings in the wild-type (WT), with ΔGAR195 displaying a statistically significant loss of binding (Figures 3C, 3D, and S4D). Surprisingly, ChIP revealed no remarkable peaks at *IgH* DNA (Figures S4E–S4F), indicating that FBL detection during initial MS analysis reflected the spatial proximity of FNLDD to GLT RNA rather than a direct chromatin association.

**Figure 3.**
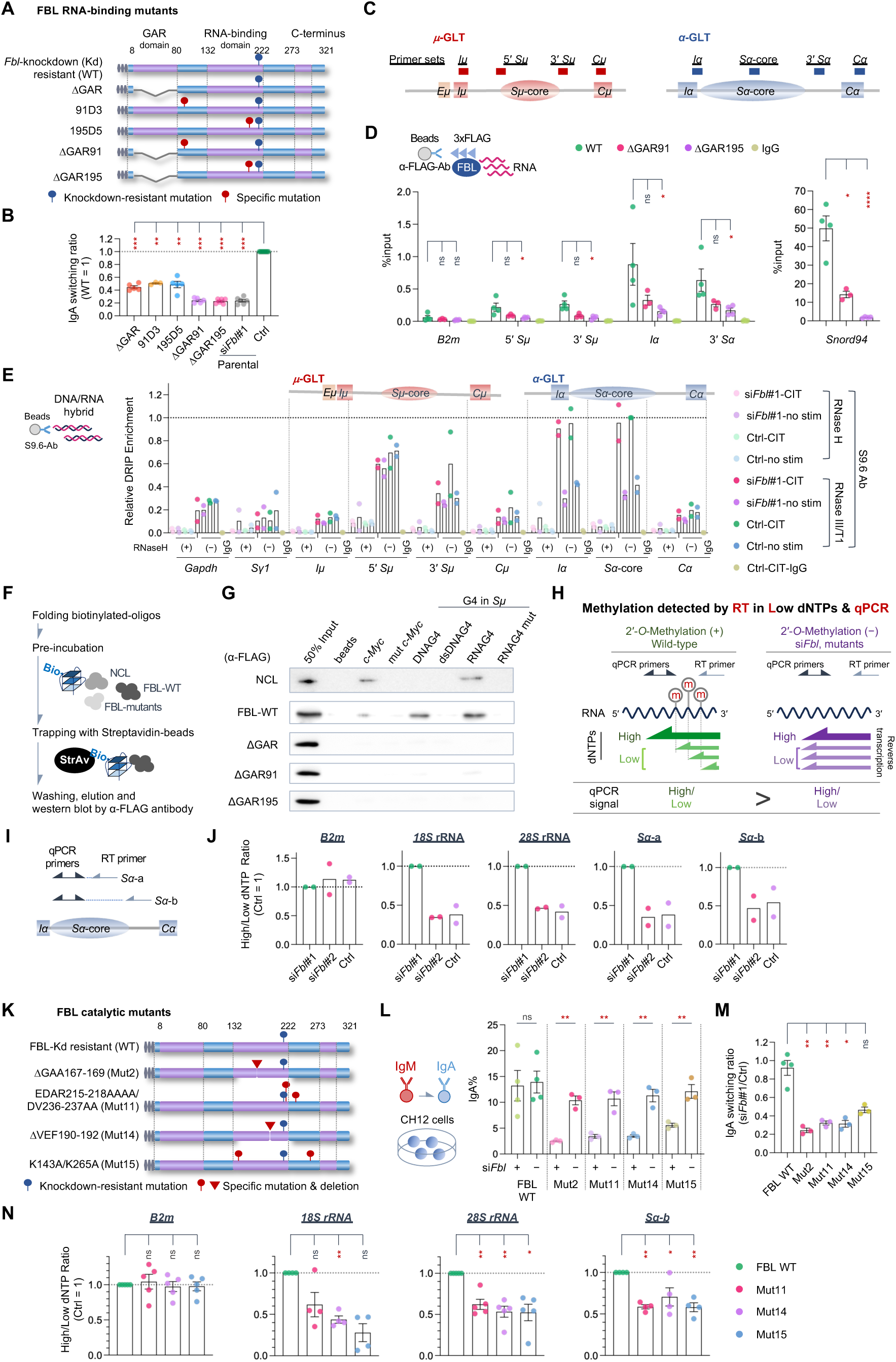
FBL promotes CSR through G4-binding and 2′-*O*-methyltransferase activity. (A) Schematic of the wild-type (WT) and RNA-binding mutants of human FBL. (B) IgA switching efficiency of FBL RNA-binding mutants relative to that of WT (*n* = 3–11). (C–E) Primer positions in the *IgH* locus (C), RNA-IP–qPCR of *IgH* transcripts (D; *n* = 3–4), and DRIP-qPCR of R-loops (E; *n* = 2) in FBL mutants. In (E), values were normalized to the Ctrl-CIT conditions treated with RNase III/T1 in the *Sα*-core (set to 1). See also Figure S4D. (F and G) Biotinylated RNA and DNA pulldown workflow (F) and western blot (G) assessing FBL binding to canonical switch-region G4 structures derived from *S*μ. (H) Principle of RTL-qPCR for detecting 2′-*O*-methylation (ⓜ). Low dNTP conditions stall reverse transcription (RT) elongation specifically at methylated sites (ⓜ) in WT cells, whereas hypomethylation (si*Fbl* or mutants) permits unhindered RT. Consequently, a lower “High/Low dNTP ratio” reflects reduced 2′-*O*-methylation levels. (I) Primer positions for reverse transcription and qPCR in the *Sα* region for RTL-qPCR. (J) RTL-qPCR analysis for *B2m*, 18s/28s rRNA, *Sα*-a, and *Sα*-b in *Fbl* knockdown cells. *n* = 2. (K–N) FBL methyltransferase mutants: domain structures (K), IgA switching efficiencies (L and M; *n* = 3–4), and RTL-qPCR rescue analysis (N; *n* = 4–5). Data represent the mean ± SEM, except in (E) and (J), where individual values are plotted as dots. Dashed horizontal lines indicate normalized control levels (1.0). \**P*_adj <_ 0.05, \*\**P*_adj <_ 0.01 (\*\**P* < 0.01 for Figure 3L), \*\*\**P*_adj <_ 0.001, \*\*\*\**P*_adj <_ 0.0001; ns, not significant. See STAR Methods for statistical details.

DNA/RNA immunoprecipitation (DRIP)-qPCR assays using the S9.6 antibody revealed that *Fbl* knockdown did not markedly alter R-loop levels across the *μ*- and *α*-GLT loci (Figure 3E). This indicates that FBL functions primarily on a posttranscriptional nuclear pool of GLTs rather than directly targeting nascent, DNA-hybridized R-loops.

Because GAR or repeat of arginine–glycine–glycine (RGG) domains frequently recognize G4 structures and GAR domain-containing FBL is known to interact with G4-forming TERRA RNA,^39^ we hypothesized that FBL targets G4 structures within GLTs. Indeed, computational analysis using G4RNA screener^40^ predicted multiple G4 motifs at comparable levels in both *Sμ* and *Sα* regions (Figures S4G–S4H). Given that *Sμ* and *Sα* regions share conserved G-rich pentameric repeat frameworks^3^ with comparable G4-folding capacities, we utilized canonical *Sμ* G4 motifs as representative targets. *In vitro* RNA and DNA pulldown assays confirmed that FBL-WT bound robustly to G4-containing *Sμ* RNA and *Sμ DNA* (RNAG4 and DNAG4, respectively), whereas RNA-binding mutants failed to interact (Figures 3F, 3G, and S4I). Furthermore, treatment with the G4-stabilizing ligand pyridostatin (PDS) inhibited CSR in WT cells in the 2–4 μM concentration range, at which cytotoxicity remained minimal (cell death < 5%; Figures S4J–S4K). In contrast, the ΔGAR195 mutant was resistant to PDS-mediated CSR suppression (Figure S4K). Thus, FBL promotes CSR by binding to G4 structures within GLTs.

### RNA 2′-*O*-methylation by Fibrillarin is essential for DNA recombination

Next, we assessed whether *Sα*-GLTs are actively 2′-*O*-methylated using reverse transcription (RT) at low dNTP concentrations followed by qPCR (RTL-qPCR; Figure 3H).^41^ This method exploits the principle that reverse transcriptase stalls at 2′-*O*-methylated ribose under low dNTP concentrations; thus, a lower “High/Low dNTP ratio” (the qPCR signal ratio of high to low dNTP conditions) reflects decreased 2′-*O*-methylation. Upon *Fbl* knockdown, High/Low dNTP ratios for 18S/28S rRNA and *Sα*-GLTs exhibited a consistent downward trend, whereas *B2m* transcripts remained unchanged (Figures 3I-3J).

To test whether catalytic activity is required for CSR, we generated FBL catalytic mutants targeting S-adenosyl methionine-binding motifs (PDB: 2IPX^36^ and 7SE7^42^; Figure S4L; Mut2, Mut11, and Mut14), as well as conserved catalytic residues identified through homology to *Escherichia coli* RrmJ^43,44^ (Mut15) (Figure 3K). All catalytic mutants (Mut2–Mut15) failed to support efficient CSR (Figures 3L-3M, and S4M). Although all variants, excluding Mut2, retained GLT-binding capacity (Figure S4N), their catalytic activity, as assessed by the High/Low dNTP ratio in RTL-qPCR for 28S rRNA and *Sα*-GLTs, was significantly reduced (Figure 3N). Together, these results demonstrate that FBL-mediated 2′-*O*-methylation of GLT transcripts is indispensable for CSR.

### MED19 cooperates with FBL to promote DNA synapsis and sustain snoRNA host gene expression

To identify cofactors acting alongside FBL during DNA synapsis, we focused on the C-terminal domain of AID, which is specifically required for *S*–*S* synapsis downstream of DNA cleavage.^16–19^ Although C-terminal truncation mutants (e.g., JP8Bdel, ΔC-AID) failed to undergo synapsis, fusion of the estrogen receptor ligand-binding domain to JP8Bdel (ΔC-AID-ER) partially restored CSR (Figures S5A–S5B). Given that nuclear receptors recruit the Mediator complex factors MED14 and MED19,^45,46^ and the physical interaction between FBL and MED19 is evolutionarily conserved,^47,48^ we evaluated MED14 and MED19. Targeted knockdown screening revealed that *Med19* depletion specifically impaired CSR (Figures S5C–S5F).

Both human and mouse MED19 feature N- and C-terminal intrinsically disordered regions containing proline-rich (PR) and lysine-rich (KR) domains, of which only KR domains are conserved from mammals to plants (Figures S5G–S5I). Knockdown-rescue assays revealed that deleting KR domains (ΔKNH, ΔKR1, and ΔKR2) substantially impaired CSR rescue, whereas deleting the PR domain (ΔPR) had no effect (Figures 4A-4C and S5J).

**Figure 4.**
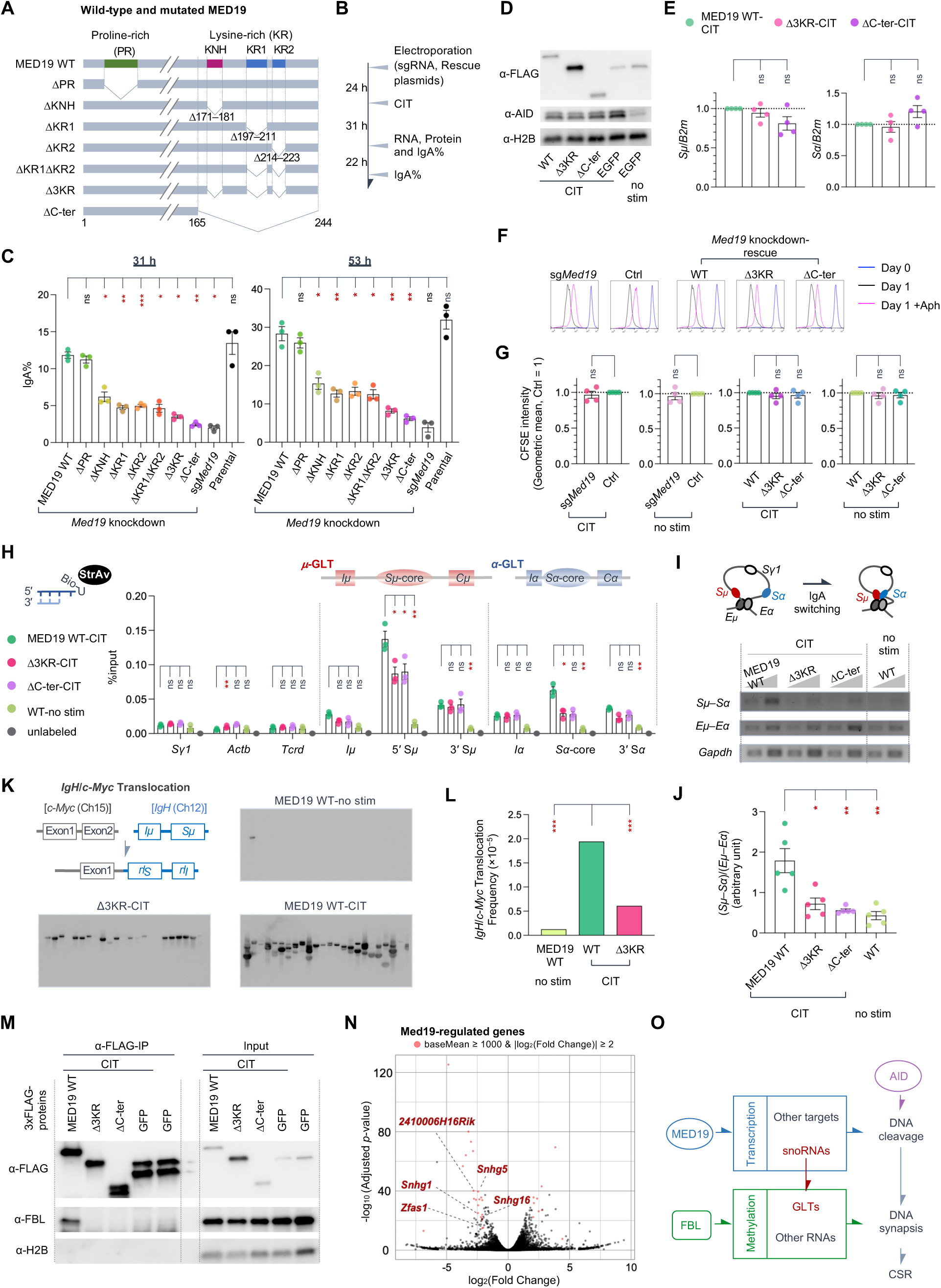
The Mediator subunit MED19 interacts with FBL to orchestrate CSR. (A–C) Structure of MED19 variants (A), knockdown-rescue timeline (B), and IgA switching efficiency (C) (*n* = 3). (D–G) Expression of AID protein (D), *Sμ*-(E, left), and *Sα*-(E, right) GLTs, CFSE histograms (F), and proliferation quantification (G). *n* = 4. (H) Biotin-dUTP DNA cleavage frequency in MED19-rescued cells (*n* = 3). (I and J) 3C assay of *Sμ*–*Sα* synapsis, presenting representative agarose gels (I) and interaction frequency (J) (*n* = 5). (K and L) Southern blotting (K) and quantification (L) of *IgH*/*c-Myc* translocation. See also Figure S5M. (M) Representative western blot of co-immunoprecipitation (Co-IP) presenting the interaction between MED19 and endogenous FBL. (N) Volcano plot of transcriptome-wide RNA-seq changes upon *Med19* knockdown. Red dots highlight highly expressed genes (baseMean ≥ 1000) that are significantly regulated (|log_2(_FC)| ≥ 2.0, *P*_adj <_ 0.05). Significantly regulated snoRNA host genes are labeled. See Table S3. (O) Proposed model for the role of MED19 in CSR. Data represent the mean ± SEM, except in (L), where values represent calculated frequencies. Dashed horizontal lines indicate normalized control levels (1.0). \**P*_adj <_ 0.05, \*\**P*_adj <_ 0.01, \*\*\**P*_adj <_ 0.001; ns, not significant. See STAR Methods for statistical details.

MED19 mutation did not alter AID/GLT expression and cell proliferation (Figures 4D-4G), likely because of the robust intrinsic apoptosis resistance in CH12 cells.^49^ Notably, unlike *Fbl* knockdown, MED19 mutants (Δ3KR and ΔC-ter) caused a 35%–50% reduction in *Sμ* and *Sα* DNA cleavage (Figure 4H), a greater than 50% loss of *Sμ*–*Sα* synapsis (Figures 4I-4J), and a drop in *IgH*/*c-Myc* translocation to ∼30% (Figures 4K, 4L, and S5K), suggesting that MED19 regulates multiple steps in CSR cascades.

Co-IP assays confirmed that the KR domains of MED19 mediate direct interactions with FBL (Figure 4M). Finally, transcriptome profiling further revealed that *Med19* knockdown led to the downregulation of several snoRNA host genes (*Snhg1*, *Snhg5*, *Snhg16, Zfas1,* and *2410006H16Rik*; Figures 4N, S5L, and S5M; Table S3; Data S6). Because host-derived snoRNAs direct FBL-mediated methylation (such as *snord1c* in α-GLT methylation),^26^ MED19 sustains FBL activity through both direct physical coupling and the regulation of snoRNA expression. Thus, MED19 serves a dual role in CSR, both supporting target DNA cleavage and cooperating with FBL to drive DNA synapsis (Figure 4O).

### Germline transcripts assemble into RNA condensates distinct from canonical nuclear bodies

Having established that FBL primarily targets pooled nuclear GLTs rather than nascent RNA/DNA hybrids, we performed RNA FISH to delineate the spatial organization of μ-, *γ1*-, and *α*-GLTs using transcript-specific probes (Figure 5A). *Sμ*-GLTs formed nuclear condensates of comparable size to *Neat1* paraspeckles and markedly larger than nascent *Gapdh* transcription loci (Figures 5B-5D). Thus, intronic *Sμ*-GLTs, generated from *μ*-GLTs via splicing, accumulate in distinct RNA condensates near the transcription loci. This finding conceptually extends a recent report identifying MS2-tagged GLT as “dynamic hubs”,^22^ demonstrating that endogenous GLTs intrinsically form nuclear RNA condensates within cells.

**Figure 5.**
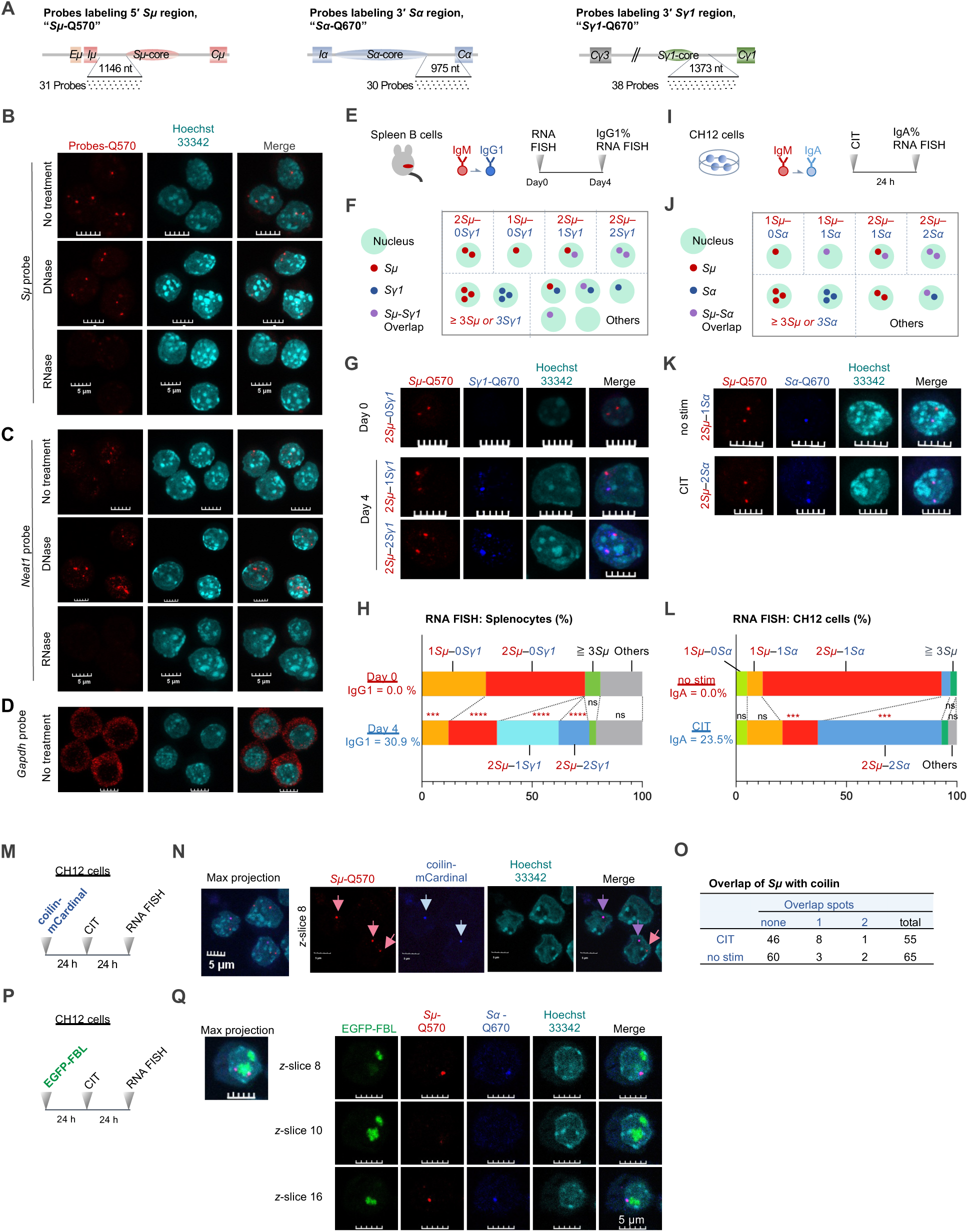
RNA FISH targeting *Sμ*-, *Sγ1*-, and *Sα*-GLTs reveals RNA condensate formation at *S* regions. (A) Target positions of *Sμ*-, *Sα*-, and *Sγ1*-GLT RNA FISH probes. (B–D) Specificity of *Sμ* (B), *Neat1* (C), and *Gapdh* (D) probes upon DNase or RNase treatment. (E–H) RNA FISH in primary spleen B cells: workflow (E), scheme (F), confocal images (G), and pattern frequencies (H) on days 0 and 4. (I–L) RNA FISH in CH12 cells: workflow (I), scheme (J), confocal images (K), and pattern frequencies (L) with or without (±) CIT stimulation. (M–O) Colocalization of *Sμ*-GLTs and mCardinal-coilin: workflow (M), confocal images (N), and quantification (O). See also Figures S6I–S6K. (P and Q) Spatial relationship between EGFP-FBL and GLT condensates: workflow (P) and confocal images (Q). See also Figures S6L–S6N. Scale bars, 5 μm. \*\*\**P*_adj <_ 0.001, **** *P*_adj <_ 0.0001; ns, not significant. See STAR Methods for statistical details.

In primary mouse B cells, stimulation with LPS and IL-4 induced distinct *Sμ*- and *Sγ1*-GLT foci (Figures 5E-5H). The signal distribution shifted from day 0 to day 4, reflecting constitutive *Sμ* expression and IL-4–induced *Sγ1* accumulation (*χ*^2^ = 108.2, *df* = 5, *P* < 0.0001; Figure 5H).

In CH12 cells, dual probing for *Sμ*- and *Sα*-GLTs highlighted a CIT-dependent distribution shift (*χ*^2^ = 100.5, *df* = 5, *P* < 0.0001; Figures 5I-5L). Notably, whereas CIT induced *Sα* expression at one allele, a second *Sα* focus was constitutively present prior to stimulation. Using *Iα* promoter knockdown, we established that CH12 cells harbor a constitutive, nonfunctional 3′ *Sα*-*Cα* allele that generates baseline transcripts without interfering with productive IgA switching (Figures S6A–S6F).

Because Cajal bodies harbor extranucleolar FBL,^50^ we tested the role of the Cajal body marker coilin in CSR. *Coil* knockdown did not impair CSR (Figures S6G–S6H). Moreover, costaining of *Sμ*-GLT transcripts with mCardinal-coilin revealed minimal overlap with *Sμ* condensates (Figures 5M-5O and S6I–S6K), distinguishing GLT condensates from Cajal bodies.

Notably, EGFP-FBL did not colocalize with GLT foci, and *Fbl* knockdown had no effect on *Sμ*- and *Sα*-GLT condensate formation (Figures 5P, 5Q, and S6L–S6N). Together, these findings indicate that FBL acts as a transient catalytic remodeler rather than a structural scaffold for GLT condensates.

### FBL actively drives the physical fusion of genomically distant GLT condensates

To profile GLT condensates under native conditions, we engineered *Sirius-8xMS2* knock-in CH12 cells at the *Sα*-core locus (Figures 6A, 6B, and S7A–S7K). Despite the minor background signal from EGFP-tagged tdMCP, tdMCP-bound *Sirius-8xMS2* loci colocalized reasonably with both *Sμ*- and *Sα*-GLT condensates (Figures 6C and S7L–S7O). Using this system, FLAG-tdMCP RIP-qPCR demonstrated that *Fbl* knockdown markedly reduced the recruitment of *Sμ*-GLT (prey) to *Sα*-GLT (bait) by > 65%, despite having a minimal effect (< 20%) on overall bait enrichment (Figure 6D). This provides direct biochemical evidence that FBL actively drives condensate fusion of genomically distant GLTs.

**Figure 6.**
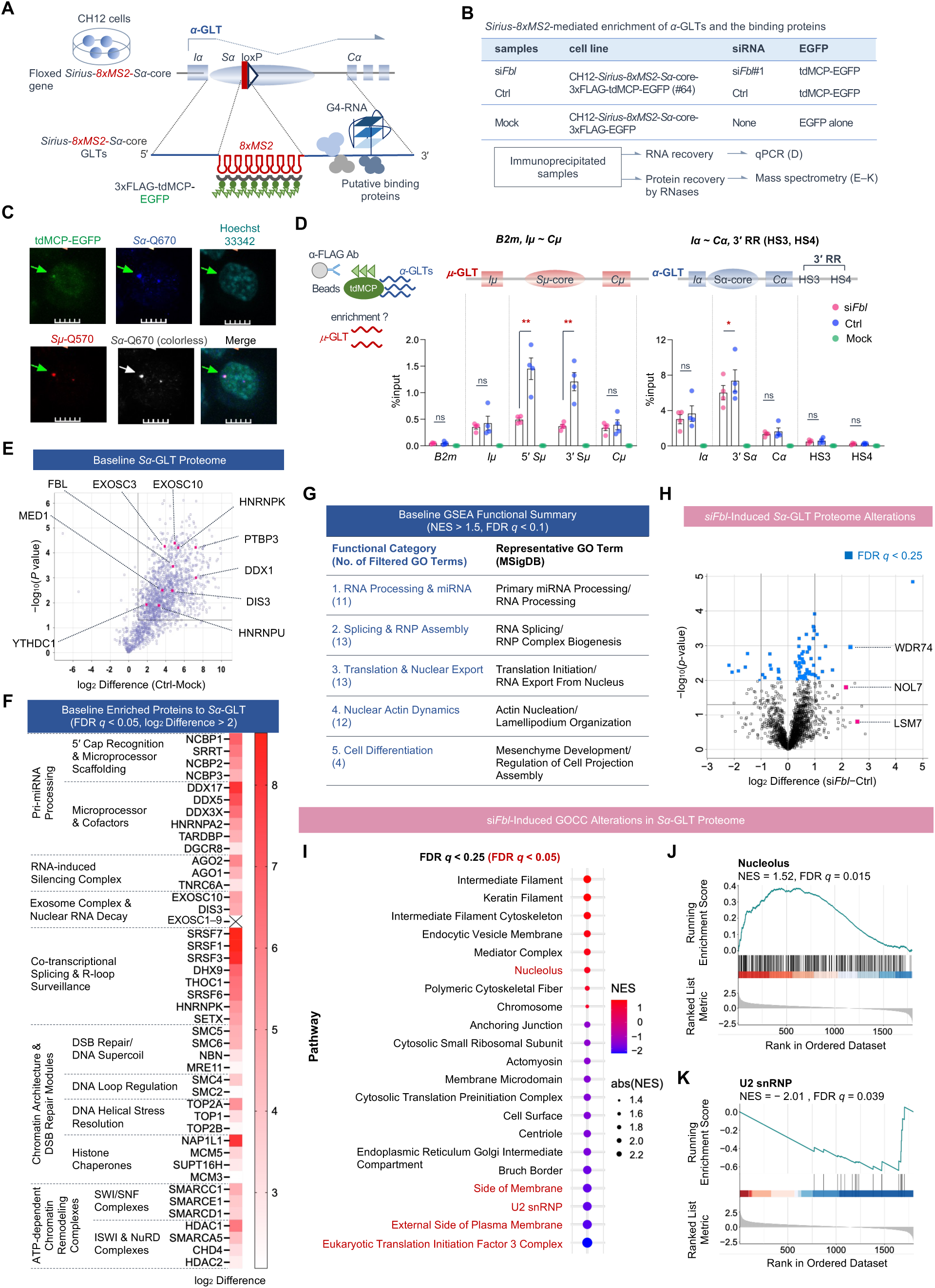
*Sirius-8xMS2*–mediated interactome of *Sα-*GLT identifies the proteome of a recombination focus regulated by FBL (A–C) *Sirius-8xMS2* system, detailing the knock-in design (A), RIP–MS workflow (B), and confocal images displaying the colocalization of tdMCP-EGFP with *Sμ*- and *Sα*-GLT RNA FISH signals (C). (D) RIP-qPCR evaluation of physical *Sμ*- and *Sα*-GLT interaction (fusion dynamics), demonstrating coprecipitation of *Sμ*-GLT upon capturing *Sα*-GLT complexes via 3xFLAG-tdMCP in *Sirius-8xMS2*-*Sα*-core cells (*n* = 4). (E–G) Baseline *Sα*-GLT proteome architecture defined by comparing control versus mock (EGFP alone). Volcano plot presenting baseline protein enrichment analyzed using Welch’s *t*-test, with previously reported GLT-binding proteins labeled (E; *n* = 3). Tabular summary of baseline enriched proteins categorized into RNA- and chromatin-related modules (F). Five baseline GOBP categories revealed by GSEA comparing control versus mock (G; *n* = 3). For (F), EXOSC1–9 components are presented as a summarized entry owing to space constraints. Full individual statistics for all EXOSC subunits (EXOSC1–EXOSC9) are listed in Data S8. For an exhaustive protein list and statistical metrics of GSEA analysis, see Data S9. RNP, ribonucleoprotein. (H–K) si*Fbl*-induced *Sα*-GLT proteome alterations defined by comparing si*Fbl* versus control. Volcano plot presenting differential protein abundance (log_2 D_ifference) between si*Fbl* and control, as analyzed using Welch’s *t*-test (H). Blue squares indicate proteins with FDR *q*-value < 0.25, with key nucleolar and RNA-processing factors labeled. See Data S10. GSEA dot plot of GO Cellular Component (GOCC) terms that were significantly enriched in (NES > 0, red) or evicted from *Sα*-GLT condensates (NES < 0, blue) following *Fbl* knockdown (I), with dot size indicating |NES|. GOCC terms with an FDR *q*-value < 0.05 are highlighted. See Data S11. GSEA enrichment plots of proteomic alterations upon *Fbl* knockdown for “Nucleolus” (J) and “U2 snRNP” (K). See also Figure S7P and S7Q. Scale bars, 5 μm. Data represent the mean ± SEM. \**P* < 0.05 and \*\**P* < 0.01; ns, not significant. Black lines in volcano plots indicate |log_2 D_ifference| = 1 and *P* = 0.05. See STAR Methods for statistical details.

### The baseline recombination focus harbors machinery for transcript processing and structural remodeling

To delineate the basal proteome of the *Sα*-GLT condensate, we performed mass spectrometry on RNase-eluted *Sα*-GLT complexes isolated from tdMCP-EGFP control versus Mock (GFP alone) cells (Figures 6E-6G; Data S7–S9). Welch’s *t*-test comparison of the eluted fractions revealed that previously reported GLT-interacting factors required for AID recruitment or CSR were, with few exceptions, successfully captured within this baseline proteome (Figure 6E).^23,26,51–59^

Proteomic profiling of these *Sα*-GLT condensates revealed a striking co-enrichment of discrete functional machinery involved in both RNA processing and chromatin dynamics (Figure 6F; Data S8). Specifically, these condensates were highly enriched for modules driving pri-miRNA processing,^60–64^ the RNA-induced silencing complex,^65,66^ nuclear exosome degradation,^67–69^ and R-loop surveillance,^70^ implicating active GLT cleavage and RNP remodeling. Concurrently, condensates selectively recruited proteins for chromatin-associated DNA damage response, DNA topology (supercoiling, helical stress, loop extrusion), and ATP-dependent chromatin remodeling,^66,71–79^ establishing these GLT condensates as functional centers for both DNA and RNA remodeling. Additionally, GSEA highlighted the enrichment of RNA- or chromatin-related pathways as well as terms associated with nuclear actin dynamics (supporting both polymerization and depolymerization) in these baseline *Sα*-GLT complexes (Figure 6G; Data S9). In contrast, NHEJ factors (Ku70/80, LIG4, 53BP1), APE1, and AID were not detected, and UNG was filtered out (log2 Difference ∼1, FDR q-value ∼ 0.1; Data S8), indicating that these condensates play a restricted role on DNA. As expected from the intrinsic physiological role of the *IgH* locus in antibody production, terms related to translation and nuclear export were also represented. Collectively, these results demonstrate that GLT condensates assemble into a dynamic and functional “recombination focus” equipped with the architectural and topological machinery required for 3D genomic reorganization during CSR.

### FBL suppression aberrantly enriches nucleolar proteins within the recombination focus

Proteomic profiling following *Fbl* knockdown revealed specific compositional shifts within *Sα*-GLT condensates (Figures 6H-6K, S7P, and S7Q; Datasets S10 and S11). A volcano plot based on Welch’s *t*-test demonstrated that a distinct set of proteins was significantly regulated by *Fbl* knockdown (Figure 6H; Data S10). Based on GSEA, *Fbl* knockdown enriched nucleolar and intermediate filament components (Figures 6I, 6J, and S7P), while depleting U2snRNP and eIF3 complexes from *Sα*-GLT condensates (Figures 6K and S7Q; Data S11). To pinpoint the key regulators of condensate fusion, we integrated the results of GSEA, Welch’s *t*-test, and the FBL interactome (Table S4; Data S12) and prioritized WDR74, LSM7, and NOL7.^80–82^ Notably, although WDR74 was undetectable (requiring imputation across all replicates) and LSM7 and NOL7 were detected only at near-background levels (requiring imputation in a subset of replicates) within baseline *Sα*-GLT condensates in control cells, *Fbl* knockdown induced their clear ectopic recruitment to these condensates.

Although WDR74 is canonically required for pre-60S ribosome biogenesis, other components of this machinery were not proportionally enriched alongside WDR74 (Figures S7R–S7S).^80,83–85^ This indicates that WDR74 is recruited to *Sα*-GLT condensates by a distinct mechanism from a generalized influx of the pre-60S biogenesis machinery. Functional validation using CRISPRi demonstrated that *Wdr74* knockdown (<10% residual expression) significantly enhanced CSR to IgA without affecting *Aicda* or GLT expression levels (Figures 7A-7E, 7I, S8A, and S8B). Conversely, knockdown of *Nol7* and *Lsm7* tended to reduce CSR, an effect caused by cell cycle delay through decreased rRNA biogenesis (Figures 7F-7I, S8A, and S8B).

**Figure 7.**
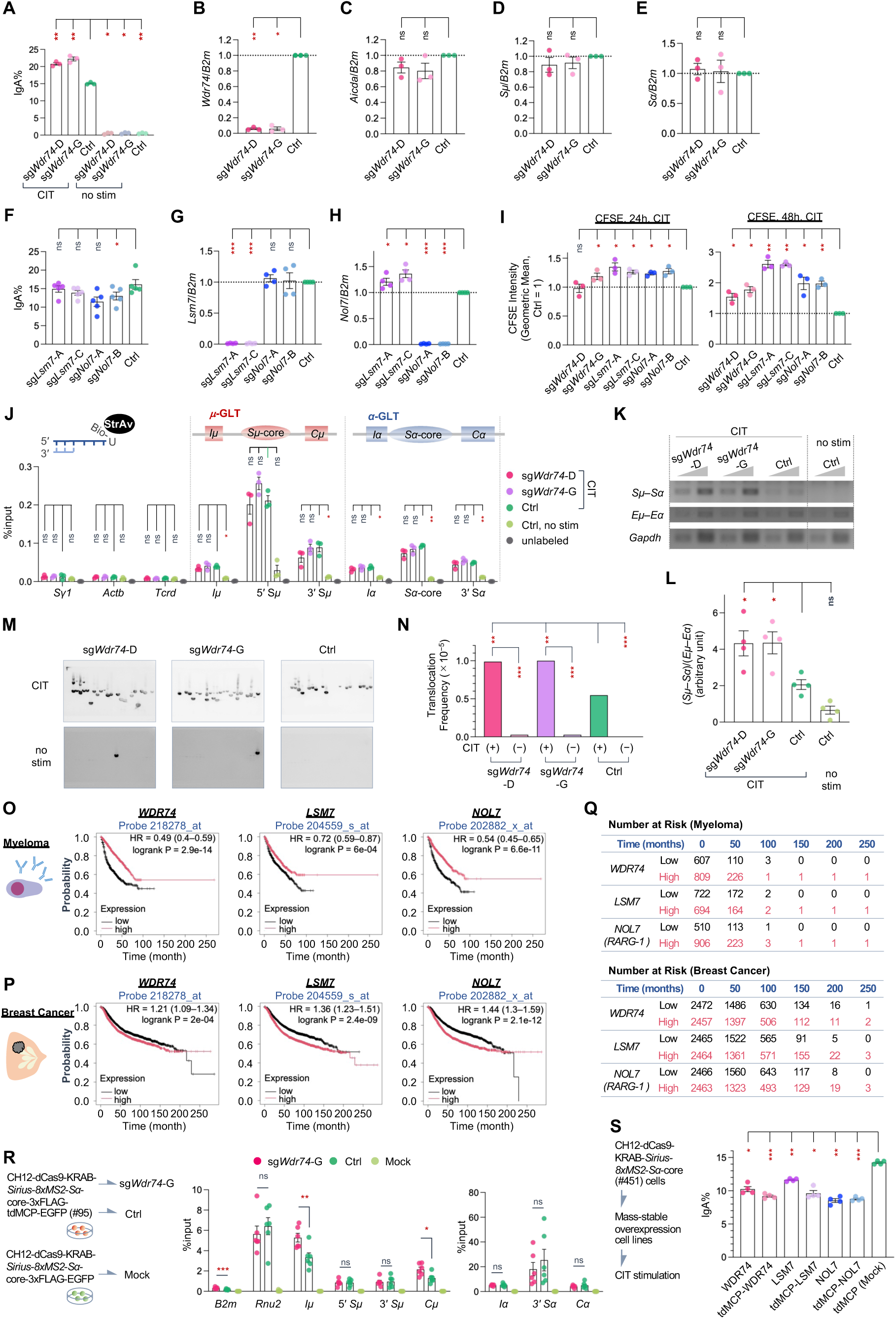
WDR74 depletion drives DNA synapsis and oncogenic translocation through condensate hyperfusion. (A–E) IgA switching efficiency (A) and RT-qPCR analysis of *Wdr74, Aicda, Sμ*-, and *Sα*-GLT transcript levels (B–E) upon *Wdr74* knockdown (*n* = 3). (F–H) IgA switching efficiency (F) and knockdown efficiencies (G and H) for *Lsm7 and Nol7* (*n* = 4–5). (I) Cell proliferation (CFSE) in *Wdr74-, Lsm7-, or Nol7-*knockdown cells (*n* = 3). See also Figures S8A and S8B. (J) DNA cleavage frequency upon *Wdr74* knockdown (*n* = 3). (K and L) 3C assay evaluating *Sμ*–*Sα* synapsis: agarose gels (K) and interaction frequency (L) upon *Wdr74* knockdown (*n* = 4). (M and N) Southern blot (M) and frequencies (N) of *IgH*/*c-Myc* translocation upon *Wdr74* knockdown. See also Figures S8C–F. (O–Q) Kaplan–Meier overall survival curves comparing high versus low *WDR74, LSM7, and NOL7* (*RARG-1*) expression in myeloma (O) and breast cancer (P), alongside number-at-risk summary tables (Q). (R) RIP-qPCR evaluation of physical *μ*–*α* GLT interaction, demonstrating coprecipitation of *Iμ*–*Cμ* GLT upon capturing *Sα*-GLT complexes via 3xFLAG–tdMCP in *Sirius-8xMS2*–knock-in cells (*n* = 6). In contrast to the si*Fbl* model in standard *Sirius-8xMS2* cells (Figure 6D), the dCas9-KRAB–expressing *Sirius-8xMS2* line exhibits a baseline predominance of spliced *μ*-GLT transcripts (*Iμ* and *Cμ*) over intronic transcripts (5′ *Sμ* and 3′ *Sμ*). (S) IgA switching efficiency upon overexpression of *WDR74*, *LSM7*, or *NOL7* with or without tdMCP fusion tags (*n* = 4). See also Figure S8G. Data represent the mean ± SEM except in (N), where values represent calculated frequencies. Dashed horizontal lines indicate normalized control levels (1.0). \**P*_adj <_ 0.05, \*\**P*_adj <_ 0.01, \*\*\**P*_adj <_ 0.001; ns, not significant. See STAR Methods for statistical details.

### WDR74 acts as a molecular brake on GLT condensate fusion to prevent oncogenic translocation

Systematic CSR evaluation revealed that *Wdr74* knockdown did not alter DNA cleavage (Figure 7J), but it significantly enhanced *Sμ*–*Sα* synapsis (Figures 7K–L) and *IgH*/*c-Myc* translocation (Figures 7M, 7N, and S8C–S8F). Thus, unlike MED19, WDR74 specifically functions as a negative regulator of DNA synapsis without affecting DNA cleavage.

Clinically, *IgH* translocation represents a well-established high-risk cytogenetic abnormality in B-cell lymphoma and multiple myeloma^86,87^. While a previous small-cohort study in diffuse large B-cell lymphoma (DLBCL)^88^ suggested a potential link between low WDR74 expression and poor survival, our analysis of a large patient cohort with myeloma (*n* = 1416, Kaplan–Meier Plotter;^89^ Figures 7O and 7Q) robustly demonstrated that low expression of *WDR74, LSM7,* or *NOL7* predicted significantly shorter overall survival. This pattern was strikingly inverted in solid tumors such as breast cancer (*n* = 4929; Figures 7P-7Q), in which their elevated expression was correlated with poor outcomes, likely driven by increased cancer cell proliferation.

Mechanistically, RIP-qPCR confirmed that *Wdr74* knockdown markedly enhanced *μ*-GLT recruitment to *Sα-*GLT condensates (Figure 7R). Conversely, the overexpression of WDR74, LSM7, and NOL7 (with or without tdMCP fusion tags) suppressed CSR (Figures 7S and S8G), demonstrating that excess nucleoplasmic accumulation of these nucleolar factors directly restrains CSR. Together, these results indicate that the loss of nucleolar factors, such as WDR74, relieves fusion constraints on RNA condensates within the recombination focus, thereby promoting oncogenesis.

Collectively, these findings establish a novel mechanistic cascade in which FBL-dependent ncRNA 2′-*O*-methylation drives WDR74 eviction, facilitating *Sμ–Sα* condensate fusion and productive CSR (Figure S8H).

## DISCUSSION

In this study, we elucidated that 2′-*O*-methylation is a major physicochemical determinant that dictates the functional properties and fusion plasticity of noncoding RNA condensates. Specifically, FBL-mediated 2′-*O*-methylation actively maintains the fusion capability of RNA condensates by preventing the aberrant binding of specific nucleolar proteins, such as WDR74, which ultimately drives *S*–*S* synapsis during class switch recombination (CSR). These findings establish FBL as a pivotal epitranscriptomic regulator controlling the dynamics of the “recombination focus” scaffolded by *Sα*-GLT acting as an architectural RNA.^90^

Importantly, our *Sirius-8xMS2*–mediated RIP-qPCR allows us to directly capture genuine RNA condensate synapsis between noncoding *μ*- and *α*-GLTs prior to DNA recombination, rather than chimeric transcripts generated from post-CSR recombined DNA. Because the 3′ *Sμ*–*Cμ* region downstream of the *Sμ*-core is deleted upon CSR, any decrease or increase in 3′ *Sμ*–*Cμ* RIP-qPCR signals directly reflects the reduction or enhancement of physical interactions involving intact, full-length *μ*- and *α*-GLTs. This technical precision demonstrates that FBL-mediated 2′-*O*-methylation directly dictates the physical fusion of *μ*- and *α*-GLT condensates before CSR. Crucially, because an RNA-specific modification directly governs genomic interactions, RNA condensate synapsis cannot be a mere byproduct of DNA recombination; rather, it acts upstream to instruct genomic synapsis.

The most striking observation in the *Sα*-GLT proteome analysis upon *Fbl* knockdown was the enrichment of nucleolar proteins. Insufficient 2′-O-methylation provokes an aberrant accumulation of nucleolar factors, most notably WDR74, leading to hypo-fusion of RNA condensates. Physiologically, a sufficient level of 2′-O-methylation by FBL strictly excludes the ectopic infiltration of WDR74 into GLT condensates and retains condensate plasticity for normal fusion, thereby enabling efficient S–S synapsis for CSR while carrying an inherent risk of oncogenic translocation. In contrast, *Wdr74* knockdown leads to hyper-fusion of the condensates and enhanced CSR, S–S synapsis, and *IgH/c-Myc* translocation. These results unveiled the mechanistic basis for the clinical paradox in which low WDR74 expression is correlated with poor prognosis in B-cell malignancies, despite conferring reduced cancer cell proliferation in solid tumors.^88^ Because it promotes additional translocations, low WDR74 expression may serve as a functional biomarker in B-cell tumors for aggressive phenotypes requiring intensified therapy.

Baseline *Sα*-GLT condensates, maintained under sufficient FBL-mediated 2′-*O*-methylation, are not passive scaffolds; rather, they compartmentalize the dynamic machinery orchestrating the remodeling of GLTs, RNPs, R-loops, and chromatin architecture. Comparison with the recently established proteomic landscape of human nuclear condensates^91^ underscores the unique identity of the *Sα*-GLT condensate. Whereas canonical nuclear bodies are typically classified into discrete functional compartments, such as nuclear speckles (RNA splicing), promyelocytic leukemia (PML) bodies (chromatin organization), or transcriptional condensates, the baseline *Sα*-GLT interactome uniquely exhibits a hybrid proteomic signature integrating all three modules alongside actin dynamics. This composite architecture demonstrates that the *Sα*-GLT–scaffolded recombination focus is a highly specialized, multifunctional nuclear condensate designed to orchestrate CSR.

Crucially, canonical NHEJ machinery (e.g., Ku70/80, LIG4, 53BP1) was not detected in this baseline proteome, underscoring that GLT condensates regulate DNA and RNA topology rather than forming NHEJ repair foci. Instead, the baseline interactome selectively captures dynamic regulatory components, including the nuclear exosome (EXOSC1–10, DIS3) associated with R-loop resolution and AID recruitment,^67–69^ alongside factors modulating RNA dynamics and chromatin topology (e.g., HNRNPK, TOP1, SUPT16H).^51,52,75,77,78^ This selective enrichment suggests a dual role for GLT condensates as architectural scaffolds coordinating DNA cleavage and synapsis. First, by fine-tuning local chromatin accessibility and RNA–DNA hybrid dynamics, these condensates may establish a permissive 3D conformation of DNA and RNA for AID-dependent DNA cleavage. Second, they safeguard cleaved DNA ends from premature local religation by shielding them from NHEJ machinery until they encounter partner *S*-region loci for productive recombination.

Rather than WDR74 alone, multiple nucleolar proteins likely contribute to this loss of fusion plasticity in RNA condensates. Biologically, the nucleolus is a multiphasic nuclear condensate consisting of three distinct layers (fibrillar center, dense fibrillar component, and granular component)^92^ that do not readily coalesce, imparting a rigid organization. Thus, the phenotypic shift of *Sα*-GLT condensates from a fusion-ready to a fusion-resistant state upon *Fbl* knockdown indicates that 2′-*O*-methylation is essential for maintaining the fusion plasticity required for CSR.

Conversely, a distinct subset of proteins, including U2 snRNP components, is evicted from *Sα*-GLT condensates upon *Fbl* knockdown. Although validating U2 snRNP overexpressions (e.g., SF3B1, SF3B3, and SF3B5),^93,94^ is technically challenging, its depletion impairs CSR.^95^ This positive link between U2 snRNP enrichment and condensate fusion is consistent with the established role of U2 snRNP in mediating the internalization of paraspeckles into nuclear speckles.^96^ Eviction of U2 snRNP raises the intriguing possibility that, as RNU2 snRNA itself is targeted by FBL-dependent 2′-*O*-methylation,^97^ insufficient modification induces conformational alterations that diminish functional U2 snRNP pools. Thus, FBL-mediated 2′-*O*-methylation on non-GLT RNAs may cooperatively support *Sα*-GLT condensate assembly and plasticity.

Notably, the recruitment of nuclear actin dynamics factors in *Sα*-GLT condensates highlights the pivotal role of the actin machinery^98,99^ and redefines the recombination focus as a structural scaffold supporting DNA recombination. Nuclear actin filaments facilitate chromatin accessibility^100^ and mediate damaged heterochromatin relocalization during repair.^101,102^ The enrichment of actin regulators suggests that the nuclear actin dynamics modulate chromatin accessibility and condensate plasticity within the recombination focus.

Beyond FBL itself, we identified MED19 as an upstream transcriptional regulator controlling snoRNA host gene expression, thereby sustaining FBL-dependent 2′-*O*-methylation. The functional divergence between MED14 and MED19 demonstrated that CSR impairment upon MED19 depletion reflects a specialized regulatory role rather than nonspecific Mediator collapse. MED19 acts as a transcriptional hub coupling RNA polymerase II to snoRNA host genes, representing a discrete branch tuning 2′-*O*-methylation and physical plasticity at *Sα*-GLT condensates.

Ultimately, our study uncovers a fundamental epitranscriptomic mechanism in which FBL-mediated 2′-*O*-methylation of *Sα*-GLT maintains ncRNA condensates in a physically plastic, fusion-ready state by preventing the ectopic infiltration of nucleolar factors. By preserving condensate plasticity and orchestrating DNA synapsis, this RNA modification enables the functional assembly of the recombination focus, providing a working model (Figure S8H) for the spatiotemporal regulation of class switch recombination.

### Limitations of the study

Although our study established a biophysical model in which FBL-mediated RNA 2′-*O*-methylation shapes GLT condensates for *S*–*S* synapsis, several limitations remain. First, direct live-cell biophysical fluidity measurements (e.g., FRAP) were technically constrained by nucleoplasmic tdMCP-EGFP background signals. Second, mapping target 2′-*O*-methylated residues at single-base resolution on GLTs awaits nanopore sequencing or direct chemical profiling. Third, dissecting the precise contribution of the nuclear actin remodeling without cytotoxic effects requires future locus-specific intervention systems. Finally, clarifying whether translocations directly drive poorer prognosis in low-WDR74 B-cell malignancies will require large-cohort genomic analysis and genetically engineered mouse models.

## RESOURCE AVAILABILITY

### Lead contact

Requests for further information and resources should be directed to and fulfilled by the lead contact, Maki Kobayashi.

### Materials availability

All unique/stable reagents generated in this study are available from the lead contact with a completed materials transfer agreement; however, we may require a payment and/or a completed materials transfer agreement if there is potential for commercial application.

### Data and code availability

Mass spectrometry (proteomics), RNA-sequencing datasets, and original R scripts generated in this study have been deposited in public repositories as listed below. All datasets and associated source data will be made publicly accessible as of the date of publication:

1. LexA-mediated *Sα*-core iChIP proteomic datasets (Set 1 and Set 2) were deposited in jPOSTrepo^103^ / ProteomeXchange under accession numbers PXD085038 and PXD085041, respectively.
2. The Med19-dependent transcriptome (RNA-seq) dataset has been deposited in the NCBI Gene Expression Omnibus (GEO) under accession number GSE349295.
3. The Sirius 8xMS2-mediated *Sα*-GLT interactome proteomic dataset has been deposited in jPOSTrepo/ProteomeXchange under accession number PXD083456.
4. The FBL interactome proteomic dataset has been deposited in jPOSTrepo/ProteomeXchange under accession number PXD083413.
5. All original R scripts for statistical analyses and custom data visualization, along with the corresponding input source data files required to run these scripts, have been deposited on GitHub and are publicly accessible (https://github.com/omurakanon/R_scripts_for_M.Kobayashi_etal_2026).

Any additional information required to reanalyze the data reported in this study is available from the lead contact upon reasonable request.

## ACKNOWLEDGMENTS

We thank the members of the Department of Immunology and Genomic Medicine and the Center for Cancer Immunotherapy and Immunobiology (CCII) at the Graduate School of Medicine, Kyoto University, for their continuous support. We are particularly grateful to Misao Takemoto, Eiko Shimizu, Maki Sasanuma, and Kyoko Hirano for their excellent technical assistance. We also thank Dr. Mayu Terakawa at the Kyoto University Office of Research Acceleration for insightful suggestions regarding manuscript figures, as well as Prof. Toshihiko Ogura and Dr. Tomohiro Yamazaki (Osaka University) for generous advice on proteomic analysis.

In addition, we express our gratitude to Evident Corporation (Tokyo, Japan) for providing access to the FLUOVIEW FV4000 confocal laser scanning microscope, and to the Kyoto University Live Imaging Center for facility support. Mass spectrometry analysis was supported by the Joint Usage and Joint Research Programs of the Institute of Advanced Medical Sciences, Tokushima University.

This work was supported by JSPS KAKENHI Grant Numbers JP26K09317 (to M.K.) and JP25H01028 (to M.K. and T.H.). This research was conducted at CCII, which is supported by the Yanai Fund. The authors thank Enago for English language editing.

## AUTHOR CONTRIBUTIONS

**Conceptualization**, M.K. and T.M.; **Methodology**, M.K., J.X., A.H., T.F., H.F., T.O., and H.K.; **Software**, K.O. and K.I.; **Validation**, M.K.; **Formal Analysis**, M.K., K.O., K.I., and H.K.; **Investigation**, M.K., J.X., A.H., T.M., Z.Y., X.C., W.W., T.F., H.F., L.L.U., T.O., and H.K.; **Resources**, M.K., J.X., A.H., T.F., H.F., and T.H.; **Data Curation**, M.K., K.O., and K.I.; **Writing – Original Draft**, M.K., K.O., K.I., and H.K.; **Writing – Review & Editing**, M.K., K.I., and T.H.; **Visualization**, M.K., K.O., and K.I.; **Supervision**, M.K. and T.H.; **Project Administration**, M.K.; **Funding Acquisition**, M.K. and T.H.

## DECLARATION OF GENERATIVE AI AND AI-ASSISTED TECHNOLOGIES IN THE WRITING PROCESS

During the preparation of this manuscript, the authors used Gemini (Google) and Paperpal (Cactus Communications) to obtain suggestions on grammar, spelling, and style. After using these tools, the authors reviewed and edited the content as needed and took full responsibility for the content of the publication.

## SUPPLEMENTAL INFORMATION

### [Supplemental Information]

Supplemental_Information.pdf

(Figures S1–S8, and Tables S1–S4)

The following supplemental materials referenced in the text will be made available upon formal publication (or upon reasonable request):

### [Supplementary Datasets]

Data_S1_Oligonucleotides_Used_in_This_Study.xlsx

Data_S2_LexA_SaCore_iChIP_Proteome_Set1-Set2-Combined_Related-to-Fig1C.xlsx

Data_S3_LexA_SaCore_iChIP_Proteome_enrichGO_Related-to-Fig1D-1E.xlsx

Data_S4_LexA_SaCore_iChIP_Proteome_GSEA_Related-to-Fig1F-1G.xlsx

Data_S5_FBL_Junction-Seq_Related-to-FigS3H–I.xlsx

Data_S6_Med19-DEG_Related-to-Fig4N.xlsx

Data_S7_MS2_SaCore_RIP_Proteome_Unprocessed_MS-signals_Related-to-Fig6.xlsx

Data_S8_MS2_SaCore_RIP_Proteome_Baseline_t-test_Related-to-Fig6E–F.xlsx

Data_S9_MS2_SaCore_RIP_Proteome_Baseline_GSEA_Related-to-Fig6G.xlsx

Data_S10_MS2_SaCore_RIP_Proteome_siFbl-induced_t-test_Related-to-Fig6H.xlsx

Data_S11_MS2_SaCore_RIP_Proteome_siFbl-induced_GSEA_Related-to-Fig6I-6K.xlsx

Data_S12_FBL-Interactome_Related-to-Fig6H.xlsx

### [Source Data]

Source_Data_1_Gels_and_Blots.pdf

Source_Data_2_3C assay.pdf

Source_Data_Numerical_Figure1-S1-S2.xlsx

Source_Data_Numerical_Figure2-S3.xlsx

Source_Data_Numerical_Figure3-S4.xlsx

Source_Data_Numerical_Figure4-S5.xlsx

Source_Data_Numerical_Figure5-S6.xlsx

Source_Data_Numerical_Figure6-S7.xlsx

Source_Data_Numerical_Figure7-S8.xlsx

## STAR★METHODS

### KEY RESOURCES TABLE

| REAGENT OR RESOURCE | SOURCE | IDENTIFIER |
|---|---|---|
| Antibodies |  |  |
| PE-labeled goat anti-mouse IgA | SouthernBiotec | RRID: AB_2794375; Cat# 1040-09 |
| BD Pharmingen Biotin Rat Anti-Mouse IgG3, clone R40-82 | BD Biosciences | RRID: AB_394838.Cat# 553401 |
| Ultra-LEAF Purified anti-mouse CD16/32 Clone93 | BioLegend | RRID: AB_2561482; Cat# 101330 |
| Anti-β-Actin (ACTB) Antibody | Merck | RRID: AB_476692; Cat# A1978 |
| AID Monoclonal Antibody (mAID-2), eBioscience | Thermo Fisher Scientific | RRID: AB_10698016; Cat# 14-5959-80 |
| Anti-DNA/RNA hybrid S9.6 antibody | Merck | RRID: AB_2861387; Cat# MAB1095 |
| ANTI-FLAG M2 antibody, Mouse monoclonal, glycerol solution | Sigma-Aldrich | RRID: AB_262044; Cat# F1804 |
| ANTI-FLAG M2 antibody, Mouse monoclonal, glycerol-free | Sigma-Aldrich | RRID: AB_2910145; Cat# B3111 |
| Anti-Fibrillarin antibody | Abcam | RRID: AB_2936302; Cat# ab226178 |
| Anti-Glyceraldehyde-3-Phosphate Dehydrogenase Antibody, clone 6C5 | Merck | RRID: AB_2107445; Cat# MAB374 |
| Rb pAb to Histone H2B | Abcam | RRID: AB_302612; Cat# ab1790 |
| Anti-Histone H3 antibody-Nuclear Marker and ChIP Grade | Abcam | RRID: AB_302613; Cat# ab1791 |
| hnRNP U Antibody (3G6) | Santa Cruz | RRID: AB_627741; Cat# sc-32315 |
| Myc antibody (9E10) | Santa Cruz | RRID: AB_627268; Cat# sc-40 |
| C23 (Nucleolin) antibody (MS-3) | Santa Cruz | RRID: AB_670271; Cat# sc-8031 |
| B23 (Nucleophosmin) antibody (FC-8791) | Santa Cruz | RRID: AB_628035; Cat# sc-32256 |
| Anti-α-Tubulin Mouse mAb (DM1A) | Merck | RRID: AB_2617116; Cat# CP06 |
| Bacterial and virus strains |  |  |
| XL10-Gold *E. coli* | Honjo Lab Stock | N/A |
| Biological samples |  |  |
| Primary mouse splenic B cells | This study | N/A |
| Chemicals, peptides, and recombinant proteins |  |  |
| 4-Hydroxytamoxifen (4-OHT) | Sigma-Aldrich | Cat# H6578-10MG |
| AluI | Takara Bio | Cat# 1004B |
| AseI | New England Biolabs (NEB) | Cat# R0526S |
| Benzonase Nuclease | Merck | Cat# 70746-3CN |
| Biotin-16-dUTP | Roche Diagnostics | Cat# 11093070910 |
| Blasticidin | InvivoGen | Cat# ant-bl-1 |
| cOmplete, EDTA-free, Protease Inhibitor Cocktail | Merck | Cat# 11873580001 |
| DNase I, grade II, from bovine pancreas | Merck | Cat# 10104159001 |
| HaeIII (High Concentration) | Takara Bio | Cat# 1051BH |
| HindIII (High Concentration) | Takara Bio | Cat# 1060BH |
| Interleukin-4, Mouse, recombinant, Animal-derived-free | Fujifilm Wako Pure Chemical | Cat# 090-07601 |
| KOD FX Neo | TOYOBO | Cat# 07097-34 |
| Ligation high | TOYOBO | Cat# LGK-101 |
| Lipofectamine 3000 Transfection Reagent | Thermo Fisher Scientific | Cat# L3000001 |
| Maxima H Minus Reverse Transcriptase | Thermo Fisher Scientific | Cat# EP0752 |
| Medium NCTC-109 (1X) | Thermo Fisher Scientific | Cat# 21340039 |
| Platinum Taq DNA Polymerase | Thermo Fisher Scientific | Cat# 10966026 |
| Platinum II Taq Hot-Start DNA Polymerase | Thermo Fisher Scientific | Cat# 14966005 |
| PowerUp SYBR Green Master Mix | Thermo Fisher Scientific | Cat# A25742 |
| PrimeSTAR GXL DNA Polymerase | Takara Bio | Cat# R050A |
| ProLong Glass Antifade Mountant | Thermo Fisher Scientific | Cat# P36982 |
| Puromycin | Nacalai Tesque | Cat# 19752-22 |
| Pyridostatin hydrochloride (PDS) | Sigma-Aldrich | Cat# SML2690 |
| Q5 High-Fidelity 2X Master Mix | New England Biolabs (NEB) | Cat# M0492S |
| Recombinant Human TGFβ1 Protein | R&D Systems | Cat# 240-B-010/CF |
| Ribonuclease (DNase free) Glycerol Solution (RNase A) | Nippon Gene | Cat# 312-01931 |
| RNase H | New England Biolabs (NEB) | Cat# M0297L |
| RNaseOUT Recombinant Ribonuclease Inhibitor | Thermo Fisher Scientific | Cat# 10777019 |
| RNase T1 | Thermo Fisher Scientific | Cat# EN0541 |
| Sepazol-RNA II Super | Nacalai Tesque | Cat# 30487-46 |
| ShortCut RNase III | New England Biolabs (NEB) | Cat# M0245S |
| Streptavidin-PE | BioLegend | Cat# 405203 |
| SuperScript IV Reverse Transcriptase | Thermo Fisher Scientific | Cat# 18090050 |
| SuperScript IV VILO Master Mix with ezDNase | Thermo Fisher Scientific | Cat# 11766050 |
| T4 DNA Ligase | Takara Bio | Cat# 2011B |
| T4 Polynucleotide Kinase | New England Biolabs (NEB) | Cat# M0201L |
| Terminal Deoxynucleotidyl Transferase | Takara Bio | Cat# 2230B |
| TRIzol Reagent | Thermo Fisher Scientific | Cat# 15596026 |
| TURBO DNase | Thermo Fisher Scientific | Cat# AM2238 |
| Critical commercial assays |  |  |
| CellTrace CFSE Cell Proliferation Kit | Thermo Fisher Scientific | Cat# C34554 |
| Chemi-Lumi One L | Nacalai Tesque | Cat# 07880-70 |
| Chemi-Lumi One Super | Nacalai Tesque | Cat# 02230-30 |
| DNA Damage Detection Kit - γH2AX - Deep Red | Dojindo | Cat# G265 |
| ExoSAP-IT *Express* PCR Product Cleanup | Thermo Fisher Scientific | Cat# 75001.200.UL |
| FuGENE 6 | Promega | Cat# E2691 |
| GeneArt Gibson Assembly HiFi Master Mix | Thermo Fisher Scientific | Cat# A46627 |
| GeneArt Gibson Assembly HiFi Cloning Kit | Thermo Fisher Scientific | Cat# A46624 |
| GeneArt Seamless Cloning Enzyme Mix | Thermo Fisher Scientific | Cat# A14606 |
| IDT for Illumina - DNA/RNA UD Indexes, Tagmentation | Illumina | Cat# 20090651, 20090656, 20091658, or 20091660 |
| KOD-Plus-Mutagenesis Kit | TOYOBO | Cat# SMK-101 |
| Maxwell RSC simplyRNA Cells Kit | Promega | Cat# AS1390 |
| MojoSort Mouse Dead Cell Removal Kit | BioLegend | Cat# 480157 |
| MojoSort Mouse Pan B Cell Isolation Kit | BioLegend | Cat# 480051 |
| Nextera XT DNA Library Prep Kit | Illumina | Cat# FC-131-1024 |
| NovaSeq 6000 S4 Reagent Kit v1.5 | Illumina | Cat# 20028312 |
| NovaSeq Xp 4-Lane Kit v1.5 | Illumina | Cat# 20043131 |
| pBluescript II KS (+) Phagemid Kit | Agilent | Cat# 212207 |
| pGEM-T Easy | Promega | Cat# A1360 |
| RNA Clean & Concentrator-5 | Zymo Research | Cat# R1014 |
| RNA Clean & Concentrator-25 | Zymo Research | Cat# R1017 |
| SF Cell Line 96-well Nucleofector Kit | Lonza | Cat# V4SC-2096 |
| SMART-Seq v4 Ultra Low Input RNA Kit for Sequencing | Takara Bio | Cat# Z4889N |
| TaKaRa Universal Buffer, 10× M | Takara Bio | Cat# N/A |
| Trypsin/Lys-C Mix, Mass Spec Grade | Promega | Cat# V5071 |
| truChIP Chromatin Shearing Kit with Formaldehyde | Covaris | Cat# 520154 |
| ViaFect Transfection Reagent | Promega | Cat# E4981 |
| Zero Blunt™ TOPO™ PCR Cloning Kit, without competent cells | Thermo Fisher Scientific | Cat# 451245 |
| Deposited data |  |  |
| FBL interactome | jPOST/ProteomeXchange | jPOST: PXD083413 |
| LexA-mediated *Sα*-core iChIP proteomic datasets | jPOST/ProteomeXchange | jPOST: Set1, PXD085038；Set 2, PXD085041 |
| Med19-dependent transcriptome (RNA-seq) data | NCBI Gene Expression Omnibus (GEO) | GEO: GSE349295 |
| Sirius 8xMS2-mediated Sα-GLT RIP interactome | jPOST/ProteomeXchange | jPOST: PXD083456 |
| Experimental models: Cell lines |  |  |
| CH12 (CH12F3-2A) wild-type (WT) | Honjo Laboratory stock | RRID: CVCL_E068; Cat# N/A |
| CH12 (CH12F3-2A)-Bcl2-Neo | Honjo Laboratory stock | RRID: N/A;Cat# N/A |
| CH12 (CH12F3-2A)-Bcl2-BSD | This study | RRID: N/A;Cat# N/A |
| CH12 (CH12F3-2A)-Bcl2-Neo-FBL-WT | This study | RRID: N/A;Cat# N/A |
| CH12 (CH12F3-2A)-Bcl2-Neo-FBL-ΔGAR | This study | RRID: N/A;Cat# N/A |
| CH12 (CH12F3-2A)-Bcl2-Neo-FBL-91D3 | This study | RRID: N/A;Cat# N/A |
| CH12 (CH12F3-2A)-Bcl2-Neo-FBL-195D5 | This study | RRID: N/A;Cat# N/A |
| CH12 (CH12F3-2A)-Bcl2-Neo-FBL-ΔGAR91 | This study | RRID: N/A;Cat# N/A |
| CH12 (CH12F3-2A)-Bcl2-Neo-FBL-ΔGAR195 | This study | RRID: N/A;Cat# N/A |
| CH12 (CH12F3-2A)-Bcl2-Neo-FBL-WT | This study | RRID: N/A;Cat# N/A |
| CH12 (CH12F3-2A)-Bcl2-Neo-FBL-ΔGAA167-169 (Mut2) | This study | RRID: N/A;Cat# N/A |
| CH12 (CH12F3-2A)-Bcl2-Neo-FBL-EDAR215-218AAAA/DV236-237AA (Mut11) | This study | RRID: N/A;Cat# N/A |
| CH12 (CH12F3-2A)-Bcl2-Neo-FBL-ΔVEF190-192 (Mut14) | This study | RRID: N/A; Cat# N/A |
| CH12 (CH12F3-2A)-Bcl2-Neo-FBL-K143A/K265A (Mut15) | This study | RRID: N/A; Cat# N/A |
| CH12 (CH12F3-2A)-AID-Knockout (AID-KO) | Al Ismail et al., 2017 | RRID: N/A; Cat# N/A |
| CH12-dCas9-KRAB (#40) | Kobayashi et al., 2023 | RRID: N/A; Cat# N/A |
| CH12-FNLDD-8xLexA-*Sα*-core | This study | RRID: N/A; Cat# N/A |
| CH12-FNLDD-alone | Husain et al., 2020 | RRID: N/A; Cat# N/A |
| CH12-AID-KO-mAIDER-IRES-Puro | This study | RRID: N/A; Cat# N/A |
| CH12-AID-KO-ΔAID-ER-IRES-Puro (CH12-JP8Bdel-ER-IRES-Puro) | This study | RRID: N/A; Cat# N/A |
| CH12-*Sirius-8xMS2-Sα*-core (#11) | This study | RRID: N/A; Cat# N/A |
| CH12-*Sirius-8xMS2-Sα*-core-3xFLAG-tdMCP-EGFP (#64) | This study | RRID: N/A; Cat# N/A |
| CH12-*Sirius-8xMS2-Sα*-core-3xFLAG-EGFP | This study | RRID: N/A; Cat# N/A |
| CH12-dCas9-KRAB-*Sirius-8xMS2-Sα*-core (#451) | This study | RRID: N/A; Cat# N/A |
| CH12-dCas9-KRAB-*Sirius-8xMS2-Sα*-core-3xFLAG-tdMCP-EGFP (#95) | This study | RRID: N/A; Cat# N/A |
| CH12-dCas9-KRAB-*Sirius-8xMS2-Sα*-core-3xFLAG-EGFP | This study | RRID: N/A; Cat# N/A |
| H1299dA3-1 | Ogiwara et al. 2011 | RRID: CVCL_0060; Cat# N/A |
| HEK293T | Honjo Laboratory stock | RRID: CVCL_0063; Cat# N/A |
| HeLa | Honjo Laboratory stock | RRID: CVCL_0030; Cat# N/A |
| Experimental models: Organisms/strains |  |  |
| Mouse: C57BL/6NCrSlc | Japan SLC | N/A |
| Oligonucleotides |  |  |
| Primers for PCR, RT-qPCR, and sequencing and oligonucleotides | This study | See Data S1 |
| sgRNA sequences for CRISPRi / CRISPR-Cas9 | This study | See Data S1 |
| si*Fbl*#1 | Thermo Fisher Scientific | Cat# s65880 |
| si*Fbl*#2 | Thermo Fisher Scientific | Cat# s201334 |
| si*Fbl*#3 | Thermo Fisher Scientific | Cat# s4820 |
| si*Fbl*#4 | Thermo Fisher Scientific | Cat# s4822 |
| si*Ncl*#1 | Thermo Fisher Scientific | Cat# s70420 |
| si*Ncl*#2 | Thermo Fisher Scientific | Cat# s70421 |
| Custom Stellaris FISH Probes (*Sμ*-Q570, *Sα*-Q670, *Sγ1*-Q670) | LGC Biosearch Technologies | See Table S1 |
| Stellaris FISH Probes, Mouse *Neat1* 5′ Segment with Quasar 570 Dye | LGC Biosearch Technologies | Cat# VSMF-3030-5 |
| Stellaris FISH Probes, Mouse *Gapdh* with Quasar 570 Dye | LGC Biosearch Technologies | Cat# VSMF-3014-5 |
| Recombinant DNA |  |  |
| 3xFNLDD/pCMV-7.1 | Fujita and Fujii, 2012 | RRID: Addgene_48874; Addgene Cat# 48874 |
| 8xLexA-binding elements/pMD20 | Fujita and Fujii, 2011 | RRID: Addgene_48807; Addgene:Cat# 48807 |
| COIL-TOPO vector | This study | Cat# N/A |
| DT-A/loxP/PGK-Neo/loxP | RIKEN BDR (LARGE) | Cat# N/A |
| GFP-Nucleolin | Takagi et al. 2005 | RRID: Addgene_28176; Addgene Cat# 28176 |
| GFP-NPM WT | Wang et al. 2005 | RRID: Addgene_17578; Addgene Cat# 17578 |
| hsMed19_201_pMA-RQ | Thermo Fisher Scientific | Cat# N/A |
| mCardinal-C1 | Addgene | RRID: Addgene_54799; Addgene Cat# 54799 |
| p20-GFP-FHpFBP | Okazaki et al., 2011 | Cat# N/A |
| p3xFLAG-CMV-14 | Sigma Aldrich | Cat# E7908 |
| pBR322 | New England Biolabs (NEB) | Cat# N3033S |
| pCBASce | Richardson et al., 1998 | RRID: Addgene_26477; Addgene Cat# 26477 |
| pCre-ERT2 | Dr. Pierre Chambon (Feil et al.；1997, Indra et al., 1999；Li et al, 2000) | Cat# N/A, RRID N/A |
| pCMV-3xFLAG-NLS-EGFP | This study | Cat# N/A |
| pCMV-3xFLAG-tdMCP-EGFP | This study | Cat# N/A |
| pCMV-3xFLAG-FBL | This study | Cat# N/A |
| pCMV3TAG1A | Agilent Technologies | Cat# 240195 |
| pCMV6-3xFLAG | This study | Cat# N/A |
| pCMV6-Entry Vector | Origene | Cat# PS100001 |
| pCMV6-mAID-3xFLAG | This study | Cat# N/A |
| pCMV6-MED19-MycFLAG | This study | Cat# N/A |
| pCMV6-MED19-MycSpot | This study | Cat# N/A |
| pCMV6-MED19ΔPR-MycSpot | This study | Cat# N/A |
| pCMV6-MED19ΔKNH-MycSpot | This study | Cat# N/A |
| pCMV6-MED19Δ1stKR-MycSpot | This study | Cat# N/A |
| pCMV6-MED19Δ2ndKR-MycSpot | This study | Cat# N/A |
| pCMV6-MED19Δ1stKRΔ2ndKR-MycSpot | This study | Cat# N/A |
| pCMV6-MED19Δ3KR-MycSpot | This study | Cat# N/A |
| pCMV6-MED19ΔC-ter-MycSpot | This study | Cat# N/A |
| pCMV6-MED19-3xFLAG | This study | Cat# N/A |
| pCMV6-MED19Δ3KR-3xFLAG | This study | Cat# N/A |
| pCMV6-MED19ΔC-ter-3xFLAG | This study | Cat# N/A |
| pCMV6-P1-3xFLAG | This study | Cat# N/A |
| pCMV6-P7-3xFLAG | This study | Cat# N/A |
| pCMV6-P20-3xFLAG | This study | Cat# N/A |
| pCMV-ATG-tdMCP-LSM7-3xFLAG | This study | Cat# N/A |
| pCMV-ATG-tdMCP-NOL7-3xFLAG | This study | Cat# N/A |
| pCMV-ATG-tdMCP-WDR74-3xFLAG | This study | Cat# N/A |
| pCMV-COIL-EGFP | This study | Cat# N/A |
| pCMV-COIL-mCardinal | This study | Cat# N/A |
| pCMV-EGFP-FBL | This study | Cat# N/A |
| pCMV-LSM7-3xFLAG | This study | Cat# N/A |
| pCMV-LSM7-3xFLAG-P2A-Puro | This study | Cat# N/A |
| pCMV-mAID.FLAG.myc | Honjo Laboratory | Cat# N/A |
| pCMV-NOL7-3xFLAG | This study | Cat# N/A |
| pCMV-tdMCP-LSM7-3xFLAG | This study | Cat# N/A |
| pCMV-tdMCP-LSM7-3xFLAG-P2A-Puro | This study | Cat# N/A |
| pCMV-tdMCP-MOCK-3xFLAG | This study | Cat# N/A |
| pCMV-tdMCP-MOCK-3xFLAG-P2A-Puro | This study | Cat# N/A |
| pCMV-tdMCP-NOL7-3xFLAG | This study | Cat# N/A |
| pCMV-tdMCP-WDR74-3xFLAG | This study | Cat# N/A |
| pCMV-tdMCP-WDR74-3xFLAG-P2A-Puro | This study | Cat# N/A |
| pCMV-WDR74-3xFLAG | This study | Cat# N/A |
| pCMV-WDR74-3xFLAG-P2A-Puro | This study | Cat# N/A |
| pDT-A/loxP/PGK-EGFP-IRES-Puro/loxP | This study | Cat# N/A |
| pDT-A/8xLexA-Sα-core-EGFP-Puro | This study | Cat# N/A |
| pDT-A/Sirius-8xMS2-Sα-core-EGFP-Puro | This study | Cat# N/A |
| pEF1α-IRES-ZsGreen1 | Takara Bio | Cat# 631976 |
| pEF-3xFLAG-FBL-IRES-Puro | This study | Cat# N/A |
| pEF-FBL-91D3 | This study | Cat# N/A |
| pEF-FBL-195D5 | This study | Cat# N/A |
| pEF-FBL-ΔGAR | This study | Cat# N/A |
| pEF-FBL-ΔGAR91 | This study | Cat# N/A |
| pEF-FBL-ΔGAR195 | This study | Cat# N/A |
| pEF-FBL-WT (pEF-FBL-Kd-Resistant-WT) | This study | Cat# N/A |
| pEF-FBL-ΔGAA167-169 (Mut2) | This study | Cat# N/A |
| pEF-FBL-EDAR215-218AAAA/DV236-237AA (Mut11) | This study | Cat# N/A |
| pEF-FBL-ΔVEF190-192 (Mut14) | This study | Cat# N/A |
| pEF-FBL-K143A/K265A (Mut15) | This study | Cat# N/A |
| pEF1α-FNLDD-IRES-ZsGreen1 | Husain et al., 2020 | Cat# N/A |
| pEF-IRES Puro | Amin et al., 2023 | Cat# N/A |
| pEF-NOL7-3xFLAG-P2A-Puro | This study | Cat# N/A |
| pEF-mAID-ER-3xFLAG-IRES-Puro | Honjo Laboratory | Cat# N/A |
| pEF-ΔC-AID-ER (JP8Bdel-ER)-IRES-Puro | Honjo Laboratory | Cat# N/A |
| pEF1α-IRES-ZsGreen1 Vector | Clontech | Cat# 631976 (Discontinued) |
| pEF-tdMCP-NOL7-3xFLAG-P2A-Puro | This study | Cat# N/A |
| pEF-tdMCP-WDR74-3xFLAG-P2A-Puro | This study | Cat# N/A |
| pEGFP-C2 | Clontech | Cat# N/A (Discontinued) |
| pEGFP-N1 | Clontech | Cat# N/A (Discontinued) |
| pEGFP-3xFLAG-FBL | This study | Cat# N/A |
| pFB-mAIDER-EGFP-IRES-Puro | Honjo Laboratory | Cat# N/A |
| pHR-TRE3G-NLS-tdMCP-EGFP | This study | Cat# N/A |
| pHR-TRE3G-NLS-tdMCP-mCherry | Shao et al., 2016 | RRID: Addgene_99909; Addgene Cat# 99909 |
| pHR-TRE3G-NLS-tdPCP-EGFP | Shao et al., 2016 | RRID: Addgene_99910; Addgene Cat# 99910 |
| pLenti-FNLS-P2A-Puro | Zefra et al., 2018 | RRID: Addgene_110841; Addgene Cat# 110841 |
| pMSCV-mouseR24W(P7)-IRES.DsRed | Honjo Laboratory | Cat# N/A |
| pMSCV-mouseW68R(P1)-IRES.DsRed | Honjo Laboratory | Cat# N/A |
| pPUR-hU6-sgRNA-Sirius-8xMS2 | Ma et al., 2018 | RRID: Addgene_121942; Addgene Cat# 121942 |
| pSPgRNA | Perez-Pinera et al., 2013 | RRID: Addgene_47108; Addgene Cat# 47108 |
| pX335-U6-Chimeric_BB-CBh-hSpCas9n(D10A) | Cong et al., 2013 | RRID: Addgene_42335; Addgene Cat# 42335 |
| pX335-U6-Sα-#11_Chimeric_BB-CBh-hSpCas9n(D10A) | This study | Cat# N/A |
| pX335-U6-Sα-#21_Chimeric_BB-CBh-hSpCas9n(D10A) | This study | Cat# N/A |
| Software and algorithms |  |  |
| bcl2fastq2 | Illumina | RRID: SCR_015058; v2.20 |
| CHOPCHOP | Montague et al., 2014; Labun et al., 2019 | RRID: SCR_015723; https://chopchop.cbu.uib.no/ |
| clusterProfiler (ver. 4.14.6) | Bioconductor | RRID: SCR_016884; https://bioconductor.org/packages/clusterProfiler/ |
| Custom Alt-R CRISPR Design Tool | Integrated DNA Technologies (IDT) | https://www.idtdna.com/site/order/designtool/index/CRISPR_CUSTOM |
| DIA-NN (version 1.9) | Demichev et al., 2020 | RRID: SCR_022865; https://github.com/vdemichev/diann |
| DRAGEN Bio-IT Platform | Illumina | RRID: SCR_028815; v3.7.5 |
| GraphPad Prism 9 (9.1.0, 11.1.0) | GraphPad | RRID: SCR_002798 |
| ImageJ (1.53t) | Schneider et al., 2012 | RRID: SCR_003070; https://ImageJ.net |
| Jetset scoring system | Li et al., 2011 | http://www.cbs.dtu.dk/biotools/jetset/ |
| Kaplan-Meier Plotter | Győrffy et al., 2026 | RRID: SCR_018753; http://www.kmplot.com |
| Mascot search engine | Matrix Science | RRID: SCR_014322 |
| MSigDB GO Gene Sets | Liberzon et al., 2011 | RRID: SCR_016863. m5.go.v2025.1.Mm.symbols.gmt |
| MSigDB GO Gene Sets | Liberzon et al., 2011 | RRID: SCR_016863. m5.go.cc.v2025.1.Mm.symbols.gmt |
| NCBI BLAST | NCBI | RRID: SCR_004870; https://blast.ncbi.nlm.nih.gov/ |
| NovaSeq Control Software | Illumina | Version 1.7.5 |
| org.Mm.eg.db (ver. 3.20.0) | Bioconductor | https://bioconductor.org/packages/org.Mm.eg.db](https://bioconductor.org/packages/org.Mm.eg.db) |
| Percolator | Käll et al., 2007 | RRID: SCR_005040; https://github.com/percolator/percolator |
| Perseus (version 3.0.1.0) | Max Planck Institute of Biochemistry | RRID: SCR_015753; https://maxquant.net/perseus/ |
| Proteome Discoverer 1.2 (in Figure 1, iChIP) | Thermo Fisher Scientific | RRID: SCR_014477 |
| Proteome Discoverer 2.5 (in Figure 6, FBL interactome) | Thermo Fisher Scientific | RRID: SCR_014477 |
| R (version 4.4.3) | R Project for Statistical Computing | RRID: SCR_001905; https://www.r-project.org/ |
| R studio (version. 2024.12.1+563) | Posit Software, PBC | RRID: SCR_000432; https://posit.co/products/open-source/rstudio/ |
| Real Time Analysis (RTA) | Illumina | RRID: SCR_014332; v3.4.4 |
| Stellaris FISH Probe Designer | LGC Biosearch Technologies | https://www.biosearchtech.com/ |
| Xcalibur | Thermo Fisher Scientific | RRID: SCR_014593 |
| Other |  |  |
| 18 mm Poly-L-lysine Coated Coverslips, #1.5 | Neuvitro | Cat# GG-18-15-PLL |
| ABI 3500xL Genetic Analyzer | Thermo Fisher Scientific | RRID: SCR_021901 |
| Advance nanoLC | Michrom BioResources | RRID: N/A |
| AMPure XP beads | Beckman Coulter | Cat# A63880 |
| Anti-FLAG M2 Affinity Gel | Sigma-Aldrich | Cat# A2220 |
| Anti-FLAG M2 magnetic beads | Sigma-Aldrich | Cat# M8823 |
| Applied Biosystems 7500 | Thermo Fisher Scientific | RRID:SCR_018051 |
| Bioruptor II (TYPE12) | BM Bio | Cat# BR2012A |
| Bolt 4–12% Bis-Tris Plus Gels | Thermo Fisher Scientific | Cat# NW04120BOX |
| BZ-X800 confocal microscope | Keyence | RRID: SCR_023617 |
| C18 reversed-phase column (75 μm × 150 mm) | Nikkyo Technos | Cat# NTCC-360/75-3-155 |
| Covaris Focused-Ultrasonicator | Covaris | Cat# S2 |
| Covaris milliTUBE 1 ml | Covaris | Cat# 520128 |
| Covaris Holder milliTUBE 1 ml | Covaris | Cat# 500371 |
| Dynabeads M-280 Tosylactivated | Thermo Fisher Scientific | Cat# DB14203 |
| Dynabeads MyOne Streptavidin C1 | Thermo Fisher Scientific | Cat# DB65001 |
| Dynabeads Protein G | Thermo Fisher Scientific | Cat# DB10004 |
| EASY-nLC 1200 system | Thermo Fisher Scientific | RRID: SCR_026978 |
| FACSCalibur flow cytometer | BD Biosciences | RRID: SCR_000401 |
| FACSAria II | BD Biosciences | RRID: SCR_018934 |
| FACSMelody cell sorter | BD Biosciences | RRID: SCR_023209 |
| GL-Tip SDB | GL Sciences | Cat# 7820-11200 |
| HTC-PAL autosampler | CTC Analytics | RRID: N/A |
| ImageQuant LAS4000 | Cytiva (GE Healthcare) | Cat# 28-9558-10 |
| LTQ Orbitrap Velos mass spectrometer | Thermo Fisher Scientific | RRID: N/A |
| Maxwell RSC | Promega | Cat# AS4500 |
| Mini-PROTEAN TGX Precast Gels (4-15%) | Bio-Rad | Cat# 4561086 |
| Mini-PROTEAN TGX Precast Gels (Any KD) | Bio-Rad | Cat# 4569036 |
| NovaSeq 6000 | Illumina | Cat# 20012850; RRID: SCR_016387 |
| Nucleofector 2b / 96-well Shuttle System | Lonza | Cat# AAB-1001 (Nucleofector 2b); Cat# AAM-1001S (Amaxa 96 well Shuttle Device) |
| Orbitrap Fusion nanoLC/MS system | Thermo Fisher Scientific | RRID:SCR_018702 |
| QuantStudio 5 | Thermo Fisher Scientific | RRID: SCR_020240 |
| Sera-Mag SpeedBead carboxylate-modified [E3] magnetic particles | Cytiva | Cat# 65152105051150 |
| Sera-Mag™ SpeedBead Carboxylate-Modified [E7] Magnetic Particles | Cytiva | Cat# 45152105051150 |
| SoftLink Soft Release Avidin Resin | Promega | Cat# V7841 |
| timsTOF HT Mass Spectrometer | Bruker | RRID: SCR_026543 |

## EXPERIMENTAL MODEL AND SUBJECT DETAILS

### Cell lines

The following cell lines were used in this study: CH12 (mouse, female, B lymphoma-derived), HeLa (human, female, cervical carcinoma-derived), HEK293T (human, female, embryonic kidney-derived), and H1299dA3-1^33^ (human, male, non-small cell lung cancer-derived). HeLa cells were used in imaging of EGFP, EGFP-FBL, EGFP-NCL, and EGFP-NPM1 expression. HEK293T cells were primarily employed for the overexpression of FLAG-tagged FBL and MED19 plasmids and subsequent co-immunoprecipitation (co-IP) assays as well as RNA and DNA pulldown assay. None of the cell lines used in this study were authenticated. CH12 cells harbor an allele-specific aberrant promoter driving constitutive 3′ S*α*–C*α* baseline transcription (Figure S6).

### Mouse strains

Wild-type (WT) female C57BL/6NCrSlc mice were obtained from Japan SLC and housed under specific-pathogen-free conditions. All mice were euthanized between 6–12 weeks of age for splenocyte isolation. All animal housing, handling, and tissue collection procedures were conducted in accordance with institutional guidelines established by the Kyoto University Animal Experimentation Committee, and protocols were approved by the Animal Research Committee of Graduate School of Medicine, Kyoto University (approval no. MedKyo: 26041).

### Human Public Datasets

Patient survival and gene expression analyses were conducted using publicly available, de-identified datasets retrieved from the Kaplan–Meier Plotter database (https://kmplot.com/). No primary human specimens were collected or directly handled in this study; thus, additional local ethical approval was not required.

## METHOD DETAILS

### Cell culture and cytokine-induced class switch recombination (CSR)

CH12, HeLa, HEK293T, and H1299dA3-1^33^ cells were cultured at 37°C in a humidified incubator containing 5% CO_2._ HEK293T cells were maintained in Dulbecco’s Modified Eagle Medium (DMEM) supplemented with 10% fetal bovine serum (FBS) and 1% penicillin-streptomycin. HeLa cells were maintained in DMEM (for imaging experiments) or RPMI-1640 (for RNA and DNA pull-down assays) supplemented with 10% FBS and 1% penicillin-streptomycin (omitted where appropriate). CH12 cells were maintained in RPMI-1640 medium supplemented with 10% FBS, 10% Medium NCTC-109 (1x), 1% penicillin-streptomycin, and 50 µM 2-mercaptoethanol. H1299dA3-1 cells were cultured in RPMI-1640 supplemented with 10% FBS and 1% penicillin-streptomycin.

Induction of class switch recombination in CH12 cells by CD40L, IL-4, and TGF-β (CIT) was performed as previously described.^104^ The efficiencies of CSR to IgA and IgG3 (percentage of IgA^+^ and IgG3^+^ cells) were evaluated at the indicated time points post-CIT stimulation using a FACSCalibur flow cytometer (BD Biosciences). Surface IgA was detected by staining with a PE-conjugated goat anti-mouse IgA antibody, whereas surface IgG3 was detected using a biotin-conjugated anti-mouse IgG3 antibody, followed by PE-conjugated streptavidin.

### Generation of 8xLexA-*Sα*-core Knock-in CH12 Cells

#### Construction of the targeting vector and the Cas9/sgRNA vector

To generate the targeting vector for knock-in of 8xLexA-binding sequence into the *Sα*-core region of the *IgH* locus in CH12 cells, the pDT-A/loxP/PGK-EGFP-IRES-Puro/loxP backbone vector was first constructed by subcloning the EGFP-IRES-Puro cassette from pFB-mAIDER-EGFP-IRES-Puro into the DT-A/loxP/PGK-Neo/loxP vector using overlap extension PCR and NotI/SwaI restriction sites. Next, the 5′ homology arm corresponding to the *Sα*-core region was amplified from CH12 genomic DNA and inserted into the NdeI and ClaI sites of 8xLexA-binding elements/pMD20^105^ to generate a fusion of the 5′ homology arm and the 8xLexA binding sequence (5′ homology arm-inserted 8xLexA/pMD20). The PCR-amplified 3′ homology arm was inserted into pDT-A/loxP/PGK-EGFP-IRES-Puro/loxP to yield pDT-A/loxP/PGK-EGFP-IRES-Puro/3′ arm. Finally, the AscI/NotI-digested 5′ homology arm–8xLexA binding sequence fragment was inserted into pDT-A/loxP/PGK-EGFP-IRES-Puro/3′ arm. The completed targeting donor vector was designated as pDT-A/8xLexA-*Sα*-core-EGFP-Puro.

The pX335-U6-Chimeric_BB-CBh-hSpCas9n(D10A) vector,^106^ encoding the D10A nickase mutant of hSpCas9 and an sgRNA expression cassette, was constructed by inserting annealed double-stranded oligonucleotides targeting the *Sα*-core region into BbsI-digested pX335. The resulting plasmid was named pX335-U6-S*α*-#21_Chimeric_BB-CBh-hSpCas9n(D10A). All oligonucleotide sequences, including qPCR primers, sgRNAs, and genotyping primers, are listed in Data S1.

#### 8xLexA motif knock-in into Sα-core CH12 cells

The 8xLexA motif was knocked in using CRISPR/Cas9-mediated gene editing, as previously described,^27,28^ with minor modifications (Figure S1). The targeting donor vector pDT-A/8xLexA-*Sα*-core-EGFP-Puro and pX335-U6-S*α*-#21_Chimeric_BB-CBh-hSpCas9n(D10A) were constructed using oligonucleotides described in Data S1 (Figure S1A). They were co-electroporated using the Amaxa 96-well Shuttle system and the SF Cell Line 96-well Nucleofector Kit (Lonza). Stable transfectants were selected using 10 μg/mL puromycin. GFP-positive cells were isolated using a FACSAria II cell sorter and seeded by limiting dilution for single-cell cloning (Figure S1B). The isolated clones were evaluated for CIT-induced IgA switching efficiency (Figure S1C) and analyzed by Southern blotting following EcoRI digestion using a DIG-labeled probe (Figure S1D).

To delete the PGK-EGFP-IRES-Puro selection cassette, selected clones (#12 and #16) were subsequently electroporated with the pCre-ERT2,^107–109^ a kind gift from Dr. Pierre Chambon. The following day, cells were treated with 4-hydroxytamoxifen (4-OHT) to induce Cre activity. The GFP-negative population was sorted using a FACSAria II cell sorter (Figure S1E) and subjected to single-cell cloning by limiting dilution. Successful excision was verified by an IgA switching assay and Southern blotting (Figure S1F).

#### Construction and introduction of FLAG-tagged LexA-binding protein fused with NLS into 8xLexA-Sα-core CH12 cells

The LexA DNA-binding domain (LDD) fused with 3xFLAG and a nuclear localization signal (NLS), derived from 3xFNLDD/pCMV-7.1,^110^ was inserted into the pEF1α-IRES-ZsGreen1 vector. Briefly, the 3xFNLDD fragment was excised from 3xFNLDD/pCMV-7.1 using SacI and SalI, and the ends were blunted. The pEF1α-IRES-ZsGreen1 vector was digested with EcoRI and similarly blunted. Following ligation, the resulting construct (pEF1α-FNLDD-IRES-ZsGreen1) was linearized with AseI and electroporated into the selected 8xLexA knock-in clones to generate “FNLDD-8xLexA-*Sα*-core” cell lines. As a negative control, the pEF1α-FNLDD-IRES-ZsGreen1 vector alone was introduced into parental CH12 cells to generate “FNLDD-alone” cells (Figure S1G). Target enrichment of FNLDD at the *Sα*-core region was validated by a ChIP assay using an anti-FLAG M2 antibody, as previously described^111^ (Figure S1H). FNLDD-8xLexA-*Sα*-core clone #12-9 cells was used for subsequent studies.

### SILAC-iChIP Assay and Mass Spectrometry

Insertional chromatin immunoprecipitation (iChIP) was performed as previously described,^27,28^ with minor modifications (Figure S1). Briefly, for SILAC (Stable Isotope Labeling by Amino acids in Cell culture) labeling, FNLDD-8xLexA-*Sα*-core cells were cultured in media containing “heavy” [^13^C_6,_ ^15^N_2]_-L-lysine and [^13^C_6,_ ^15^N_4]_-L-arginine, whereas FNLDD-alone control cells were cultured in standard medium containing unlabeled “light” amino acids. Reverse-crosslinked proteins were separated by SDS-PAGE and analyzed by LC-MS/MS, as previously described. Measurements were performed using an LTQ Orbitrap Velos mass spectrometer (Thermo Fisher Scientific) interfaced with an Advance nanoLC (Michrom BioResources) and an HTC-PAL autosampler (CTC Analytics) at the Research Institute for Microbial Diseases (RIMD, Osaka University, Osaka, Japan). Data acquisition was performed using Xcalibur software (Thermo Fisher Scientific). Peptide identification and SILAC-based quantification were executed using Proteome Discoverer (v1.2; Thermo Fisher Scientific) combined with the Mascot search engine (Matrix Science) against the Swiss-Prot database. Search criteria included trypsin/P digestion with up to two missed cleavages, precursor ion mass tolerance of 10 ppm, and fragment ion mass tolerance of 0.8 Da. Carbamidomethylation of cysteine was specified as a fixed modification, whereas N-terminal protein acetylation and methionine oxidation were selected as variable modifications. Protein quantification required a minimum of two identified peptides per protein. Two independent experiments were conducted.

### Proteomics Data Analysis for the iChIP Assay

For mass spectrometry (MS) data analysis of 8xLexA-binding proteins, a total of 234 proteins were successfully quantified with Heavy/Light abundance ratios across two independent experimental replicates. Data from the two replicates were averaged for each protein and ranked according to their Heavy/Light enrichment ratios.

Based on the mean Heavy/Light ratio, the top 50 enriched proteins were selected for downstream functional analysis. Relative abundances (Heavy/Light ratios) were visualized as a heatmap using the *pheatmap* package (Figure 1C).

Gene Ontology (GO) enrichment analysis was performed in the R statistical environment (v4.4.0) using the *org.Mm.eg.db* annotation package (v3.20.0) and the enrichGO function from the *clusterProfiler* package (v4.14.6). GO terms with a false discovery rate (FDR) *q*-value < 0.05 were subjected to redundancy reduction using the simplify function with a similarity threshold of 0.7. Subsequently, terms satisfying a fold enrichment > 15 and a gene ratio ≥ 5/50 were retained for the GO term functional summary (Figure 1D) and enrichment dot plots (Figure 1E).

Gene Set Enrichment Analysis (GSEA) was performed across all 234 proteins using the gseGO function from the *clusterProfiler* package with the MSigDB gene set collection (m5.go.v2025.1.Mm.symbols.gmt). GSEA enrichment plots for “rRNA processing” and “rRNA binding” were generated (Figures 1F and 1G).

### CRISPR Interference (CRISPRi) in CH12 Cells

CRISPRi experiments were performed as previously described,^66^ with minor modifications. Briefly, sgRNAs targeting the transcription start sites of candidate genes were designed using CHOPCHOP^112,113^ and the Integrated DNA Technologies (IDT) Custom Design Tool. The sequences of the sgRNAs are listed in Data S1. Annealed sgRNA oligos were cloned into the pSPgRNA vector,^114^ and plasmid sequences were confirmed by Sanger sequencing. The pBR322 vector was used as a control (Ctrl). Individual sgRNA plasmids were electroporated into CH12-dCas9-KRAB (#40) cells, which were cultured in medium supplemented with blasticidin S to maintain selection for dCas9-KRAB expression.^66^ For specific experiments, the sgRNA plasmids were also electroporated into *Sα*-core *Sirius-8xMS2* reporter subclones (CH12-dCas9-KRAB-*Sirius-8xMS2*-*Sα*-core-3xFLAG-tdMCP-EGFP (#95)). A combination of two sgRNAs (sg*Med19*-a and sg*Med19*-b) was used for *Med19* knockdown, unless indicated otherwise.

### siRNA-Mediated Knockdown

siRNAs targeting the nucleolin (*Ncl*) and fibrillarin (*Fbl*) genes were electroporated using a Nucleofector device (Nucleofector 2b/96-well Shuttle System) and SF Cell Line 96-well Nucleofector Kit using program CM-150. siRNA IDs are listed in Data S1.

### RNA Isolation and Reverse Transcription-Quantitative PCR (RT-qPCR)

Total RNA was extracted using Sepasol-RNA II Super or the Maxwell RSC simplyRNA Cells Kit. To eliminate genomic DNA, in-tube DNase I treatment was performed during RNA isolation. RNA concentration was measured using a NanoDrop spectrophotometer. cDNA was synthesized using SuperScript IV Reverse Transcriptase with random hexamers. Quantitative PCR was performed using an Applied Biosystems 7500 Real-Time PCR System or QuantStudio 5 with PowerUp SYBR Green Master Mix. Primer sequences are listed in Data S1.

### Western Blotting

Western blotting was performed as previously described.^66^ Proteins were separated using Bolt 4–12% Bis-Tris Plus Gels (Thermo Fisher Scientific), Mini-PROTEAN TGX Precast Gels (4–15%; Bio-Rad), or Mini-PROTEAN TGX Precast Gels (Any kD; Bio-Rad) according to the manufacturers’ instructions. The specific antibodies used in this study are listed in the Key Resources Table. Chemiluminescence signals were developed using Chemi-Lumi One L or Chemi-Lumi One Super on an ImageQuant LAS4000 imaging system.

### Cell Proliferation Assay

Cell proliferation capacity was evaluated using the CellTrace CFSE Cell Proliferation Kit for flow cytometry. A total of 1.5 × 10^6^ cells were stained with 1 μM CFSE in 3 mL of PBS(-) supplemented with 0.5% FCS for 15 min at room temperature (RT). Quenching was performed by adding 14 mL of ice-cold culture medium and incubating the suspension on ice for 5 min. After centrifugation, cells were resuspended in 6 mL of fresh, pre-warmed culture medium and incubated at 37°C for 10 min. The baseline staining efficiency was evaluated immediately using flow cytometry (0 h). Cells were then cultured with or without CIT stimulation, and CFSE fluorescence intensity was measured 24 and 48 h post-treatment. The geometric mean fluorescence intensity of CFSE was normalized to that of the corresponding control or WT cells (set to 1) in each comparison for statistical analysis.

### γH2AX Foci Formation Assay

HeLa cells were seeded onto Lab-Tek 8-chambered glass slides and transfected with EGFP-fused FBL, NCL (GFP-Nucleolin),^115^ or NPM1 (GFP-NPM WT)^116^ constructs using ViaFect Transfection Reagent. At 24 h post-transfection, cells were treated with 4 μM etoposide for 4 h, fixed with 4% paraformaldehyde for 10 min at RT, and stained for γH2AX using the DNA Damage Detection Kit - γH2AX - Deep Red. Slides were mounted with ProLong Glass Antifade Mountant and imaged using a BZ-X800 fluorescence microscope.

### Co-immunoprecipitation (Co-IP)

#### Co-IP in HEK293T cells (Figures S2B and S2C)

HEK293T cells were transfected with a plasmid expressing 3xFLAG-tagged mouse AID (pCMV6-mAID-3xFLAG) using FuGENE 6 and harvested 48 h post-transfection. Cells were lysed in RIPA buffer (10 mM Tris-HCl, pH 7.5, 150 mM NaCl, 0.5 mM EDTA, 0.1% SDS, 1% Triton X-100, and 1% sodium deoxycholate) supplemented with 1x c*O*mplete Protease Inhibitor Cocktail and 150 Kunitz U/mL DNase I for 30 min on ice. Lysates were centrifuged at 20,000 × *g* for 10 min at 4°C, and the supernatant was diluted with four volumes of Dilution Buffer (10 mM Tris-HCl, pH 7.5, 150 mM NaCl, and 0.5 mM EDTA). Immunoprecipitation was performed overnight at 4°C using an anti-FLAG M2 antibody. Dynabeads Protein G were then added for an additional 2 h, followed by six washes with Washing Buffer (10 mM Tris-HCl, pH 7.5, 150 mM NaCl, 0.5 mM EDTA, and 0.05% IGEPAL CA-630). Bound proteins were eluted in 1x SDS Sample Buffer (62.5 mM Tris-HCl, pH 6.8, 2% SDS, 10% glycerol, 0.004% bromophenol blue, and 50 mM DTT) and analyzed by western blotting.

#### Co-IP in CH12 Cells (Figures S2G–S2H)

AID knockout (AID-KO) CH12 cells^117^ were lysed in Lysis Buffer (10 mM HEPES, pH 7.4, 100 mM KCl, 5 mM MgCl_2,_ 20 mM EDTA, 0.5% IGEPAL CA-630, 1% Triton X-100, 0.05% sodium deoxycholate, and 10% glycerol) supplemented with 1x c*O*mplete Protease Inhibitor Cocktail. Cleared lysates were diluted with nine volumes of KCl Buffer (20 mM HEPES, pH 7.4, 100 mM KCl, 1 mM MgCl_2,_ 0.2 mM CaCl_2,_ 0.2 mM EDTA, 0.1% IGEPAL CA-630, and 10% glycerol) and incubated with anti-FLAG M2 Affinity Gel for 3 h at 4°C. The beads were washed six times with Washing Buffer (50 mM Tris-HCl, pH 7.4, 150 mM NaCl, 1 mM MgCl_2,_ 0.5 mM EDTA, and 0.05% IGEPAL CA-630, supplemented with 1x cOmplete Protease Inhibitor Cocktail). Bound proteins were eluted in SDS Sample Buffer and detected by western blotting.

#### RNase A–Treated Co-IP in CH12 Cells (Figure S2I)

Cells were lysed in ice-cold Lysis Buffer (10 mM HEPES, pH 7.4, 100 mM KCl, 5 mM MgCl_2,_ 0.5% IGEPAL CA-630, 1% Triton X-100, 0.05% sodium deoxycholate, and 10% glycerol) supplemented with 1x cOmplete Protease Inhibitor Cocktail. Cleared lysates were diluted six-fold with KCl Buffer (20 mM HEPES, pH 7.4, 100 mM KCl, 1 mM MgCl_2,_ 0.2 mM CaCl_2,_ 0.1% IGEPAL CA-630, and 10% glycerol) and incubated with anti-FLAG M2 Affinity Gel for 3 h at 4°C. The beads were washed sequentially with High-Salt Buffer (50 mM Tris-HCl, pH 7.4, 1 M NaCl, 1% IGEPAL CA-630, 1% sodium deoxycholate, and 1x cOmplete Protease Inhibitor Cocktail), Low-Salt Buffer (50 mM Tris-HCl, pH 7.4, 150 mM NaCl, 1% IGEPAL CA-630, 1% sodium deoxycholate, and 1x cOmplete Protease Inhibitor Cocktail), and High-Salt Buffer again. The beads were incubated in 180 μL of RNase Buffer (10 mM HEPES, pH 7.4, 2 mM MgCl_2, 0_.1% IGEPAL CA-630, and 0.05% sodium deoxycholate) with 20 μL of RNase A for 1 h at 37°C, followed by another Low-Salt/High-Salt/Low-Salt wash cycle. Eluted proteins were analyzed by western blotting.

#### Co-IP in CH12 Cells (Figure 4M)

Cells were lysed in WCE Lysis Buffer (10 mM HEPES, pH 7.4, 100 mM KCl, 5 mM MgCl_2, 0_.5 mM EDTA, 0.5% IGEPAL CA-630, 1% Triton X-100, 0.1% SDS, and 10% glycerol) supplemented with 1x cOmplete Protease Inhibitor Cocktail. Lysates were sonicated for 11 cycles (15 s ON / 60 s OFF) using a Bioruptor, centrifuged, and diluted with four volumes of NT2 Dilution Buffer (50 mM Tris-HCl, pH 7.6, 150 mM NaCl, 1 mM MgCl_2, a_nd 1 mM EDTA). Immunoprecipitation was performed overnight at 4°C using an anti-FLAG M2 antibody and BSA-blocked Dynabeads Protein G. Beads were washed six times with Washing Buffer (20% WCE lysis buffer and 80% NT2 dilution buffer), eluted in SDS Sample Buffer, and analyzed by western blotting.

### Plasmid Construction for AID Mutants

To generate wild-type (WT) and mutant AID expression constructs (pCMV6-mAID-3xFLAG, pCMV6-P7-3xFLAG, pCMV6-P1-3xFLAG, and pCMV6-P20-3xFLAG), the pCMV6-3xFLAG backbone was first constructed by subcloning the 3xFLAG sequence from p3xFLAG-CMV-14 into the pCMV6-Entry Vector using EcoRI and BstXI restriction sites. Next, WT and mutant AID fragments were PCR-amplified from pCMV-mAID.FLAG.myc, pMSCV-mouseR24W(P7)-IRES.DsRed, pMSCV-mouseW68R(P1)-IRES.DsRed, and P20-GFP-FHpFBP using forward and reverse primers containing EcoRI and SalI sites, respectively (Data S1). Amplified fragments were digested with EcoRI/SalI and inserted into EcoRI/SalI-linearized pCMV6-3xFLAG using Ligation high (Toyobo).

### Evaluation of Germline Transcripts

Total RNA was isolated as described above. Expression levels of specific germline transcripts were quantified by RT-qPCR using primers listed in Data S1.

### Somatic Hypermutation (SHM) Assay

CH12 cells were transfected with si*Fbl*#1, si*Fbl*#2, or control siRNAs and stimulated with CIT. Genomic DNA was isolated 48 h post-stimulation using phenol/chloroform extraction and diluted to 100 ng/μL in 10 mM Tris-HCl, pH 7.5. A 399-bp fragment at the 5′ region of the *Sμ*-core was amplified using Q5 High-Fidelity Master Mix with primers listed in Data S1. PCR products were treated with ExoSAP-IT, purified using AMPure XP beads, cloned into the Zero Blunt TOPO vector, and sequenced using M13 forward primers on an ABI 3500xL Genetic Analyzer, as previously described.^66^ (Tables S1 and S2).

### DNA Cleavage Assay with Biotin-dUTP Labeling

The DNA cleavage assay was performed as previously described, with minor modifications.^18,66^ Briefly, cultured cells were subjected to dead cell removal using the MojoSort Dead Cell Removal Kit. A total of 5 × 10^6^ viable cells were fixed at RT in STF Fixation Buffer (3% 2-bromo-2-nitropropane-1,3-diol, 3% diazolidinyl urea, 1.2% zinc sulfate heptahydrate, 19.7 mM trisodium citrate dihydrate, and 50 mM EDTA) for 15 min. Cells were then washed once with 1x PBS(-), once with Buffer A (10 mM HEPES, pH 6.5, 10 mM EDTA, and 0.25% Triton X-100), and once with Buffer B (10 mM HEPES, pH 6.5, 200 mM NaCl, and 1 mM EDTA). Subsequently, cells were resuspended in Buffer C (100 mM Tris-HCl, pH 7.6, 50 mM EDTA, and 1% Triton X-100) and incubated on ice for 30 min for cytoplasmic lysis.

Fixed cells were washed sequentially with Buffer D (100 mM Tris-HCl, pH 7.4, and 150 mM NaCl) and PNK Buffer, followed by incubation with T4 Polynucleotide Kinase (PNK) for 1 h at 37°C. After washing with Buffer D and terminal deoxynucleotidyl transferase (TdT) buffer containing BSA, cells were treated with TdT and biotin-16-dUTP in the presence of 0.25 mM CoCl_2 f_or 1 h at 37°C. After washing with Buffer D containing 0.01% Tween-20, cells were treated with 2 μL of RNase A in 100 μL of Buffer D at 37°C for 20 min, followed by digestion with 0.2 mg/mL Proteinase K in 500 μL of Lysis Buffer (10 mM Tris-HCl, pH 8.0, 10 mM EDTA, 150 mM NaCl, and 0.1% SDS) overnight at 55°C.

DNA was then fragmented using a Bioruptor II to a size range of 500–1,000 bp (11 cycles of 15 s ON / 90 s OFF). Lysates were supplemented with 250 μL of 5 M NaCl and centrifuged at 20,000 × *g* for 10 min. The DNA-containing supernatant was transferred to a new tube, and DNA was precipitated with 2-propanol. Purified labeled DNA (20 μg) was captured using 10 μL of BSA-blocked Dynabeads MyOne Streptavidin C1 in Trapping Buffer (5 mM Tris-HCl, pH 7.4, 0.5 mM EDTA, 0.015% Tween-20, and 1 M NaCl) for 2 h at RT. Beads were washed for three rigorous washing cycles: Cycle 1 utilized Low-Salt Buffer, High-Salt Buffer, and LiCl Buffers at RT; Cycles 2 and 3 utilized a 55°C incubation for the High-Salt Buffer wash. Bound DNA was eluted in 98 μL of formamide and 2 μL of 0.5 M EDTA, precipitated with 2-propanol in the presence of 0.2 M NaCl and glycogen, and analyzed by qPCR using primers listed in Data S1.

### *IgH*/*c-Myc* Translocation Assay

The *IgH*/*c-Myc* translocation assay was performed as previously described^49,111^ with updated primer sets (Data S1). Nested PCR was conducted using the Expand Long Template PCR System. First-round PCR conditions: initial denaturation at 94°C for 3 min; 30 cycles of 94°C for 15 s and 68 °C for 7 min 15 s (with 20 s added per cycle); final extension at 68 °C for 5 min. One microliter of the product was used for second-round PCR: 94°C for 3 min; 25 cycles of 94°C for 15 s and 68 °C for 7 min 15 s; final extension at 68 °C for 5 min. Southern hybridization was performed using ExpressHyb or ULTRAhyb buffer according to the manufacturer’s instructions.

### Chromatin Conformation Capture (3C) Assay

The 3C assays for Figures 2N and 2O were performed according to a previously described protocol, using custom-designed primer sets (Data S1) that yield PCR product sizes distinct from Figures 4I-4J and 7K–7L.^19,111^ For Figures 4I, 4J, 7K, and 7L, a modified protocol was applied, as outlined below. A total of 7 × 10^6^ cells were fixed in 20 mL of medium containing 0.8% methanol-free formaldehyde for 5 min at RT and quenched with 0.2 M glycine. Cells (3.5 × 10^6^) were lysed in Lysis Buffer (50 mM Tris-HCl, pH 7.6, 150 mM NaCl, 5 mM MgCl_2,_ 0.5% IGEPAL CA-630, and 1% Triton X-100 supplemented with 1x c*O*mplete Protease Inhibitor Cocktail) for 30 min on ice.

Nuclei were washed once with Lysis Buffer and once with 1x M Buffer, then resuspended in 100 μL of 1x M Buffer. To achieve permeabilization while strictly preventing nuclear clumping, 100 μL of 1x M Buffer containing 0.6% SDS was dispensed extremely slowly directly into the suspension. During this addition, the tube was continuously and gently tapped to ensure immediate mixing and keep nuclei uniformly suspended. Nuclei were further incubated at 37°C for 2 h with intermittent shaking (1100 rpm for 15 s every 5 min) to ensure sufficient permeabilization. To quench the SDS, Triton X-100 was added to a final concentration of 1.8%, and cells were incubated for 1 h at 37°C. Chromatin was digested overnight at 37°C with 450 U of HindIII in 1x M Buffer.

Digestion was terminated by adding SDS to a final concentration of 0.85% and incubating at 65°C for 25 min. Digested chromatin equivalent to 2.1 × 10^6^ cells was diluted in 4.5 mL of 1x T4 DNA Ligase Reaction Buffer containing 1% Triton X-100. Intermolecular ligation was performed using T4 DNA Ligase and 0.1 mM ATP at 16°C overnight. Samples were treated with Proteinase K at 55°C for 4 h and reverse-crosslinked overnight at 68°C. DNA was purified by 2-propanol precipitation to remove detergents from the reaction, treated with RNase A, extracted with phenol/chloroform, and resuspended in 10 mM Tris-HCl, pH 7.6, at a concentration of 100 ng/μL.

Semi-quantitative PCR was performed to analyze *Sμ*–S*α* and *Eμ–Eα* interactions, with *Gapdh* serving as a loading control, using two different template DNA amounts per sample to ensure linear amplification. Reactions were carried out using Platinum Taq DNA Polymerase (*Sμ*–S*α* and *Eμ–Eα* interactions) or KOD FX Neo DNA Polymerase (*Gapdh*) with primers listed in Data S1. PCR products were resolved on 2% agarose gels and visualized using ethidium bromide. Band intensities were quantified using ImageJ software^118^ to calculate the (*Sμ–Sα*)/(*Eμ–Eα*) ratio.

### Non-Homologous End Joining (NHEJ) Assay

The NHEJ assay was performed as previously described.^119^ Briefly, 12 pmol of control or human *FBL*-specific siRNAs (si*Fbl*#3 or si*Fbl*#4; Data S1) were electroporated into 1 × 10^5^ H1299dA3-1 cells^120^ using an Amaxa 96-well Shuttle Nucleofector with SF Cell Line 96-well Nucleofector Kit and program EW-127. The following day, cells were transfected with the I-SceI expression plasmid (pCBASce)^121^ using Lipofectamine 3000. At 48 h post-transfection, the percentage of GFP^+^ cells was analyzed using flow cytometry (GFP reversion frequency), or genomic DNA was extracted for qPCR analysis to determine joined and uncut allele frequencies using primers listed in Data S1.

### *Sμ–Sα* Switch Junction Microhomology Sequence Analysis

CH12F3-2A cells were electroporated with si*Fbl*#1, si*Fbl*#2, or control siRNAs and stimulated with CIT 24 h later. After 48 h of stimulation, IgA^+^ (PE-positive) cells were isolated using a BD FACSMelody Cell Sorter. Genomic DNA was extracted, and *Sμ–Sα* junctions were amplified by nested PCR using primers specific for *Iμ* and *Cα* regions (Data S1) with PrimeSTAR GXL DNA Polymerase. Gel-purified PCR products (0.3–2 kb) were cloned into HindIII/BamHI-digested pBluescript vectors and transformed into *Escherichia coli* XL10-Gold. Positive clones were sequenced using an ABI 3500xL Genetic Analyzer. Junction sequences were aligned to the mouse *μ* (AH005309.2) and *α* (D11468.1) switch regions using NCBI BLAST (Data S5).

### Construction of Human FBL Expression Vectors

#### Human FBL Cloning and RNA-Binding Mutants

WT and mutant human *FBL* expression plasmids were generated as described in Data S1. WT *FBL* cDNA was amplified from the cDNA pool derived from HEK293T cells and initially subcloned into the pCMV3TAG1A vector using BamHI and EcoRI restriction sites with Ligation high. The 3xFLAG-tagged *FBL* insert was then transferred into pEF-IRES-Puro^49^ using GeneArt Seamless Cloning Enzyme Mix. An si*Fbl*#1-resistant silent mutation was introduced by site-directed mutagenesis via inverse PCR using primers containing overlapping sequences. The resulting pEF-FBL-Kd-Resistant-WT (pEF-FBL-WT) was subsequently used as a template to generate domain deletions and point mutants by blunt-end self-ligation following 5′ phosphorylation with T4 Polynucleotide Kinase and Ligation high. All constructs were verified using Sanger sequencing.

#### FBL Catalytic Mutants

FBL catalytic mutants were generated as detailed in Data S1, using pEF-FBL-WT as the parental template. Mut2, Mut11, and Mut15 were generated using site-directed mutagenesis. For Mut11 and Mut15, which harbor two distant mutation sites, mutations were introduced sequentially in two steps. Mut14 was constructed by assembling two overlapping PCR fragments (Part 1 and Part 2) harboring the target mutation using GeneArt Gibson Assembly HiFi Master Mix. All constructs were verified using Sanger sequencing.

#### EGFP-FBL Expression Plasmid

3xFLAG-tagged FBL (WT) was inserted into the pEGFP-C2 vector using Gibson Assembly HiFi Master Mix and primers listed in Data S1.

### Knockdown-Rescue Experiments in CH12 Cells

Linearized pEF-FBL expression vectors (encoding WT or mutant human *FBL*) were electroporated into CH12 cells, followed by selection with 1 μg/mL puromycin for 7–10 days. The resulting stable cell lines were subsequently transfected with si*Fbl*#1 to specifically deplete endogenous mouse FBL.

### RNA Immunoprecipitation (RIP)

A total of 1 × 10^7^ cells were lysed in IGEPAL CA-630 Lysis Buffer (50 mM HEPES-KOH, pH 7.4, 100 mM KCl, 5 mM MgCl_2,_ 0.1% IGEPAL CA-630, 0.01% sodium deoxycholate, 10% glycerol, 1 mM DTT, 150 Kunitz U/mL DNase I, 1x cOmplete Protease Inhibitor Cocktail, and RNaseOUT) for 30 min on ice. Lysates were sonicated using a Bioruptor II (9 cycles of 30 s ON / 60 s OFF) and cleared by centrifugation. Supernatant was incubated with anti-FLAG M2-conjugated Dynabeads Protein G for 2 h at 4°C. Beads were washed three times with Wash Buffer 1 (containing 100 mM KCl) and three times with Wash Buffer 2 (containing 200 mM KCl). Bound RNA was extracted directly from beads using TRIzol Feagent and RNA Clean & Concentrator-5, reverse-transcribed using SuperScript IV VILO Master Mix, and analyzed by RT-qPCR.

### ChIP Assay

Chromatin was prepared from 2.5 × 10^7^ cells using the truChIP Chromatin Shearing Kit, according to the manufacturer’s protocol. Immunoprecipitation was performed overnight at 4°C in IP Buffer (20 mM Tris-HCl, pH 7.4, 150 mM NaCl, 1.5 mM EDTA, 1% Triton X-100, and 0.1% SDS). BSA-blocked Dynabeads Protein G were added to the mixture for 2 h. Bead-chromatin complexes were washed twice with Low-Salt Buffer (containing 150 mM NaCl), twice with High-Salt Buffer (containing 500 mM NaCl), and twice with LiCl Buffer (containing 250 mM LiCl). Chromatin was eluted in Elution Buffer (0.1 M NaHCO_3, 1_% SDS), reverse-crosslinked overnight at 65°C, and DNA was extracted using phenol/chloroform. Recovered DNA was analyzed using qPCR with the primers listed in Data S1.

### Pyridostatin (PDS) Treatment

Pyridostatin hydrochloride (PDS) was dissolved in dimethyl sulfoxide (DMSO). CH12 cells (including WT or knockdown-rescue lines) were treated with the indicated concentrations of PDS concurrently with CIT stimulation to evaluate the effect of G-quadruplex stabilization on CSR efficiency. During flow cytometric analysis of IgA^+^ cells using a BD FACSCalibur, the percentage of dead cells was concomitantly evaluated by gating on the propidium iodide (PI)-positive population.

### R-loop Detection by DNA/RNA Hybrid (S9.6) Immunoprecipitation

CH12 cells were transfected with si*Fbl*#1 or control siRNA and stimulated with CIT for 24 h. Genomic DNA was extracted by incubating cells in Lysis Buffer (10 mM Tris-HCl, pH 8.0, 1 mM EDTA, 0.625% SDS, and 62.5 μg/mL Proteinase K) overnight at 37°C. Extracted DNA was fragmented to a size range of 500–1000 bp using a Bioruptor II (12 cycles of 15 s ON / 90 s OFF) and purified via phenol/chloroform extraction.

DNA was precipitated by adding 150 mM NaCl and an equal volume of 2-propanol. After washing with 80% ethanol, pellet was dissolved in TE Buffer (10 mM Tris-HCl, pH 8.0, and 1 mM EDTA). Fragmentation size was confirmed by running 100 ng of sheared DNA on a 1% agarose gel. Next, 15 μg of sheared DNA was treated with RNase T1 and RNase III in Reaction Buffer (50 mM Tris-HCl, pH 7.5, 150 mM KCl, 3 mM MgCl_2,_ and 1 mM DTT) to eliminate single- and double-stranded RNAs. For negative control samples, RNase H was added alongside RNase T1 and RNase III. RNase-treated DNA was re-purified using phenol/chloroform extraction and resuspended in TE Buffer.

For immunoprecipitation, 4.4 μg of treated DNA was incubated with 2.5 μg of anti-DNA/RNA hybrid S9.6 antibody in DRIP Binding Buffer (10 mM sodium phosphate, pH 7.0, 140 mM NaCl, and 0.1% Tween-20) overnight at 4°C with end-over-end rotation. Dynabeads Protein G (10 μL per sample), pre-blocked with BSA and 5 μg of RNase H-treated herring sperm DNA, were added to the mixture and incubated for 2 h at 4°C. After washing three times with DRIP Binding Buffer, bound hybrids were eluted by incubating beads in 300 μL of DRIP Elution Buffer (50 mM Tris-HCl, pH 7.5, 10 mM EDTA, and 0.5% SDS) containing 7 μL of 20 mg/mL Proteinase K at 55°C for 45 min. Eluate was mixed with 150 μL of 5 M NaCl and centrifuged at 20,000 × *g* for 10 min at 4°C. Supernatant was precipitated with 400 μL of 2-propanol supplemented with 2 μL of 10 mg/mL glycogen. Resulting DNA pellet was dissolved in 80 μL of 10 mM Tris-HCl, pH 7.5, and analyzed by qPCR using primers listed in Data S1.

### RNA and DNA Pull-Down Assay and Circular Dichroism (CD) Spectroscopy

G-quadruplex (G4) formation by *c-Myc* and mutant *c-Myc* (mut *c-Myc*) DNA oligonucleotides was confirmed by circular dichroism (CD) spectroscopy, as previously described (Figure S4I).^122^ CD spectra were recorded using a model J-820 spectropolarimeter (JASCO) with a 0.2 cm pathlength cell at RT. Oligonucleotides were measured at a strand concentration of 8 μM in a buffer containing 100 mM KCl and 50 mM Tris-HCl, pH 7.5. Buffer background spectra were subtracted, and raw data were used without further processing, such as smoothing. Sequences and preparations of other Sμ G4-forming oligonucleotides were identical to those previously described ^21^.

Biotinylated RNA or DNA oligonucleotides (Data S1) were denatured at 95°C for 5 min and allowed to renature at RT to facilitate proper folding. HeLa cells expressing FLAG-tagged *NCL* or *FBL* constructs were harvested in Suspension Buffer (20 mM Tris-HCl, pH 7.5, 100 mM KCl, 1 mM EDTA, and 1x Protease Inhibitor Cocktail) and sonicated. Pre-blocked SoftLink Soft Release Avidin Resin was incubated with 0.1 μM folded oligonucleotides and cell lysates (0.1 mg/mL total protein) in TE Buffer for 1 h at 4°C. Resin was washed with Washing Buffer (50 mM Tris-HCl, pH 7.5, 150 mM KCl, 1 mM EDTA, 1 mM DTT, and 1% Triton X-100). Bound proteins were eluted in SDS Sample Buffer and analyzed by western blotting using an anti-FLAG antibody.

### Reverse Transcription at Low dNTP Concentration (RTL)-qPCR

RTL-qPCR was performed as previously described^41^ with minor modifications. Total RNA was extracted from CH12 cells using TRIzol Reagent, the RNA Clean & Concentrator-25 Kit, or the Maxwell RSC simplyRNA Cells Kit. To stringently eliminate residual genomic DNA, purified RNA was treated with a cocktail of AluI, HaeIII, and TURBO DNase, followed by re-purification using Sepasol-RNA II Super. RNA was reverse-transcribed using low concentrations of dNTPs, gene-specific primers, and enzymes (Data S1). cDNA amplification efficiency was evaluated by qPCR using a QuantStudio 5 Real-Time PCR System. To quantify 2′-*O*-methylation, we calculated the ratio of RT efficiency under high versus low dNTP conditions (High/Low dNTP ratio). A higher ratio reflects enhanced RT stalling induced by site-specific 2′-*O*-methylation.

### Construction of MED19 WT and Mutant Expression Plasmids

Human *MED19* cDNA was synthesized by Thermo Fisher Scientific (hsMed19_201_pMA-RQ; see Key Resources Table), and its codon-optimized sequence is described in Data S1. Synthesized DNA was cloned into the pCMV6-Entry Vector containing a Myc-FLAG tag using EcoRI and XhoI restriction sites. To replace the FLAG tag with a Spot tag, site-directed mutagenesis was performed using primers containing Spot tag sequences, yielding pCMV6-*MED19*-MycSpot.

The resulting pCMV6-*MED19*-MycSpot vector was used as parental plasmid for generating deletion mutants. The deletion constructs (ΔPR, ΔKNH, Δ1stKR, Δ2ndKR, and Δ1KRΔ2KR) were generated by blunt-end self-ligation following 5′ phosphorylation with T4 Polynucleotide Kinase and Ligation high. The Δ3KR and ΔC-ter deletion mutants were subsequently generated from the ΔKNH construct using the same method.

To generate 3xFLAG-tagged expression vectors, pCMV6-*MED19*-3xFLAG was constructed by site-directed mutagenesis using pCMV6-*MED19*-MycFLAG as a template. Next, pCMV6-*MED19*Δ3KR-3xFLAG and pCMV6-*MED19*ΔC-ter-3xFLAG were generated by replacing WT *MED19* coding sequence in pCMV6-*MED19*-3xFLAG via Gibson Assembly using GeneArt Gibson Assembly HiFi Master Mix. All constructed vectors were verified by Sanger sequencing. All oligonucleotides are listed in Data S1.

### Prediction of Intrinsically Disordered Domain of MED19

To evaluate the disordered tendency of MED19, *Arabidopsis thaliana* MED19a, MED19b, and human *MED19* were analyzed using PONDR VSL2 predictor^123^. Amino acid sequences of Q9FMP0 (MD19A_ARATH), Q6NQD9 (MD19B_ARATH), A0JLT2 (MED19_HUMAN) from UniProt were used for *A. thaliana* MED19a, *A. thaliana* MED19b and human MED19, respectively.

### Transcriptome Analysis of *Med19*-Knockdown CH12 Cells

#### Med19 Knockdown and RNA Extraction

Biological duplicate total RNA samples from the *Med19*-knockdown (using dual sgRNAs of sg*Med19*-a and sg*Med19*-b) and control CH12-dCas9-KRAB (#40) cells were prepared by using a Maxwell RSC with the Maxwell RSC simply RNA Cells Kit. RNA integrity (RIN) was evaluated using an Agilent Bioanalyzer.

#### RNA-Sequencing Library Preparation

Library preparation and RNA sequencing were performed by Takara Bio Inc. RNA-seq libraries were prepared using the SMART-Seq v4 Ultra Low Input RNA Kit for Sequencing for full-length cDNA synthesis, followed by tagmentation and indexing with the Nextera XT DNA Library Prep Kit and IDT for Illumina Unique Dual Indexes. Sequencing was performed on an Illumina NovaSeq 6000 system (S4 flow cell, 300 cycles for 150 bp paired-end sequencing) using NovaSeq6000 S4 Reagent Kit v1.5 and NovaSeq Xp 4-Lane Kit v1.5. Base calling and demultiplexing were processed using bcl2fastq2 or the DRAGEN Bio-IT Platform to generate FASTQ files.

#### RNA-Seq Data Analysis

Nextera adapters were removed using Cutadapt (v3.4). Subsequently, RNA-seq reads were mapped to the mouse reference genome (GRCm38) using STAR software (v2.7.1a). Transcript abundance and raw counts were obtained using RSEM (v1.2.31). Differential expression analysis was performed on all 55,291 annotated genes using DESeq2 (v1.26.0) in the R environment. Pairwise Pearson correlation coefficients (*R*^2^) between biological replicates (*n* = 2) were calculated using normalized count data, and scatter plots displayed the log_2-_transformed expression levels [log_2(_TPM + 1)] of all quantified genes between the biological replicates (Figure S5N). TPM values were calculated for a curated set of 21,859 genes; genes outside this set are indicated as “N/A” in the Data S6. Genes meeting the criteria of *P*_adj <_ 0.05, |log_2(_Fold Change)| ≥ 1, baseMean ≥ 31, and mean TPM ≥ 1 (in either condition) were classified as differentially expressed genes (DEGs). Additionally, a subset of highly expressed and significantly regulated genes was defined by applying a stricter threshold: *P*_adj <_ 0.05, |log_2(_Fold Change)| > 2, and baseMean > 1000. A volcano plot displayed all quantified DEGs, with highly expressed and significantly regulated genes highlighted (Figure 4N; Table S3).

### RNA Fluorescence In Situ Hybridization (RNA FISH)

RNA FISH was performed on both primary mouse splenic B cells and cultured CH12 cells. Primary B cells were isolated from WT mouse splenocytes using the MojoSort Mouse Pan B Cell Isolation Kit. Unstimulated cells were harvested immediately as day 0 controls, whereas remaining cells were cultured in the presence of LPS and IL-4 for 4 days to induce CSR to IgG1.

Custom tiling oligonucleotide probes targeting the 5′ region of the *Sμ*-core, 3′ region of the *Sα*-core, and 3′ region of the *Sγ1*-core were designed using Stellaris FISH Probe Designer (sequences listed in Data S1). For the hybridization assay, 1.2 × 10^6^ cells were seeded onto 18 mm round poly-L-lysine-coated coverslips placed in 12-well plates and centrifuged at 1200 rpm for 5 min to facilitate adherence. Cells were then fixed and hybridized with Quasar 570 (Q570)- or Quasar 670 (Q670)-labeled probe sets according to manufacturer’s protocol for adherent cells.

Images were acquired using an Evident FLUOVIEW FV4000 confocal microscope (*z*-step size, 0.39 μm) and are presented as maximum intensity projections of *z*-stacks alongside single *z*-slices. Nuclei were stained with Hoechst 33342. Scale bars, 5 μm. RNA FISH signals of *Sμ*, *Sα,* and *Sγ1* foci were counted based on maximum intensity projections of *z*-stacks and verified using individual *z*-slices.

### Expression of Coilin-mCardinal

Human *COIL* cDNA was amplified from the cDNA pool derived from HeLa cells and initially subcloned into the pCR4Blunt-TOPO vector using the Zero Blunt TOPO PCR Cloning Kit for Sequencing. The pCMV-*COIL*-EGFP construct was generated using pEGFP-N1 as backbone vector. Subsequently, the EGFP sequence was replaced with mCardinal^124^ using GeneArt Gibson Assembly HiFi Master Mix. All primers are listed in Data S1. The plasmids were transiently transfected into CH12 cells for RNA FISH analysis.

### Generation of *Sirius-8xMS2* Knock-In CH12 Cell Lines

#### Construction of Targeting Vector and Cas9/sgRNA Plasmids

A *Sirius-8xMS2* sequence^125^ was inserted into the *Sα*-core region of the *IgH* locus in CH12 (CH12F3-2A)-Bcl2-Neo cells and CH12-dCas9-KRAB (#40) cells.^66^ The insertion strategy was similar to that used for 8xLexA insertion,^28^ with minor modifications (Figure S7). Briefly, the *Sirius-8xMS2* sequence derived from pPUR-hU6-sgRNA-*Sirius-8xMS2*^125^ was cloned into the targeting vector in place of the 8xLexA sequence using GeneArt Gibson Assembly HiFi Master Mix, generating pDT-A/*Sirius-8xMS2*-*Sα*-core-EGFP-Puro (Figures S7A and S7B). Additional pX335-U6-S*α*-#11_Chimeric_BB-CBh-hSpCas9n(D10A) vector was generated. Targeting vector was co-electroporated with two pX335 vectors encoding *Sα*-core-targeting sgRNAs (pX335-U6-S*α*-#11_Chimeric_BB-CBh-hSpCas9n(D10A) and pX335-U6-S*α*-#21_Chimeric_BB-CBh-hSpCas9n(D10A)). Stable clones were selected as described for the 8xLexA insertion, and successful *Sirius-8xMS2* integration was verified by PCR (Figure S7C). Clones #6, #8, #11, and #16 were selected based on their preserved IgA switching efficiency (IgA%) (Figure S7D). To remove selection cassette, the PGK-GFP-IRES-Puro cassette was excised by transient Cre recombinase expression, and GFP-negative cells were isolated using a BD FACSMelody Cell Sorter (Figure S7E). Clone #11 was selected for subsequent experiments and designated CH12-*Sirius-8xMS2*-*Sα*-core (#11).

To visualize and immunoprecipitate *Sα*-GLTs, the expression vector, pCMV-3xFLAG-tdMCP-EGFP, which encodes the MS2-binding protein tdMCP fused to a 3xFLAG tag, nuclear localization signal (NLS), and EGFP was constructed in two steps. First, mCherry was replaced with EGFP from the pHR-TRE3G-NLS-tdPCP-EGFP vector^126^ in the pHR-TRE3G-NLS-tdMCP-mCherry vector^126^ using GeneArt Gibson Assembly HiFi Cloning Kit to generate pHR-TRE3G-NLS-tdMCP-EGFP. Second, the tdMCP-EGFP cassette was inserted into the pCMV-3xFLAG-FBL vector backbone, and PCR fragments from both vectors were assembled via Gibson Assembly to generate the final pCMV-3xFLAG-tdMCP-EGFP construct.

The tdMCP expression vector was stably introduced into clone #11 (Figure S7F), and GFP^+^ cells were isolated by flow cytometry sorting, followed by limiting dilution cloning. Among single-cell clones, clone #64 was specifically chosen because it exhibited successful cassette deletion (Figure S7G), acceptable IgA switching efficiency upon CIT stimulation (Figure S7H), and minimal GFP and tdMCP expression (Figure S7I), thereby minimizing nonspecific immunoprecipitation. The insertion of Sirius-8xMS2 sequence in clone #64 was verified (Figure S7J). This clone was designated CH12-*Sirius-8xMS2*-*Sα*-core-3xF-tdMCP-EGFP (#64). RNA immunoprecipitation (RIP) using an anti-FLAG antibody verified the selective enrichment of Sα germline transcripts (GLTs) in this clone (Figure S7K). As negative control (referred to as “Mock cells” in Figure 6), the pCMV-3xFLAG-NLS-EGFP vector lacking tdMCP was stably introduced into clone #11, generating CH12-*Sirius-8xMS2*-*Sα*-core-3xFLAG-EGFP cells.

The insertion of the *Sirius-8xMS2* sequence into CH12-dCas9-KRAB (#40) cells followed identical steps used to generate clone #11. The resulting clone, featuring the *Sirius-8xMS2* insertion at the *Sα-*core and successful deletion of the PGK-EGFP-IRES-Puro cassette, was designated CH12-dCas9-KRAB-*Sirius-8xMS2*-*Sα*-core (#451). This clone was then stably transfected with pCMV-3xFLAG-tdMCP-EGFP to generate the CH12-dCas9-KRAB-*Sirius-8xMS2*-*Sα*-core-3xFLAG-tdMCP-EGFP (#95) cell line. Similarly, stable transfection of pCMV-3xFLAG-NLS-EGFP into clone #451 yielded the CH12-dCas9-KRAB-*Sirius-8xMS2*-*Sα*-core-3xFLAG-EGFP control (“Mock”) cell line.

### *Sirius-8xMS2*-Mediated Ribonucleoprotein Isolation

#### Conjugation of Anti-FLAG Antibody to Tosyl-Activated Beads

A volume of 165 μL of tosyl-activated beads was washed three times with Buffer B (sodium phosphate buffer, pH 7.4). Beads were then resuspended in 150 μL of Buffer B containing 100 μg of glycerol-free anti-FLAG M2 antibody. Following the addition of 100 μL of Buffer C (pH 7.4 sodium phosphate buffer containing 3 M ammonium sulfate), the mixture was incubated on a rotator for 18 h at 37°C. Tubes were placed on a magnet stand to discard supernatant. Beads were subsequently blocked by adding 1 mL of Buffer D (0.5% BSA in PBS) and incubated on a rotator for 1 h at 37°C. After washing with 1 mL of Buffer E (0.1% BSA in PBS), Conjugated beads were resuspended in 240 μL of Buffer E, supplemented with 0.05% sodium azide (1.2 μL of 10% NaN_3)_, and stored at 4°C until use.

#### Sirius-8xMS2-Mediated Isolation of Ribonucleoproteins

For each sample, 20 μL of tosyl-activated beads conjugated with anti-FLAG antibody were washed with Wash Buffer (50 mM HEPES-KOH, pH 7.5, 100 mM KCl, 0.5 mM EDTA, 0.05% NP-40) and resuspended in 20 μL of same buffer. To isolate endogenous RNA-protein complexes under native conditions, 50 μL of cell pellets (without prior chemical crosslinking) were lysed in 150 μL of NP-40 Mild Lysis Buffer (50 mM HEPES-KOH, pH 7.5, 100 mM KCl, 0.5% IGEPAL CA-630, 0.01% sodium deoxycholate, 10% glycerol) supplemented with 1 mM DTT, 1.5 μL RNaseOUT, and 1x cOmplete Protease Inhibitor Cocktail. Following complete cell lysis, 0.6 μL of DNase I (50 Kunitz U/μL) and 1 μL of 1 M MgCl_2 w_ere added, and lysates were incubated on ice for 30 min. Lysates were cleared by centrifugation at 20,000 × *g* for 15 min at 4°C. Then, 160 μL of cleared lysates were mixed with 640 μL of Dilution Buffer (50 mM HEPES-KOH, pH 7.5, 100 mM KCl, 0.5 mM EDTA) and 20 μL of pre-washed beads. The mixture was incubated on a rotator at 4°C for 2 h.

Beads were subsequently washed six times with Wash Buffer (each wash included a 3 min incubation on a rotator at 4°C). To verify pull-down efficiency, 10% of the bead suspension was aliquoted into 500 μL of TRIzol Reagent for RNA isolation. Isolated RNA was reverse-transcribed using SuperScript IV VILO Master Mix and analyzed by qPCR.

To specifically elute RNA-scaffolded protein complexes, remaining 90% of the beads were resuspended in 20 μL of RNase Buffer (50 mM HEPES-KOH, pH 7.5, 75 mM KCl, 3 mM MgCl_2,_ 1 mM DTT) containing 0.5 μL RNase A, 0.5 μL ShortCut RNase III, and 0.4 μL 1x cOmplete Protease Inhibitor Cocktail. This mixture was incubated for 15 min at 37°C. Supernatant containing specifically eluted proteins was transferred to a new tube. To maximize protein recovery, remaining beads were rinsed twice with 10 μL of RNase Buffer, and these washes were pooled with initial 20 μL eluate. Recovered protein samples were snap-frozen at −80°C.

### Mass Spectrometry Analysis of tdMCP-Bound Proteins

#### SP3-Based Sample Preparation and Peptide Digestion

Proteins were reduced with 5 mM DTT for 30 min at RT and alkylated with 27.5 mM iodoacetamide for 30 min in the dark. Samples were subjected to single-pot solid-phase-enhanced sample preparation (SP3) using Sera-Mag SpeedBead carboxylate-modified magnetic particles (Cytiva). Beads were washed with 80% ethanol and acetonitrile and resuspended in 50 mM triethylammonium bicarbonate. Proteins bound to beads were digested with 400 ng of trypsin/Lys-C mix (Promega) overnight at 37°C. Resulting peptides were acidified with TFA, desalted using GL-Tip SDB (GL Sciences), concentrated using a SpeedVac, and reconstituted in 3% ACN/0.1% TFA.

#### LC-MS/MS Analysis

Peptides were analyzed using a nanoElute 2 system coupled to a timsTOF HT mass spectrometer (Bruker) and separated on a 75 μm inner diameter × 150 mm C18 reversed-phase column (NTCC-360/75-3-155; Nikkyo Technos). LC gradient was 5%–20% ACN from 0 to 40 min, 20%–35% ACN from 40 to 60 min, followed by an increase to 95% ACN from 60 to 61 min, and a hold at 95% ACN until 75 min. Data were acquired in DIA-PASEF mode with an MS1 *m*/*z* range of 100–1700 and an ion mobility (1/K_0)_ range of 0.6–1.6. Ramp time was 100 ms, with a 100% duty cycle. MS2 polygon was defined by the following four vertices: (*m*/*z* = 300, 1/*K*_0 =_ 0.64), (*m*/*z* = 300, 1/*K*_0 =_ 0.80), (*m*/*z* = 770, 1/*K*_0 =_ 0.97), and (*m*/*z* = 770, 1/*K*_0_ = 1.15). An 8 Da isolation window with a 0.5 Da overlap was used, resulting in 62 windows per cycle and a cycle time of 3.36 s.

#### DIA Data Analysis

Protein identification and quantification were performed using DIA-NN (version 1.9)^127^ in library-free mode against the *Mus musculus* UniProt reference proteome (UP000000589). Parameters for *in silico* library generation included trypsin as digestion enzyme, one allowed missed cleavage, peptide lengths of 7–45 amino acids, precursor charge states of 2–4, and a fragment ion m/z range of 200–1800. Deep learning-based retention time and ion mobility predictions, N-terminal methionine cleavage, and cysteine carbamidomethylation were enabled. MS1 and MS2 mass accuracies were determined automatically, neural network classifiers were configured for single-pass mode, and quantification strategy was set to QuantUMS (high precision).

### Proteomics Data Analysis for *Sirius-8xMS2*-Mediated Interactome Mass Spectrometry

For analysis of mass spectrometry (MS) data regarding *Sirius-8xMS2*-mediated *Sα*-GLT-binding proteins, protein intensity values were processed using Perseus software (v3.0). Raw intensity values were log_2-_transformed, and missing values were imputed from a normal distribution. A two-sided Welch’s *t*-test with Benjamini-Hochberg false discovery rate (FDR) correction was performed (FDR *q*-value < 0.05 for Control vs. Mock [GFP alone] and *q* < 0.25 for si*Fbl* vs. Control).

For baseline comparative analysis (Control vs. Mock), 1813 proteins were quantified using Welch’s *t*-test. Volcano plots were generated by plotting log_2 D_ifference against −log_10(_*P*-value), with previously reported GLT-binding proteins highlighted (Figure 6E). Applying stringent thresholds of FDR *q*-value < 0.05 and log_2 D_ifference > 2 yielded 1176 significantly enriched proteins. Among these, functional modules related to GLT processing, R-loop processing, and DNA recombination were curated and visualized as a heatmap (Figure 6F; Data S8).

Gene Set Enrichment Analysis (GSEA) for baseline enriched proteins (Control vs. Mock) was performed against the Gene Ontology Biological Process (GOBP) gene set (m5.go.v2025.1.Mm.symbols.gmt). From an initial identification of 105 GOBP terms, significant pathways were filtered using thresholds of NES > 1.5 and FDR *q*-value < 0.1, yielding 53 refined GOBP terms. These 53 terms were categorized into five major functional categories for GSEA summary (Figure 6G; Data S9).

For analyzing si*Fbl*-induced changes (si*Fbl* vs. Control), volcano plots were generated following Welch’s *t*-test with FDR correction, in a manner similar to Control vs. Mock analysis (Figure 6H; Data S10), with candidate proteins regulated by si*Fbl* (e.g., WDR74 and NOL7) highlighted.

GSEA was conducted against the GO Cellular Component (GOCC) gene set (m5.go.cc.v2025.1.Mm.symbols.gmt), while all other parameters remained identical to those described above. All 21 GOCC terms identified by GSEA were visualized as a dot plot (Figure 6I; Data S11). GSEA enrichment plots were generated for specific GOCC terms, including “Nucleolus” (Figure 6J), “U2snRNP” (Figure 6K), “Intermediate Filament” (Figure S7P), and “Eukaryotic Translation Initiation Factor 3 Complex” (Figure S7Q).

In raw mass spectrometry data files deposited in jPOST/ProteomeXchange, as well as in the associated R scripts, samples designated as siFBL, siCtrl, and GFP only correspond to si*Fbl,* Ctrl, and GFP alone (Mock) in the main text and figures, respectively.

### Interactome Analysis of FBL

#### Immunoprecipitation of 3xFLAG-Tagged FBL in CH12 Cells

To identify FBL-interacting proteins, CH12F3-2A-Bcl2-Neo cells stably expressing 3xFLAG-tagged WT human FBL (CH12F3-2A-Bcl2-Neo-FBL-WT) were electroporated with si*Fbl*#1 to deplete endogenous mouse *Fbl* and cultured for 24 h. Cells were subsequently cultured for an additional 24 h, with or without CIT stimulation. CIT-stimulated and non-stimulated CH12F3-2A-Bcl2-Neo-FBL-WT and CIT-stimulated WT CH12F3-2A cells (Mock) were used for interactome analysis (*n* = 2). Crosslinking-based immunoprecipitation was performed using anti-FLAG magnetic beads, according to a previously published protocol ^128^ with minor modifications.

Briefly, a total of 8 × 10^6^ cells were fixed by adding formaldehyde to 20 mL cell culture medium to a final concentration of 0.1% for 10 min at RT. After quenching fixation by adding glycine to a final concentration of 100 mM, cells were incubated for an additional 4 min at RT. Cells were washed twice with 5 mL ice-cold HBS Buffer (20 mM HEPES-NaOH, pH 7.5, 150 mM NaCl) and lysed in 1 mL of HEPES-RIPA Buffer (20 mM HEPES-NaOH, pH 7.5, 150 mM NaCl, 1 mM MgCl_2,_ 1% NP-40, 1 mM EGTA, 0.05% SDS, and 0.25% sodium deoxycholate) supplemented with Benzonase (2 μL), 1x cOmplete Protease Inhibitor Cocktail, and 1x PhosSTOP. Lysates were sonicated using a Bioruptor II (5 cycles of 30 s ON / 60 s OFF). Following centrifugation at 20,000 × *g* for 15 min at 4°C, cleared supernatant was transferred to a new tube.

For each sample, 3.5 μL of anti-FLAG magnetic beads were pre-washed three times with Antibody Binding Buffer (20 mM HEPES-NaOH, pH 7.5, 5% glycerol, 150 mM NaCl, 1 mM MgCl_2,_ 1% NP-40, and 1 mM EGTA). Cleared lysate was mixed with pre-washed beads and incubated on a rotator for 2 h at 4°C. Beads were then washed five times with 1 mL of HEPES-RIPA Buffer and finally resuspended in 50 μL of 50 mM ammonium bicarbonate. Proteins bound to beads were digested with 200 ng of trypsin/Lys-C mix overnight at 37°C with shaking. Peptides were reduced, alkylated, acidified, and desalted using GL-Tip SDB. Eluates were concentrated using a SpeedVac and dissolved in 3% ACN/0.1% TFA.

#### Mass Spectrometry of FBL Interactome

Peptides were analyzed using an EASY-nLC 1200 system coupled to an Orbitrap Fusion mass spectrometer (Thermo Fisher Scientific). Peptides were separated on a 15 cm C18 capillary column at a flow rate of 300 nL/min. LC gradient was increased from 4% to 32% ACN over 100 min, followed by an increase to 80% ACN over 10 min and a 10 min hold at 80% ACN. Orbitrap Fusion was operated in data-dependent acquisition mode with a cycle time of 3 s. Full MS spectra were acquired in the Orbitrap over an *m/z* range of 350–1500 at a resolution of 60,000, with an AGC target of 4 × 10^5^ and a maximum injection time of 50 ms. Dynamic exclusion was set to 15 s. Precursor ions were isolated using a quadrupole isolation window of 1.6 *m/z* and fragmented by higher-energy collisional dissociation (HCD) at a normalized collision energy of 30%. MS/MS spectra were acquired in Orbitrap at a resolution of 30,000, with an AGC target of 5 × 10^4^ and a maximum injection time of 54 ms.

Raw mass spectrometry data were processed using Proteome Discoverer version (v2.5; Thermo Fisher Scientific). Tandem mass spectra were searched against the UniProt *Mus musculus* protein sequence database supplemented with the sequence of 3xFLAG-tagged human FBL. Precursor and fragment mass tolerances were set to 10 ppm and 0.02 Da, respectively. Carbamidomethylation of cysteine was specified as a fixed modification. Oxidation of methionine; methylation of lysine, glutamine, and arginine; and protein N-terminal acetylation were specified as variable modifications. Peptide identifications were filtered at an FDR of 1% using Percolator.^129^ Label-free quantification was performed based on the precursor ion intensities. Peptide abundances were normalized by total peptide amount across all samples. Data S12 was used to evaluate the interaction levels of the candidate proteins WDR74, LSM7, and NOL7 with FBL.

### Kaplan-Meier Survival Analysis

Prognostic value of *WDR74*, *LSM7*, and *NOL7* mRNA expression was evaluated using the Kaplan-Meier Plotter dataset for myeloma (*n* = 1416) and breast cancer (*n* = 4929) cohorts.^89^ Optimal Jetset probes^130^ were selected to ensure specificity and sensitivity. Survival curves were compared using log-rank test. For survival analyses in the Kaplan-Meier Plotter database, patients were split into high and low expression groups based on either the “Auto select best cutoff” criteria (for myeloma cohort) or median expression level (for breast cancer cohort).

### Overexpression of WDR74, LSM7 and NOL7 with or without tdMCP Fusion in CH12 Cells

Coding fragments of *WDR74*, *LSM7*, and *NOL7* were amplified from cDNA generated from HeLa cells. Expression vectors pCMV-WDR74-3xFLAG-P2A-Puro, pEF-tdMCP-WDR74-3xFLAG-P2A-Puro, pCMV-LSM7-3xFLAG-P2A-Puro, pCMV-tdMCP-LSM7-3xFLAG-P2A-Puro, pEF-NOL7-3xFLAG-P2A-Puro, pEF-tdMCP-NOL7-3xFLAG-P2A-Puro, and pCMV-tdMCP-MOCK-3xFLAG-P2A-Puro were generated using primers listed in Data S1. Vector backbones were derived from pCMV6-mAID-3xFLAG vector and pEF1α-IRES-ZsGreen1 vector, and P2A-Puro fragment was derived from pLenti-FNLS-P2A-Puro.^131^ Resulting expression plasmids were linearized by digestion with AseI. A total of 1.2 μg of each linearized plasmid was electroporated into CH12-dCas9-KRAB-*Sirius-8xMS2*-*Sα*-core (#451) cells, followed by selection with puromycin for 10 days to establish mass stable cell lines. These stable transfectants were then stimulated with CIT, and efficiency of CSR to IgA was evaluated 24 h post-stimulation by flow cytometry.

## QUANTIFICATION AND STATISTICAL ANALYSIS

### General Statistical Framework

Data are presented as the mean ± SEM unless otherwise specified. For RT-qPCR assays, technical replicates with C_T v_alues differing by > 0.5 were excluded as outliers prior to calculating mean values. For experiments involving technical replicates, data are presented simply to display mean values, with individual technical replicates shown as points without formal statistical hypothesis testing. All statistical analyses and graph generation were performed using GraphPad Prism (v9.1.0 and v11.1.0) and R (v4.4.3). Exact statistical tests applied, along with the corresponding *n* values (representing independent biological replicates) and exact *P values* or adjusted *P* values where applicable, are detailed in respective figure legends. In figure legends, *P* values adjusted for multiple testing (e.g., Dunnett’s test or BKY FDR procedure) are uniformly denoted as *P_adj_*. Unless otherwise specified, a *P value* or adjusted *P* values of < 0.05 was considered statistically significant.

### Statistical methodologies were chosen based on experimental design and data structure

1. **Comparison of three or more matched experimental groups:** To compare three or more matched experimental groups across independent biological replicates, raw data were log_2-_transformed to account for multiplicative variations and inter-experimental batch effects. Transformed data were subsequently analyzed using a repeated measures (RM) one-way ANOVA (or a mixed-effects model fitted via restricted maximum likelihood [REML] for datasets with missing values) followed by Dunnett’s multiple comparisons test against control group. When fitting mixed-effects models, if estimated random-effect (batch) variance was zero or negative, the random factor was removed to avoid model over-parameterization, falling back to ordinary one-way ANOVA.
2. **Comparison of two matched experimental groups:** To compare two matched experimental groups subjected to similar batch-to-batch baseline fluctuations, two-tailed ratio-paired Student’s *t*-test was utilized.
3. **Normalized quantitative data:** For quantitative data (e.g., RT-qPCR or geometric mean fluorescence intensity by flow cytometry) normalized to a theoretical control value of 1.0, the data were log_2-_transformed prior to statistical testing and evaluated using a two-tailed one-sample *t*-test against a theoretical mean of 0 [log_2(_1.0) = 0], with FDR adjustment for multiple comparisons using the Benjamini–Krieger–Yekutieli method (*Q* = 5%) relative to control or WT group.
4. **Proteomic data quantification and differential expression analysis:** For LexA-mediated *Sα*-core iChIP proteomic data (Figure 1), mean Heavy/Light enrichment ratio across independent replicates was used for ranking, and functional profiling was performed in R using the clusterProfiler package (v4.14.6). To filter proteins for enrichGO analysis, an FDR *q*-value < 0.05 threshold was applied (Figures 1D and 1E). For GSEA, an FDR *q*-value < 0.25 was considered statistically significant (Figures 1F and 1G). For *Sirius-8xMS2*-mediated *Sα*-GLT-RIP proteomic data (Figure 6), raw protein intensities were processed using Perseus software (v3.0), in which log_2-_transformed abundances were evaluated using a two-tailed Welch’s *t*-test with Benjamini-Hochberg FDR correction. Log_2-_transformed values were used for GSEA with clusterProfiler. In baseline *Sα*-proteome analysis comparing control versus mock, proteins were filtered based on Welch’s *t*-test results using criteria of FDR *q*-value < 0.05 and log_2 D_ifference > 2 (Figure 6F). For corresponding GSEA analysis, GO terms with NES > 1.5 and FDR *q*-value < 0.1 were selected (Figure 6G). In si*Fbl*-induced proteome change, FDR *q*-value < 0.25 was applied to both Welch’s *t*-test and GSEA analysis (Figures 6H-6K and S7P–S7Q).
5. **Contingency table analyses (2 × 2):** For all 2 × 2 contingency table analyses (including *IgH*/*c-Myc* translocation frequencies and somatic hypermutation rates), statistical significance was determined using a two-sided Fisher’s exact test or Chi-square test with Yates’ correction. To account for multiple testing within each assay, raw *P* values were adjusted by controlling FDR at 5% using the two-stage linear step-up procedure of Benjamini-Krieger-Yekutieli method (*Q* = 5%). Adjusted *P* values (*Q* values) < 0.05 were considered statistically significant.
6. **Distribution pattern and microhomology analyses:** To assess overall distribution shifts across multiple categories (e.g., junctional microhomology profile analysis and RNA FISH distribution patterns), statistical significance was determined using a Chi-square test on raw junction numbers or cell counts. Where pairwise comparisons were necessary in RNA FISH, individual patterns were evaluated using Fisher’s exact test with FDR adjustment for multiple comparisons using the two-stage linear step-up procedure of Benjamini-Krieger-Yekutieli method (*Q* = 5%).

Negative controls (e.g., normal IgG in immunoprecipitation or unlabeled controls) were presented to confirm assay background levels and were excluded from statistical hypothesis testing and multiple comparison models.

