## Supplemental Figures and Tables for "2′-*O*-Methylated ncRNA condensates form a recombination focus for *IgH* synapsis"

[1] Department of Immunology and Genomic Medicine, Center for Cancer Immunotherapy and Immunobiology, Graduate School of Medicine, Kyoto University, Kyoto 606-8501, Japan
[2] Department of Biochemistry and Genome Biology, Hirosaki University Graduate School of Medicine, Hirosaki, Aomori 036-8562, Japan
[3] Graduate School of Science and Technology, Shizuoka University, Shizuoka 422-8529, Japan
[4] Research Institute of Green Science and Technology, Shizuoka University, Shizuoka 422-8529, Japan
[5] Graduate School of Science and Engineering, Kindai University, Higashiosaka, Osaka 577-8502, Japan
[6] Informatics Platform, Center for Cancer Immunotherapy and Immunobiology, Graduate School of Medicine, Kyoto University, Kyoto 606-8501, Japan
[7] Faculty of Science and Engineering, Kindai University, Higashiosaka, Osaka 577-8502, Japan
[8] Division of Cell Signaling, Institute of Advanced Medical Sciences, Tokushima University, Tokushima 770-8503, Japan
[9] Laboratory of Proteomics, Institute of Photonics and Human Health Frontier, Tokushima University, Tokushima 770-8503, Japan
[10] Present address: The Wistar Institute, Philadelphia, PA 19104, USA
[11] Present address: Department of Biochemistry, Faculty of Life Sciences, Aligarh Muslim University, Aligarh, Uttar Pradesh 202002, India
[12] Present address: Tritium Research Center, Institute for Environmental Sciences, Rokkasho, Aomori 039-3212, Japan
[13] Present address: The Specialty Discovery Pharmacology Department, Chugai Pharmaceutical Co., Ltd., Tokyo, Japan
[14] Present address: Guangdong Key Laboratory for Biomedical Measurements and Ultrasound Imaging, National-Regional Key Technology Engineering Laboratory for Medical Ultrasound, School of Biomedical Engineering, Shenzhen University Medical School, Shenzhen 518060, China.
[15] Present address: Frontier Institute for Biomolecular Engineering Research, Konan University, Kobe, Hyogo 650-0047, Japan
* lead contact and correspondence:
Maki Kobayashi

#### This PDF file includes:

Figures S1–S8
Tables S1–S4

The following supplemental materials referenced in the text will be made available upon formal publication:

**[Supplemental Datasets]**

Data_S1_Oligonucleotides_Used_in_This_Study.xlsx
Data_S2_LexA_SaCore_iChIP_Proteome_Set1-Set2-Combined_Related-to-Fig1C.xlsx
Data_S3_LexA_SaCore_iChIP_Proteome_enrichGO_Related-to-Fig1D-1E.xlsx
Data_S4_LexA_SaCore_iChIP_Proteome_GSEA_Related-to-Fig1F-1G.xlsx
Data_S5_FBL_Junction-Seq_Related-to-FigS3H–I.xlsx
Data_S6_Med19-DEG_Related-to-Fig4N.xlsx
Data_S7_MS2_SaCore_RIP_Proteome_Unprocessed_MS-signals_Related-to-Fig6.xlsx
Data_S8_MS2_SaCore_RIP_Proteome_Baseline_t-test_Related-to-Fig6E–F.xlsx
Data_S9_MS2_SaCore_RIP_Proteome_Baseline_GSEA_Related-to-Fig6G.xlsx
Data_S10_MS2_SaCore_RIP_Proteome_siFbl-induced_t-test_Related-to-Fig6H.xlsx
Data_S11_MS2_SaCore_RIP_Proteome_siFbl-induced_GSEA_Related-to-Fig6I-6K.xlsx
Data_S12_FBL-Interactome_Related-to-Fig6H.xlsx

**[Source Data]**

Source_Data_Gels_Blots.pdf
Source_Data_Numerical_Figure1-S1-S2.xlsx
Source_Data_Numerical_Figure2-S3.xlsx
Source_Data_Numerical_Figure3-S4.xlsx
Source_Data_Numerical_Figure4-S5.xlsx
Source_Data_Numerical_Figure5-S6.xlsx
Source_Data_Numerical_Figure6-S7.xlsx
Source_Data_Numerical_Figure7-S8.xlsx

### Figures S1–S8

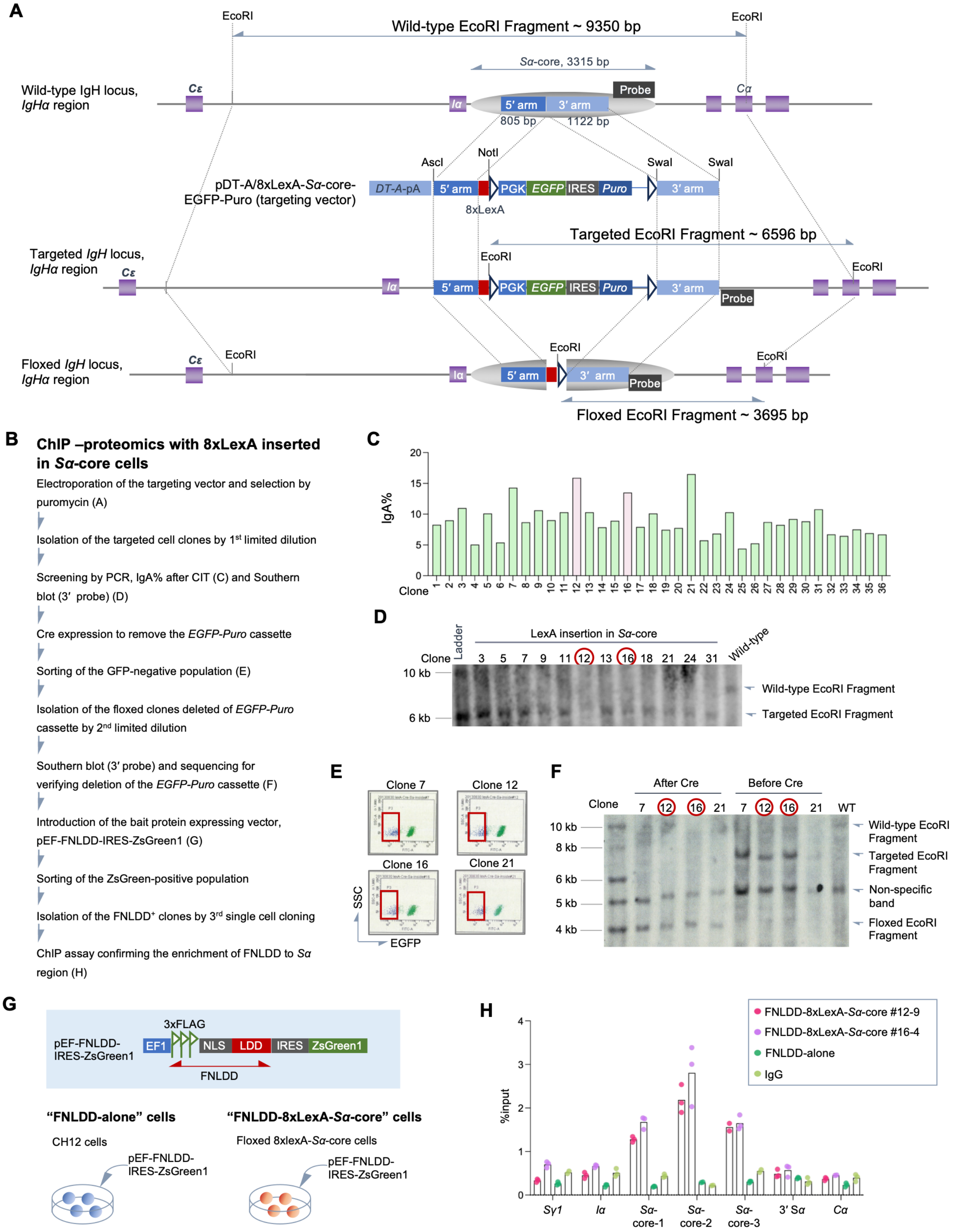

**Figure S1. Generation of 8xLexA-inserted CH12 cells at the *Sα* region of the *IgH* locus using CRISPR-Cas9, Related to Figure 1**

(A) Schematic representation of LexA sequence insertion into the *Sα* region of the *IgH* locus using a targeting vector containing an *EGFP-IRES-Puro* resistance cassette.[27,28]

(B) Experimental workflow for preparing FNLDD-8xLexA-*Sα*-core cells.

(C) IgA class switching efficiency in clones obtained from initial LexA knock-in screening (*n* = 1). Clones #12 and #16, used in subsequent experiments, are highlighted in pink. No statistical hypothesis testing was performed.

(D) Southern blot analysis of candidate clones using a DIG-labeled probe to confirm LexA insertion into the *Sα* region.

(E) Flow cytometric sorting of the EGFP-negative population following transient Cre recombinase expression in selected clones.

(F) Southern blot analysis of selected clones using a DIG-labeled probe. "After Cre" (left) and "Before Cre" (right) show the reduction in allele size from ~7 kb to ~3.7 kb upon Cre-mediated deletion of the *PGK-EGFP-IRES-Puro* cassette.

(G) Expression of 3xFLAG-tagged LexA DNA binding domain (LDD) fused to a nuclear localization signal (NLS) (FNLDD) in parental CH12 cells (FNLDD-alone) and in cells carrying floxed 8xLexA at the *Sα*-core (FNLDD-8xLexA-*Sα*-core).

(H) ChIP-qPCR analysis demonstrating specific recruitment of FNLDD to the *Sα*-core region in 8xLexA-inserted cells. Data represent the mean from a single experiment (*n* = 1 with 2-3 technical replicates), with individual values plotted as dots. No statistical hypothesis testing was performed.

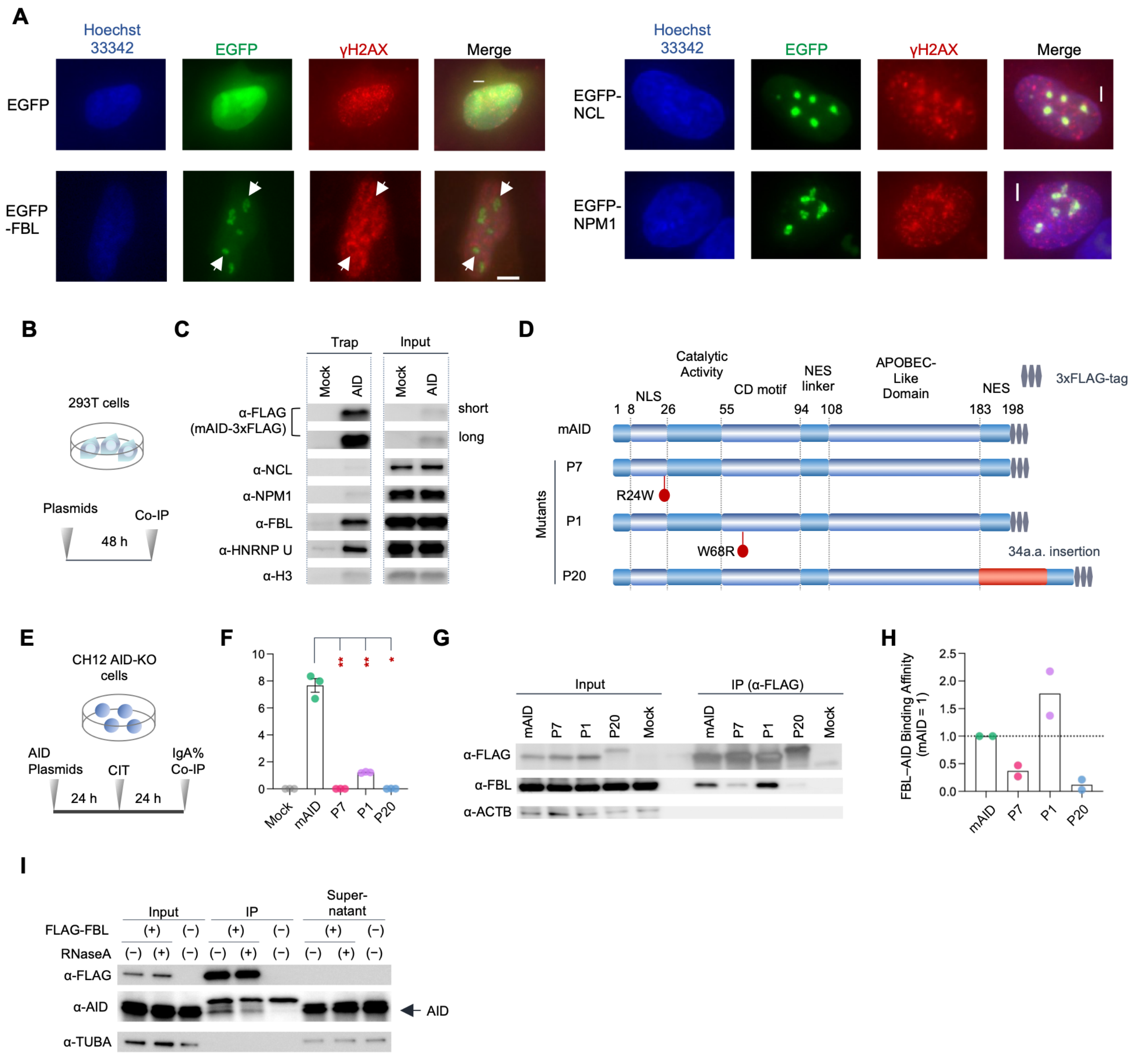


**Figure S2. Among the three nucleolus-related proteins, only FBL interacts with AID in an RNA-dependent manner, Related to Figure 1.**

(A) Immunofluorescence analysis of HeLa cells expressing EGFP alone or EGFP fused to nucleolar proteins (EGFP-FBL, EGFP-NCL, or EGFP-NPM1), treated with 4 μM etoposide for 4 h, and stained for γH2AX using an anti-mouse Alexa Fluor 555-conjugated secondary antibody is shown. Scale bars, 5 μm. EGFP, EGFP alone; EGFP-FBL, human fibrillarin fusion; EGFP-NCL, human nucleolin fusion; EGFP-NPM1, human nucleophosmin 1 fusion.

(B) Experimental timeline for co-immunoprecipitation assays.

(C) Co-immunoprecipitation of endogenous nucleolar proteins with 3xFLAG-tagged mouse AID (mAID) expressed in HEK293T cells. Mock, empty vector expressing 3xFLAG alone. Short and long exposures for anti-FLAG are shown.

(D) Schematic representation of C-terminally 3xFLAG-tagged wild-type (WT) mAID or mutant AID constructs used for rescue assays in AID-knockout (AID-KO) CH12 cells.

(E) Experimental timeline of the knockout-rescue assay using AID-KO CH12 cells.

(F) Quantification of $IgA^+$ cells in the rescued AID-KO CH12 cells 24 h post-CIT stimulation ($n$ = 3).

(G) Representative co-immunoprecipitation of 3xFLAG-tagged AID mutants with anti-FLAG antibody showing co-precipitated endogenous FBL. ACTB (β-actin) served as a loading control.

(H) Relative binding affinity of FBL to WT and mutant AID. Band intensities of co-immunoprecipitated FBL were quantified using ImageJ and normalized to immunoprecipitated AID (WT mAID was set to 1.0) ($n$ = 2). Dashed horizontal line indicates normalized control level (1.0). No statistical hypothesis testing was performed.

(I) Co-immunoprecipitation of AID with 3xFLAG-tagged FBL in the presence (+) or absence (–) of RNase A, demonstrating the RNA dependency of the AID–FBL interaction. An arrow indicates the AID-specific band. TUBA (α-tubulin) served as a loading control.

Data represent the mean ± SEM in (F) and the mean in (H). $^*P_{adj} < 0.05$, $^{**}P_{adj} < 0.01$. See STAR Methods for statistical details.

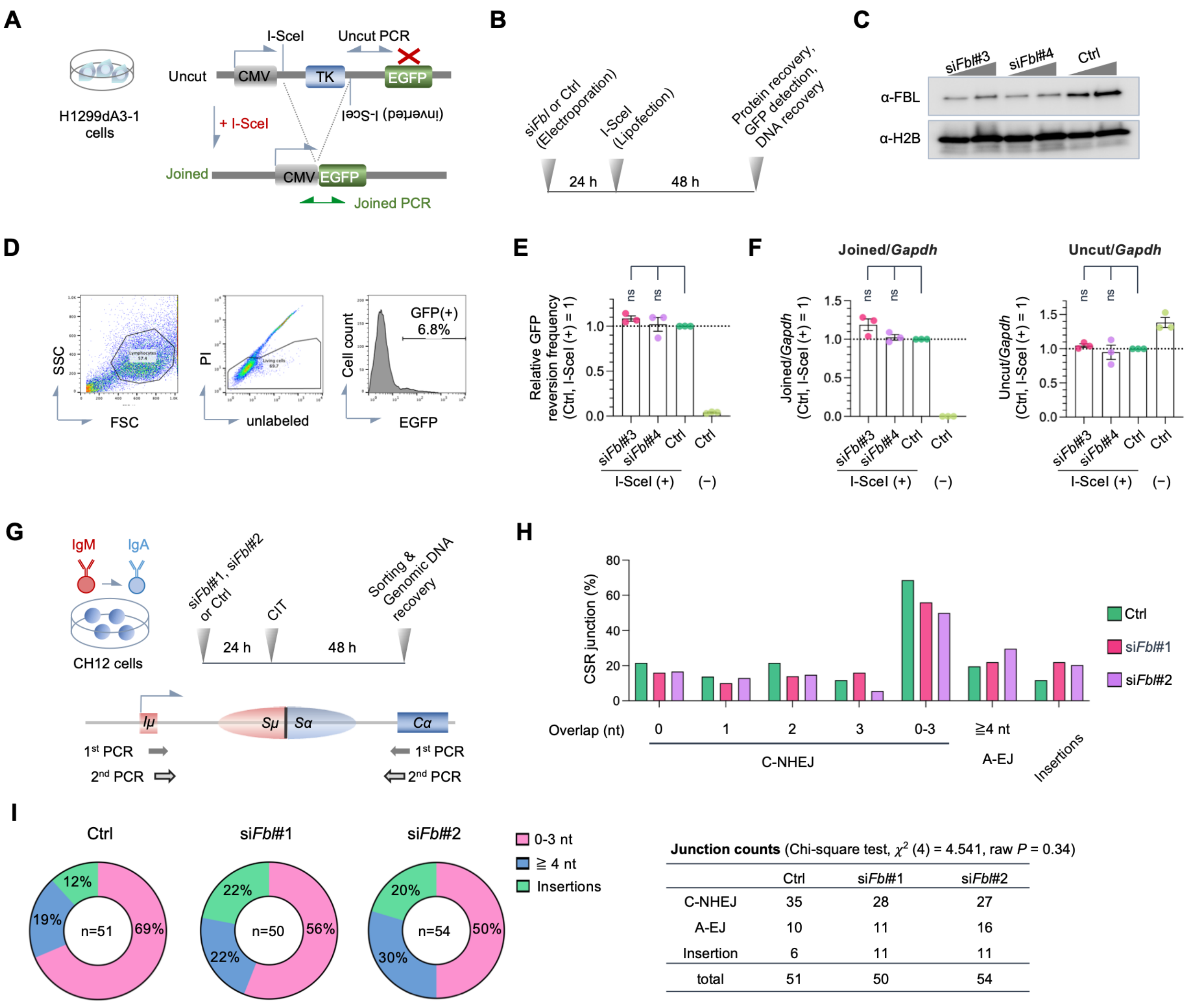


| | Ctrl | siFbl#1 | siFbl#2 |
|---|---|---|---|
| C-NHEJ | 35 | 28 | 27 |
| A-EJ | 10 | 11 | 16 |
| Insertion | 6 | 11 | 11 |
| total | 51 | 50 | 54 |

**Figure S3. *Fbl* knockdown does not affect non-homologous end-joining (NHEJ) step of CSR, Related to Figure 2**

(A) Schematic representation of an NHEJ assay based on deletion and end-processing in H1299dA3-1 cells (human lung carcinoma-derived) stably transfected with an NHEJ reporter substrate containing two inverted I-SceI cleavage sites. Following NHEJ, end-joined DNA was detected by GFP fluorescence and PCR amplification. Expected PCR amplicons for joined and uncut DNA fragments are indicated by arrows.
(B) Experimental timeline of NHEJ assay. Ctrl, non-targeting control siRNA.
(C) Western blot analysis evaluating the knockdown efficiency of si*Fbl*#3 and si*Fbl*#4, targeting human *FBL*. H2B served as a loading control.
(D) Representative flow cytometric profiles of GFP-positive cells following I-SceI expression.
(E) Quantification of GFP-positive cell frequencies (GFP reversion frequency) relative to I-SceI-treated control cells (set to 1.0) ($n$ = 3).
(F) qPCR analysis measuring joined and uncut allele frequencies normalized to *Gapdh* ($n$ = 3).

(G) Schematic strategy for *Sμ–Sα* junction microhomology analysis and position of nested primer sets at the *Iμ* (forward) and *Cα* (reverse) regions.

(H) Microhomology sequence analysis of the *Sμ–Sα* junctions in the genomic DNA of CIT-stimulated *Fbl*-knockdown cells. C-NHEJ, classical non-homologous end-joining; A-EJ, alternative end-joining.

(I) Pie charts illustrating junctional sequence characteristics at *Sμ–Sα* junctions in control and si*Fbl*-transfected cells (left) and a table of junction counts (right). Overall junction distributions showed no significant differences among groups as determined by a chi-square test. See Data S5 for statistical details.

Data represent the mean ± SEM except in (H), where values represent calculated frequencies. Dashed horizontal lines indicate the normalized control level (1.0). ns, not significant. See STAR Methods for statistical details.

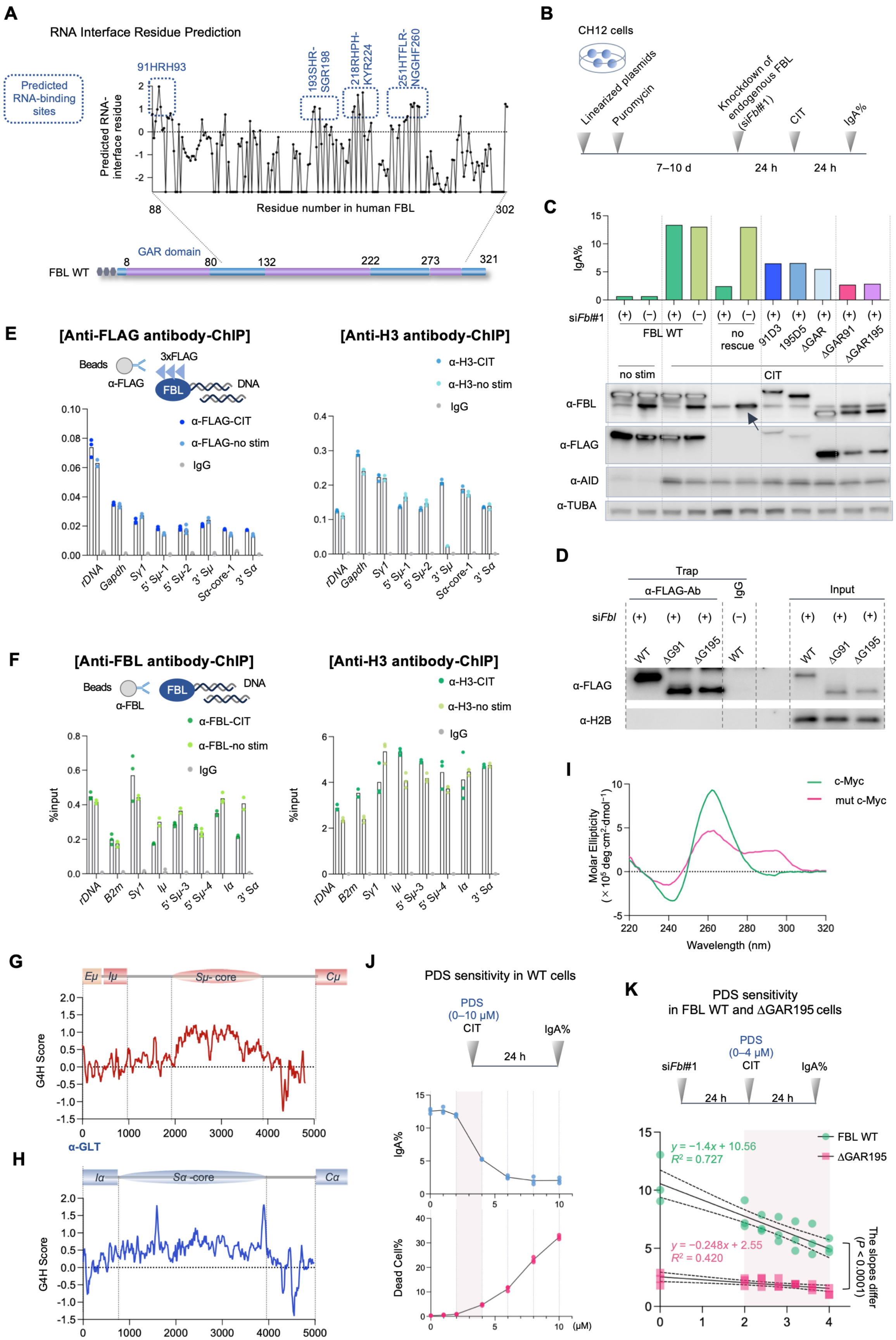
A
RNA Interface Residue Prediction
Predicted RNA-binding sites
91HRH93
193SHR-SGR198
218RHPH-KYR224
251HTFLR-NGGHF260
Predicted RNA-interface residue
Residue number in human FBL
88
302
GAR domain
FBL WT
8
80
132
222
273
321
B
CH12 cells
Linearized plasmids
Puromycin
Knockdown of endogenous FBL (siFbl#1)
CIT
IgA%
7–10 d
24 h
24 h
C
IgA%
siFbl#1
FBL WT
no rescue
91D3
195D5
ΔGAR
ΔGAR91
ΔGAR195
no stim
CIT
α-FBL
α-FLAG
α-AID
α-TUBA
D
Trap
α-FLAG-Ab
IgG
Input
siFbl
WT
ΔG91
ΔG195
α-FLAG
α-H2B
E
[Anti-FLAG antibody-ChIP]
Beads
α-FLAG
3xFLAG
FBL
DNA
α-FLAG-CIT
α-FLAG-no stim
IgG
rDNA
Gapdh
Sγ1
5′ Sμ-1
5′ Sμ-2
3′ Sμ
Sα-core-1
3′ Sα
[Anti-H3 antibody-ChIP]
α-H3-CIT
α-H3-no stim
IgG
F
[Anti-FBL antibody-ChIP]
Beads
α-FBL
FBL
DNA
α-FBL-CIT
α-FBL-no stim
IgG
%input
rDNA
B2m
Sγ1
Iμ
5′ Sμ-3
5′ Sμ-4
Iα
3′ Sα
[Anti-H3 antibody-ChIP]
α-H3-CIT
α-H3-no stim
IgG
%input
G
Eμ
Iμ
Sμ- core
Cμ
G4H Score
α-GLT
H
Iα
Sα -core
Cα
G4H Score
I
c-Myc
mut c-Myc
Molar Ellipticity (× 10⁵ deg·cm²·dmol⁻¹)
Wavelength (nm)
J
PDS sensitivity in WT cells
PDS (0–10 μM)
CIT
IgA%
24 h
IgA%
Dead Cell%
(μM)
K
PDS sensitivity in FBL WT and ΔGAR195 cells
siFbl#1
PDS (0–4 μM)
CIT
IgA%
24 h
24 h
FBL WT
ΔGAR195
y = −1.4x + 10.56
R² = 0.727
y = −0.248x + 2.55
R² = 0.420
The slopes differ (P < 0.0001)

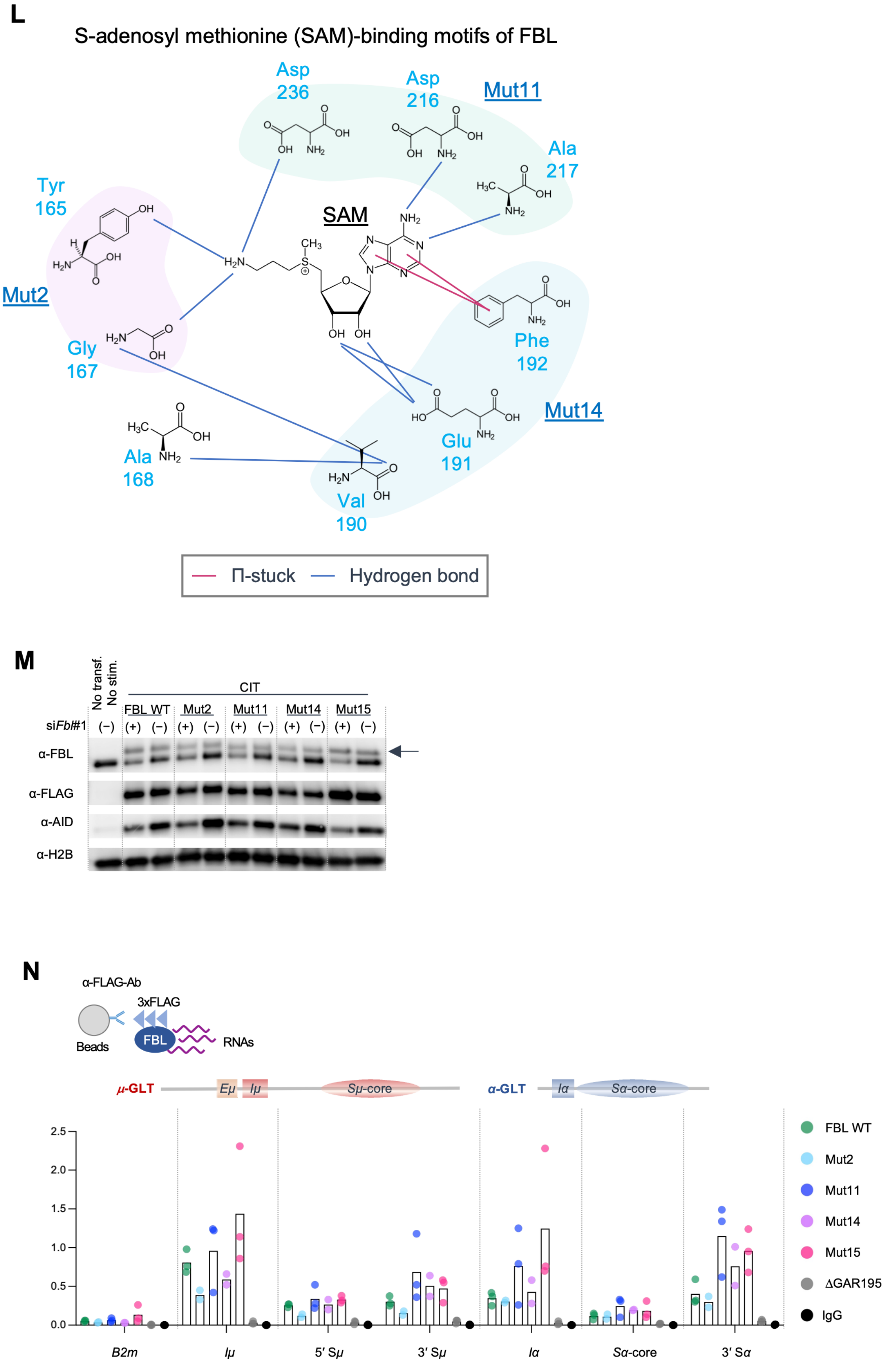
L
S-adenosyl methionine (SAM)-binding motifs of FBL
Asp 236
Asp 216
Mut11
Ala 217
Tyr 165
SAM
Mut2
Phe 192
Gly 167
Mut14
Glu 191
Ala 168
Val 190
Π-stuck
Hydrogen bond
M
No transf.
No stim.
CIT
FBL WT
Mut2
Mut11
Mut14
Mut15
siFbl#1
(−) (+) (−) (+) (−) (+) (−) (+) (−) (+) (−)
α-FBL
α-FLAG
α-AID
α-H2B
N
α-FLAG-Ab
3xFLAG
Beads
FBL
RNAs
μ-GLT
Eμ
Iμ
Sμ-core
α-GLT
Iα
Sα-core
2.5
2.0
1.5
1.0
0.5
0.0
B2m
Iμ
5′ Sμ
3′ Sμ
Iα
Sα-core
3′ Sα
FBL WT
Mut2
Mut11
Mut14
Mut15
ΔGAR195
IgG

**Figure S4. Structural and functional characterization of FBL RNA-binding and catalytic mutants, Related to Figure 3**

(A) Predicted RNA-binding sites generated using the KYG algorithm[37] based on human FBL structure (PDB ID: 2IPX).[36]

(B) Experimental timeline of knockdown-rescue experiments using stable cell lines expressing human FBL.

(C) Representative evaluation of knockdown-rescue using si*Fbl* and WT or mutant FBL: quantification of $IgA^+$ cells (top; $n = 1$) and western blot analysis of exogenous FBL variants and AID. *α*-tubulin (TUBA) served as a loading control (bottom). Expression levels of ΔGAR91 and ΔGAR195 are equivalent to endogenous FBL without si*Fbl* (lane 6, indicated by an arrow). no stim, no stimulation. No statistical hypothesis testing was performed.

(D) Representative western blot analysis evaluating anti-FLAG immunoprecipitation efficiency for RIP experiments shown in Figure 3D.

(E–F) ChIP-qPCR analysis assessing the localization of FBL to the *IgH* locus using anti-FLAG (E) or anti-FBL (F) antibody in cells overexpressing 3xFLAG-tagged WT FBL. Anti-H3 served as a positive control. $n = 1$ experiment with 2–3 technical replicates. No statistical hypothesis testing was performed.

(G–H) G-quadruplex (G4) RNA predictions calculated using G4RNA screener v0.2 (window length = 60, window step = 10), showing G4 Hunter (G4H) scores for S*μ*-GLTs (G) and S*α*-GLTs (H).

(I) Circular dichroism (CD) spectra of WT and mutant (mut) *c-Myc* G4 DNA oligonucleotides in 100 mM KCl and 50 mM Tris-HCl (pH 7.5) at a strand concentration of 8 μM. Line colors: green, *c-Myc*; magenta, mutant *c-Myc*.

(J) Inhibition of IgA switching by the G4 ligand pyridostatin hydrochloride (PDS): experimental workflow of PDS exposure (top) and quantification of $IgA^+$ cells (blue) or percentage of dead cells (magenta) across a 1–10 μM PDS range (bottom) ($n = 1$). No statistical hypothesis testing was performed.

(K) IgA switching inhibition by PDS in *Fbl*-knockdown CH12 cells rescued by WT or ΔGAR195 FBL: experimental workflow for PDS exposure (top) and quantification of $IgA^+$ cells in WT-rescued (green) or ΔGAR195-rescued (magenta) cells across a 0 to 4 μM PDS range (bottom). Slopes were compared using linear regression analysis ($P < 0.0001$; $n = 3$ independent experiments). Solid and dashed lines represent the linear regression fit and 95% confidence intervals (CI), respectively.

(L) Amino acid residues in FBL interacting with S-adenosylmethionine (SAM) and the design of the catalytic mutants identified from human FBL structures (PDB:2IPX[36] and 7SE7[42]).

(M) Representative western blot analysis showing expression levels of endogenous and exogenous FBL variants, AID, and H2B, shown in Figure 3L. H2B served as a loading control. An arrow indicates exogenous FBL mutants.

(N) RNA-immunoprecipitation (RIP)-qPCR analysis assessing binding of 3xFLAG-tagged FBL WT and catalytic mutants to germline transcripts (GLTs). $n = 3$ independent experiments ($n = 2$ for Mut2 and Mut14). No statistical hypothesis testing was performed.

Data represent the mean with individual values plotted as dots.

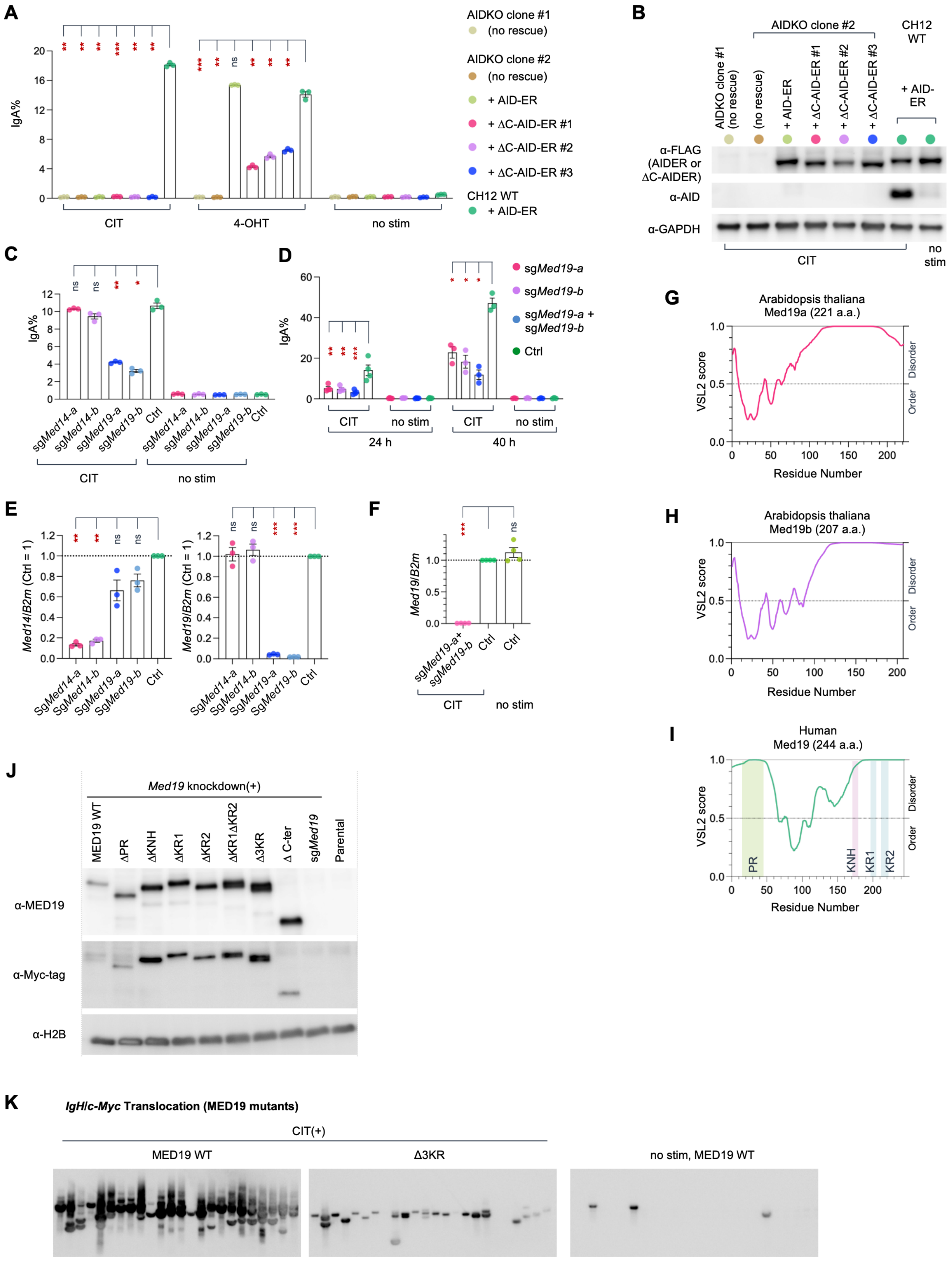
A
IgA%
CIT
4-OHT
no stim
AIDKO clone #1
(no rescue)
AIDKO clone #2
(no rescue)
+ AID-ER
+ ΔC-AID-ER #1
+ ΔC-AID-ER #2
+ ΔC-AID-ER #3
CH12 WT
+ AID-ER
B
AIDKO clone #2
CH12 WT
AIDKO clone #1 (no rescue)
(no rescue)
+ AID-ER
+ ΔC-AID-ER #1
+ ΔC-AID-ER #2
+ ΔC-AID-ER #3
+ AID-ER
α-FLAG (AIDER or ΔC-AIDER)
α-AID
α-GAPDH
CIT
no stim
C
IgA%
sgMed14-a
sgMed14-b
sgMed19-a
sgMed19-b
Ctrl
CIT
no stim
D
IgA%
sgMed19-a
sgMed19-b
sgMed19-a + sgMed19-b
Ctrl
CIT
no stim
24 h
40 h
E
Med14/B2m (Ctrl = 1)
Med19/B2m (Ctrl = 1)
SgMed14-a
SgMed14-b
SgMed19-a
SgMed19-b
Ctrl
F
Med19/B2m
sgMed19-a+ sgMed19-b
Ctrl
CIT
no stim
G
Arabidopsis thaliana
Med19a (221 a.a.)
VSL2 score
Disorder
Order
Residue Number
H
Arabidopsis thaliana
Med19b (207 a.a.)
VSL2 score
Disorder
Order
Residue Number
I
Human
Med19 (244 a.a.)
VSL2 score
PR
KNH
KR1
KR2
Disorder
Order
Residue Number
J
Med19 knockdown(+)
MED19 WT
ΔPR
ΔKNH
ΔKR1
ΔKR2
ΔKR1ΔKR2
Δ3KR
Δ C-ter
sgMed19
Parental
α-MED19
α-Myc-tag
α-H2B
K
IgH/c-Myc Translocation (MED19 mutants)
CIT(+)
MED19 WT
Δ3KR
no stim, MED19 WT

**L**

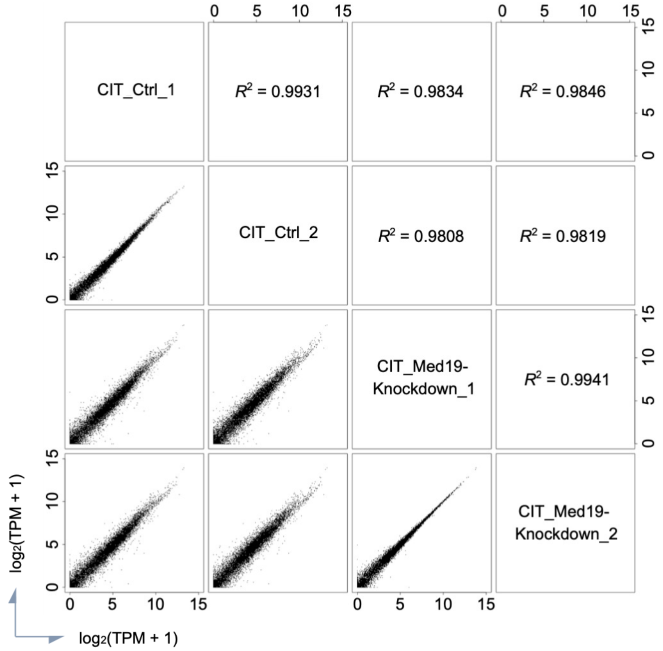


**M**

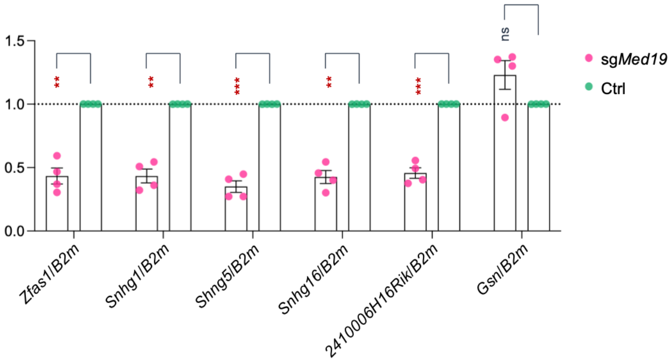


**Figure S5. MED19 regulates CSR and *IgH*/*c-Myc* translocation, Related to Figure 4**

(A) Quantification of IgA$^+$ cells of AID-knockout (AID-KO) or AID WT (AID-WT) cells stimulated with CIT or 4-OHT for 24 h, stably expressing 3xFLAG-tagged AID-estrogen receptor ligand-binding domain (ER) or JP8Bdel (ΔC)-AID-ER ($n$ = 3).

(B) Representative western blot analysis evaluating expression of AID-ER and ΔC-AID-ER in AID-KO or AID-WT cells. GAPDH served as a loading control.

(C) Quantification of IgA$^+$ cells 24 h post-CIT stimulation in *Med14*- and *Med19*-knockdown cells ($n$ = 3).

(D) Quantification of IgA$^+$ cells 24 h ($n$ = 4) and 40 h ($n$ = 3) post-CIT stimulation in *Med19*-knockdown cells expressing single or dual sgRNAs.

(E) Reverse transcription-quantitative PCR (RT-qPCR) analysis evaluating *Med14* (left) and *Med19* (right) mRNA levels normalized to *B2m*. Expression levels in control samples were set to 1.0 ($n$ = 3).

(F) RT-qPCR analysis evaluating *Med19* mRNA levels normalized to *B2m*. Expression levels in control samples were set to 1.0 ($n$ = 4).

(G−I) Prediction of intrinsically disordered regions in MED19: propensity for intrinsic disorder in *Arabidopsis thaliana* Med19a (G) and Med19b (H) and human MED19 (I) evaluated using the PONDR VSL2 predictor.[123] In human Med19, amino acids corresponding to deleted motifs (Figure 4A) are highlighted (PR, green; KNH, pink; KR1 and KR2, blue).

(J) Representative western blot analysis showing expression of exogenous WT and mutant human MED19 in CH12 cells.

(K) Southern blot images from an independent experiment analyzing *IgH*/*c-Myc* translocations in *Med19*-knockdown cells rescued by WT and Δ3KR MED19. The main dataset is presented in Figure 4K and 4L.

(L) Scatter plots showing pairwise correlations of $\log_2$-transformed gene expression levels [$\log_2$(TPM + 1)] between biological replicates ($n$ = 2). Lower triangle displays scatter plots, and upper triangle indicates Pearson's correlation coefficients ($R^2$).

(M) RT-qPCR analysis evaluating host gene levels of snoRNAs normalized to *B2m*. Expression levels in control samples were set to 1.0 ($n$ = 4). *Gsn* served as a gene negatively regulated by MED19 [baseMean = 8956, $\log_2$(Fold Change of sg*Med19*/Ctrl) = 2.358, $P_{\mathrm{adj}}$ = 1.57E-35].

Data represent the mean ± SEM. Dashed horizontal lines indicate normalized control levels (1.0). *$P_{adj} < 0.05$, **$P_{adj} < 0.01$, ***$P_{adj} < 0.001$; ns, not significant. See STAR Methods for statistical details.

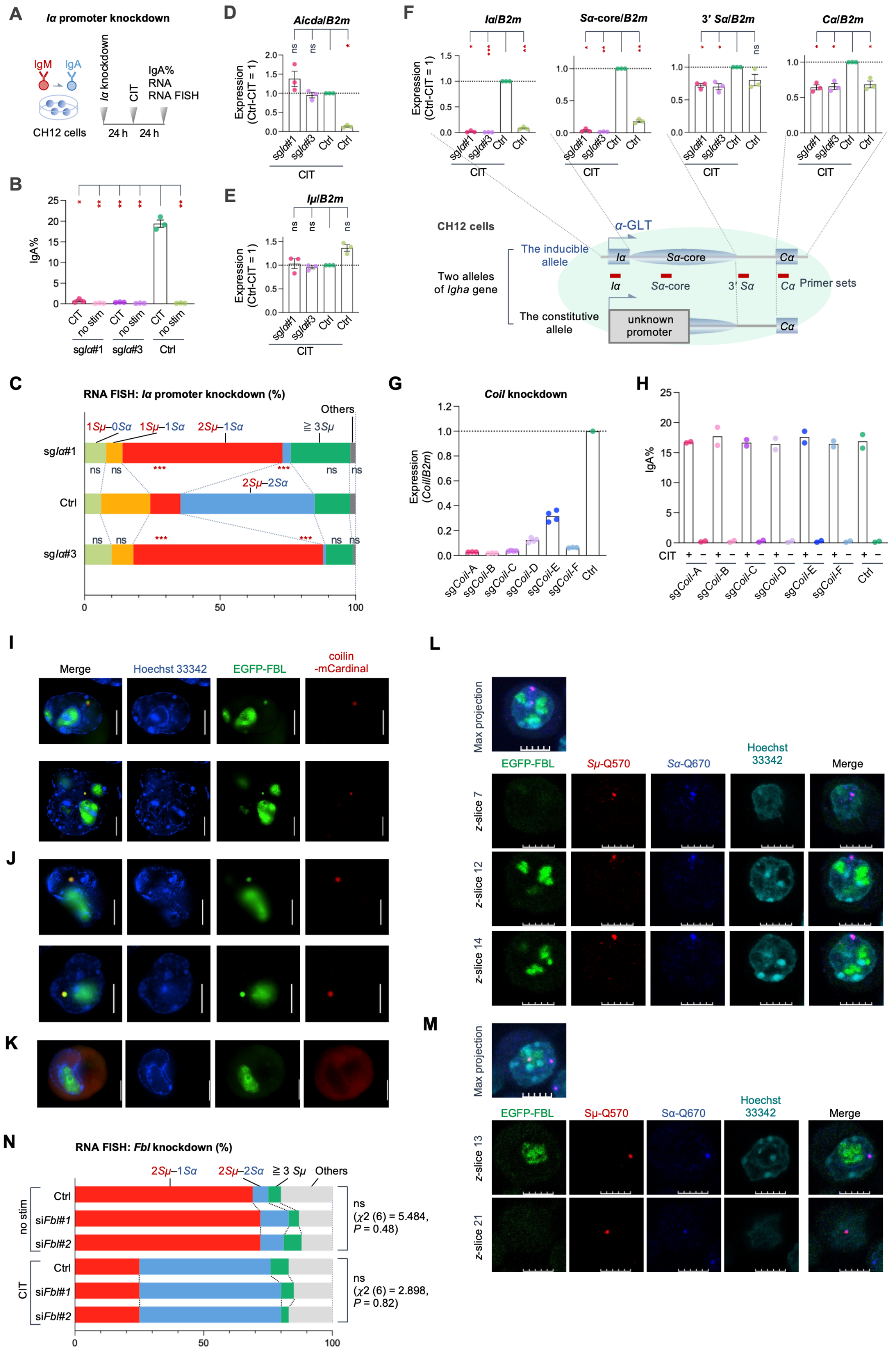
A
Iα promoter knockdown
IgM
IgA
CH12 cells
Iα knockdown
CIT
24 h
24 h
IgA%
RNA
RNA FISH
B
IgA%
CIT
no stim
sgIα#1
sgIα#3
Ctrl
C
RNA FISH: Iα promoter knockdown (%)
1Sμ–0Sα
1Sμ–1Sα
2Sμ–1Sα
≧ 3Sμ
Others
2Sμ–2Sα
sgIα#1
Ctrl
sgIα#3
D
Aicda/B2m
Expression (Ctrl-CIT = 1)
E
Iμ/B2m
F
Iα/B2m
Sα-core/B2m
3' Sα/B2m
Cα/B2m
CH12 cells
α-GLT
The inducible allele
Two alleles of Igha gene
The constitutive allele
unknown promoter
Iα
Sα-core
3' Sα
Cα
Primer sets
G
Coil knockdown
Expression (Coil/B2m)
sgCoil-A
sgCoil-B
sgCoil-C
sgCoil-D
sgCoil-E
sgCoil-F
Ctrl
H
IgA%
I
Merge
Hoechst 33342
EGFP-FBL
coilin -mCardinal
J
K
L
Max projection
EGFP-FBL
Sμ-Q570
Sα-Q670
Hoechst 33342
Merge
z-slice 7
z-slice 12
z-slice 14
M
Max projection
z-slice 13
z-slice 21
N
RNA FISH: Fbl knockdown (%)
2Sμ–1Sα
2Sμ–2Sα
≧ 3 Sμ
Others
no stim
Ctrl
siFbl#1
siFbl#2
CIT
ns
(χ2 (6) = 5.484, P = 0.48)
ns
(χ2 (6) = 2.898, P = 0.82)

**Figure S6. Characterization of constitutive *Sα* transcription and spatial distinction of GLT condensates from nuclear bodies, Related to Figure 5**

(A) Experimental workflow for CRISPR interference (CRISPRi)-mediated knockdown of the *Iα* promoter.

(B) Quantification of IgA$^+$ cells following *Iα* knockdown.

(C) Quantification of *Sμ*- and *Sα*-GLT foci detected by RNA FISH following *Iα* knockdown in CIT-stimulated cells expressing sg*Iα*#1, sg*Iα*#3, or non-targeting control. Prior to CIT stimulation, CH12 cells exhibited a single focus per cell co-localizing *Sμ* and *Sα* (see Figure 5L). Upon CIT stimulation, the proportion of cells displaying two *Sα* foci markedly increased, directly correlating with active CSR (Figure 5L, CIT vs. unstimulated). Among the two 3′ *Sα* RNA FISH foci observed upon stimulation, one was responsive to CIT and *Iα* knockdown, whereas the other was entirely non-responsive.

(D–F) RT-qPCR analysis evaluating transcript levels of *Aicda* (D), *Iμ* (E), and *Iα*, *Sα*-core, 3′ *Sα*, and *Cα* (F) normalized to *B2m* ($n$ = 3). While RT-qPCR signals spanning the upstream *Iα* to the *Sα*-core region were strictly responsive to both *Iα* knockdown and CIT, a major fraction (over 60%) of the RT-qPCR signals originating from the 3′ *Sα*-to-*Cα* region was constitutively expressed and refractory to either treatment, with only the remaining fraction (~40%) being responsive to CIT stimulation and *Iα* knockdown. These findings indicate that an allele-specific deletion of the *Iα*-to-*Sα*-core region on the non-switching allele replaces the native promoter with a constitutive active, non-responsive aberrant promoter in CH12 cells. Because steady-state RNA abundance reflects local promoter strength, this non-switching allele accounts for the dominant baseline 3′ *Sα*–*Cα* transcription, serving as a cell-line-specific background uncoupled from productive IgA switching.

(G and H) CRISPRi-mediated knockdown of *Coil*: *Coil* mRNA levels evaluated by RT-qPCR and normalized to *B2m* (G; $n$ = 1, with 4 technical replicates) and quantification of IgA$^+$ cells upon *Coil* knockdown (H; $n$ = 2). No statistical hypothesis testing was performed.

(I–K) Representative live-cell images showing subcellular localization of EGFP-FBL and coilin-mCardinal in CIT-stimulated (I) and unstimulated (J) CH12 cells, or EGFP-FBL with empty mCardinal vector control (K). Images of I−K were acquired using a Nikon N-SIM S super-resolution microscope without fixation.

(L and M) Representative images showing EGFP-FBL localization distinct from *Sμ*- and *Sα*-GLT RNA FISH foci.

(N) RNA FISH analysis of *Sμ*- and *Sα*-GLT foci following *Fbl* knockdown in unstimulated (no stim) and CIT-stimulated CH12 cells transfected with si*Fbl*#1, si*Fbl*#2, or Control siRNA.

Scale bars, 5 μm. Data represent the mean ± SEM, except in (G) and (H), where individual values are plotted as dots. Dashed horizontal lines indicate the normalized control level (1.0). *$P_{adj}$ < 0.05, **$P_{adj}$ < 0.01, ***$P_{adj}$ < 0.001; ns, not significant. See STAR Methods for statistical details.

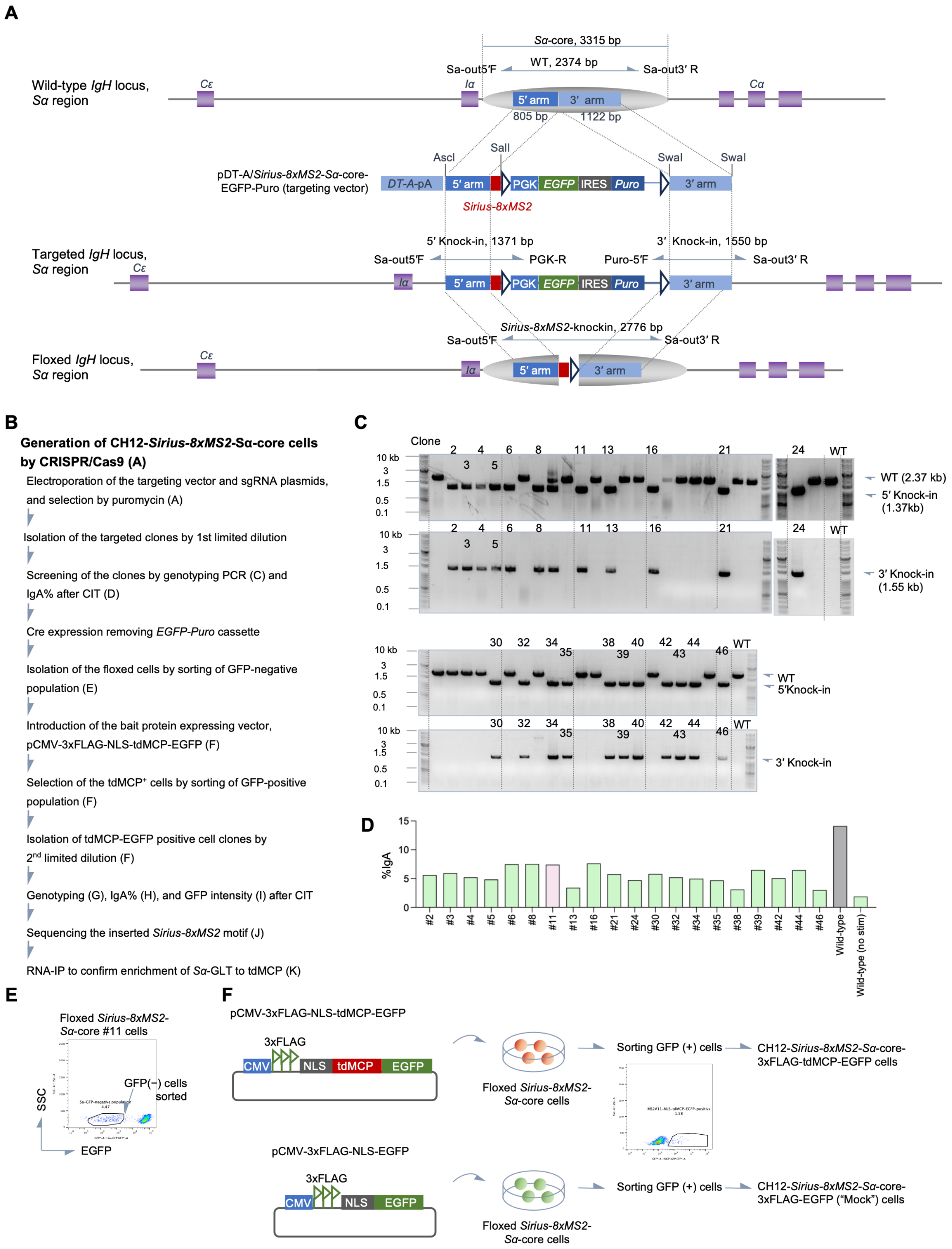
A
Sα-core, 3315 bp
WT, 2374 bp
Sa-out5′F
Sa-out3′ R
Wild-type IgH locus,
Sα region
Cε
Iα
Cα
5′ arm
3′ arm
805 bp
1122 bp
AscI
SalI
SwaI
SwaI
pDT-A/Sirius-8xMS2-Sα-core-
EGFP-Puro (targeting vector)
DT-A-pA
PGK
EGFP
IRES
Puro
Sirius-8xMS2
5′ Knock-in, 1371 bp
3′ Knock-in, 1550 bp
Sa-out5′F
PGK-R
Puro-5′F
Sa-out3′ R
Targeted IgH locus,
Sα region
Sirius-8xMS2-knockin, 2776 bp
Floxed IgH locus,
Sα region
B
Generation of CH12-Sirius-8xMS2-Sα-core cells
by CRISPR/Cas9 (A)
Electroporation of the targeting vector and sgRNA plasmids, and selection by puromycin (A)
Isolation of the targeted clones by 1st limited dilution
Screening of the clones by genotyping PCR (C) and IgA% after CIT (D)
Cre expression removing EGFP-Puro cassette
Isolation of the floxed cells by sorting of GFP-negative population (E)
Introduction of the bait protein expressing vector, pCMV-3xFLAG-NLS-tdMCP-EGFP (F)
Selection of the tdMCP⁺ cells by sorting of GFP-positive population (F)
Isolation of tdMCP-EGFP positive cell clones by 2nd limited dilution (F)
Genotyping (G), IgA% (H), and GFP intensity (I) after CIT
Sequencing the inserted Sirius-8xMS2 motif (J)
RNA-IP to confirm enrichment of Sα-GLT to tdMCP (K)
C
Clone
10 kb
3
1.5
0.5
0.1
WT
WT (2.37 kb)
5′ Knock-in
(1.37kb)
3′ Knock-in
(1.55 kb)
WT
5′Knock-in
3′ Knock-in
D
%IgA
#2
#3
#4
#5
#6
#8
#11
#13
#16
#21
#24
#30
#32
#34
#35
#38
#39
#42
#44
#46
Wild-type
Wild-type (no stim)
E
Floxed Sirius-8xMS2-
Sα-core #11 cells
GFP(−) cells
sorted
SSC
EGFP
F
pCMV-3xFLAG-NLS-tdMCP-EGFP
3xFLAG
CMV
NLS
tdMCP
EGFP
Floxed Sirius-8xMS2-
Sα-core cells
Sorting GFP (+) cells
CH12-Sirius-8xMS2-Sα-core-
3xFLAG-tdMCP-EGFP cells
pCMV-3xFLAG-NLS-EGFP
CH12-Sirius-8xMS2-Sα-core-
3xFLAG-EGFP (“Mock”) cells

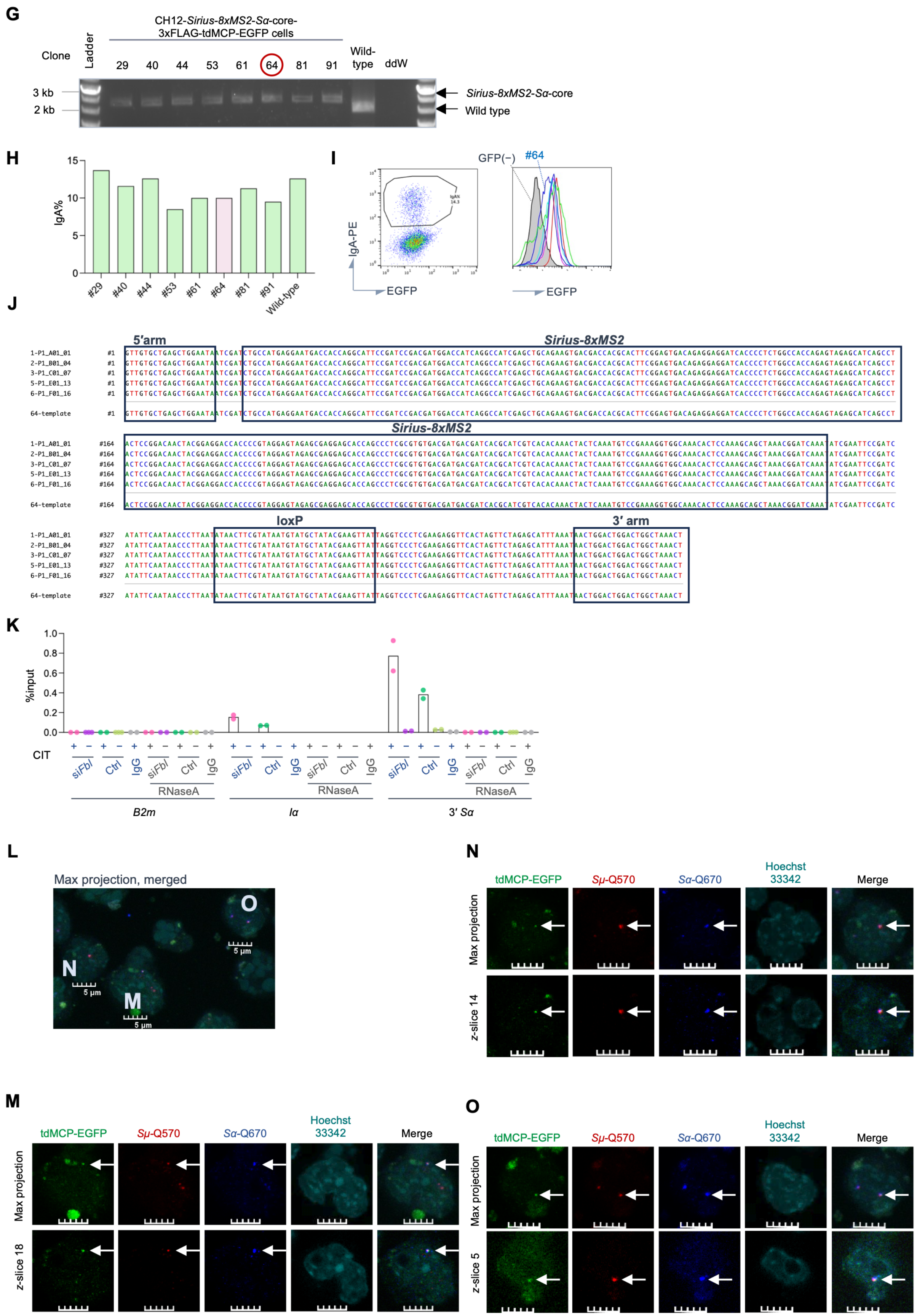

G
CH12-Sirius-8xMS2-Sα-core-3xFLAG-tdMCP-EGFP cells
Clone
Ladder
29 40 44 53 61 64 81 91
Wild-type
ddW
3 kb
2 kb
Sirius-8xMS2-Sα-core
Wild type
H
IgA%
15
10
5
0
#29 #40 #44 #53 #61 #64 #81 #91 Wild-type
I
IgA-PE
EGFP
GFP(−)
#64
EGFP
J
5′arm
Sirius-8xMS2
loxP
3′ arm
1-P1_A01_01
2-P1_B01_04
3-P1_C01_07
5-P1_E01_13
6-P1_F01_16
64-template
K
%input
1.0
0.8
0.6
0.4
0.2
0.0
CIT
siFbl
Ctrl
IgG
RNaseA
B2m
Iα
3′ Sα
L
Max projection, merged
O
N
M
5 µm
N
tdMCP-EGFP
Sμ-Q570
Sα-Q670
Hoechst 33342
Merge
Max projection
z-slice 14
M
tdMCP-EGFP
Sμ-Q570
Sα-Q670
Hoechst 33342
Merge
Max projection
z-slice 18
O
tdMCP-EGFP
Sμ-Q570
Sα-Q670
Hoechst 33342
Merge
Max projection
z-slice 5

**si*Fbl*-induced GOCC changes in *Sα*-Proteome**

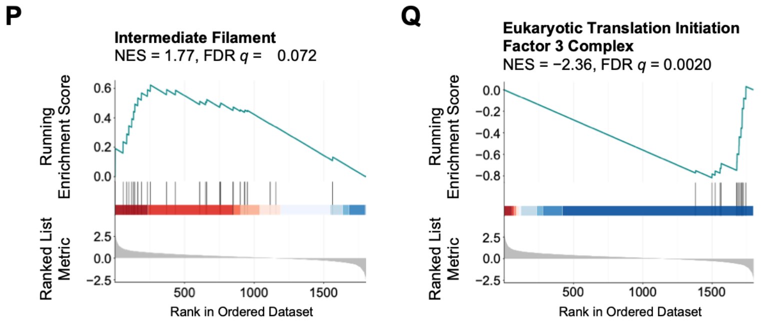


**si*Fbl*-Induced *Sα*-GLT Proteome Changes**

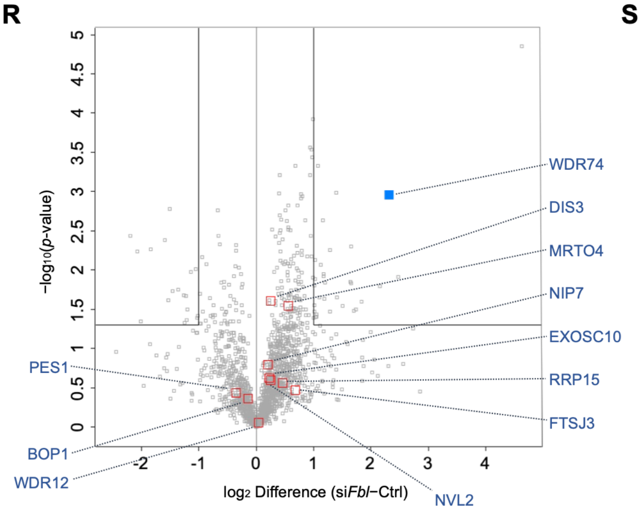


**S**

**Pre-60S Biogenesis Factor Changes upon si*Fbl***

| Functional Category | Molecules | $\log_2$ Difference (si*Fbl*−Ctrl) |
|---|---|---|
| Early pre-60S assembly | NVL2 (Rix7), FTSJ3, NIP7, RRP15 | 0.199–0.684 |
| PeBoW core complex | BOP1, PES1, WDR12 | ≤ 0.039 |
| WDR74 structural module | RPF1, MAK16, RRP1 | (not detected) |
| Intermediate/Late nuclear maturation | MRTO4 | 0.556 |
| RNA exosome & processing machinery | EXOSC1-10, DIS3 | 0.226–0.594 |
| | MTR4 | (not detected) |

**Figure S7. Generation and validation of *Sirius-8xMS2* knock-in CH12 cells for *Sα*-GLT interactome profiling, Related to Figure 6**

(A) Schematic illustration of *Sirius-8xMS2* cassette targeted integration into the *Sα* region of the *IgH* locus using a targeting vector containing a PGK-EGFP-IRES-Puro selection cassette.

(B) Experimental workflow for generating CH12 cells harboring the *Sirius-8xMS2* sequence in the *Sα*-core region and expressing the bait protein 3xFLAG-tdMCP-EGFP.

(C) Genotyping PCR confirming the targeted integration of the *Sirius-8xMS2* sequence and PGK-EGFP-IRES-Puro cassette into the *Sα*-core region of candidate clones.

(D) CSR efficiency quantified by the percentage of $IgA^+$ cells in candidate knock-in clones 24 h post-CIT stimulation ($n$ = 1).

(E) Representative flow cytometric plot showing gating of EGFP-negative cells following transient Cre recombinase expression to excise the selection cassette.

(F) Schematic representation of the pCMV-3xFLAG-tdMCP-EGFP construct containing a nuclear localization signal (NLS). Stable introduction of this plasmid into floxed *Sirius-8xMS2-Sα*-core cells generated CH12-*Sirius-8xMS2-Sα*-core-3xFLAG-tdMCP-EGFP line. As a control (EGFP alone/mock), introduction of a 3xFLAG-EGFP vector generated CH12-*Sirius-8xMS2-Sα*-core-3xFLAG-EGFP cells.

(G) Genotyping PCR validating retention of the *Sirius-8xMS2* insertion after deletion of the PGK-EGFP-IRES-Puro cassette in single-cell clones following tdMCP introduction.

(H) CSR efficiency ($IgA^+$ %) of selected clones following stable introduction of 3xFLAG-tdMCP-EGFP (*n* = 1). Clone #64, used in subsequent experiments, is highlighted in pink.

(I) Flow cytometric evaluation of EGFP expression levels across selected clones. Clone #64 was selected for downstream analyses because of its moderate EGFP-tdMCP expression level, minimizing nonspecific background enrichment.

(J) Sanger sequencing alignment of clone #64 confirming intact insertion of the *Sirius-8xMS2* sequence into the *Sα*-core region. The 444-bp PCR amplicon derived from the PCR product shown in (G) was subcloned into pGEM-T Easy Vector and subjected to Sanger sequencing. All six clones were derived from the genomic DNA of the clone #64.

(K) RIP-qPCR validation demonstrating specific capture of *Sα* GLTs following anti-FLAG immunoprecipitation (*n* = 1 with technical duplicates). Data represent the mean with individual values plotted as dots. No statistical hypothesis testing was performed.

(L–O) Representative confocal images showing co-localization of tdMCP-EGFP foci with *Sμ*- and *Sα*-GLT RNA FISH signals in CIT-stimulated cells. See also Figure 6C.

(P and Q) GSEA enrichment plots of *Sirius-8xMS2*-mediated *Sα*-GLT proteome alterations upon *Fbl* knockdown for "Intermediate Filament" (P) and "Eukaryotic Translation Initiation Factor 3 Complex" (Q).

(R and S) Volcano plot (R) and summary table (S) demonstrating the selective enrichment of WDR74 relative to other pre-60S ribosomal subunit biogenesis factors upon *Fbl* knockdown in the *Sα*-GLT proteome. Proteins were categorized based on established structural and functional studies.[80,83–85]

See STAR Methods for statistical details.

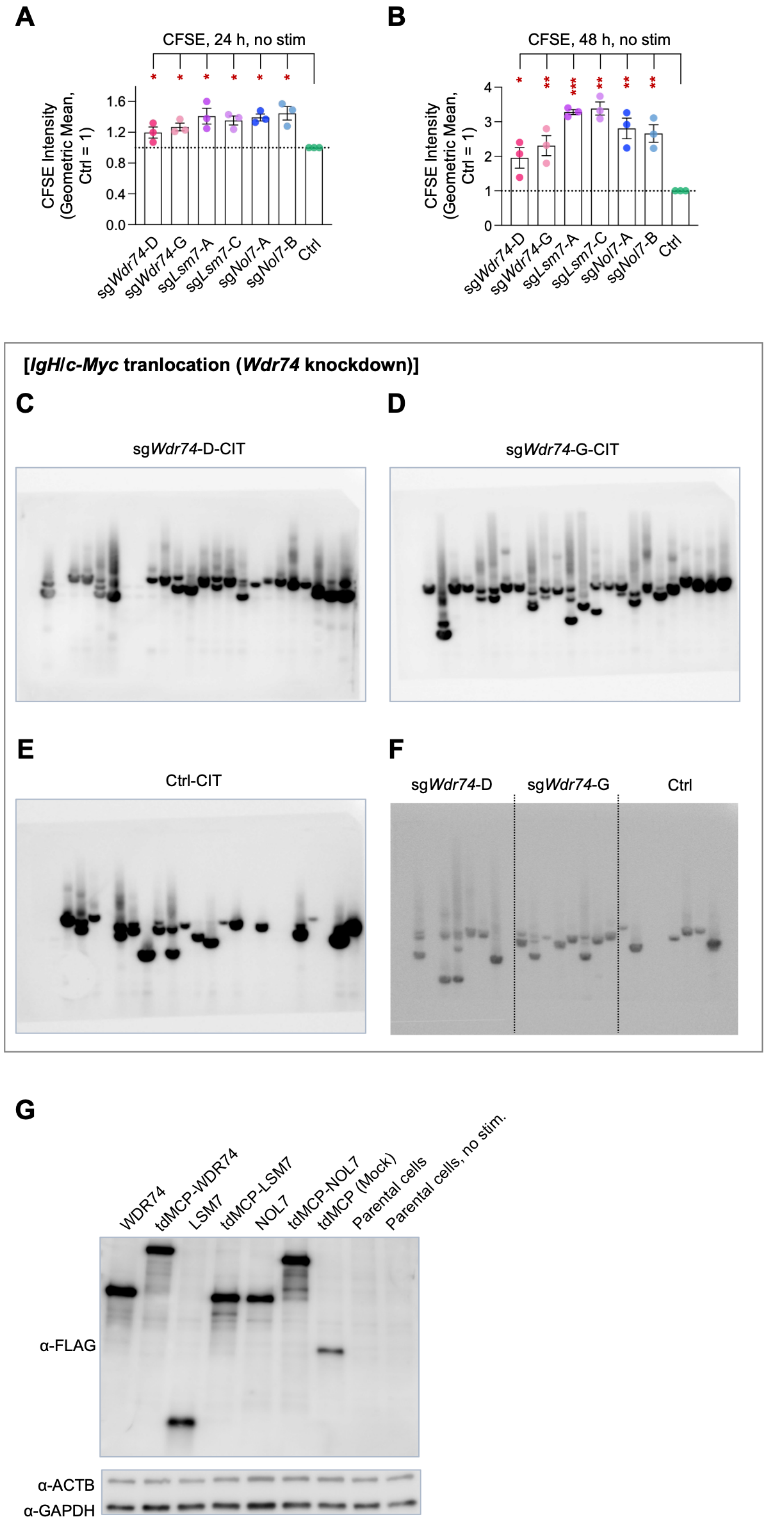
A
CFSE, 24 h, no stim
CFSE Intensity (Geometric Mean, Ctrl = 1)
1.6
1.2
0.8
0.4
0.0
sgWdr74-D
sgWdr74-G
sgLsm7-A
sgLsm7-C
sgNol7-A
sgNol7-B
Ctrl
B
CFSE, 48 h, no stim
CFSE Intensity (Geometric Mean, Ctrl = 1)
4
3
2
1
0
sgWdr74-D
sgWdr74-G
sgLsm7-A
sgLsm7-C
sgNol7-A
sgNol7-B
Ctrl
[IgH/c-Myc tranlocation (Wdr74 knockdown)]
C
sgWdr74-D-CIT
D
sgWdr74-G-CIT
E
Ctrl-CIT
F
sgWdr74-D
sgWdr74-G
Ctrl
G
WDR74
tdMCP-WDR74
LSM7
tdMCP-LSM7
NOL7
tdMCP-NOL7
tdMCP (Mock)
Parental cells
Parental cells, no stim.
α-FLAG
α-ACTB
α-GAPDH

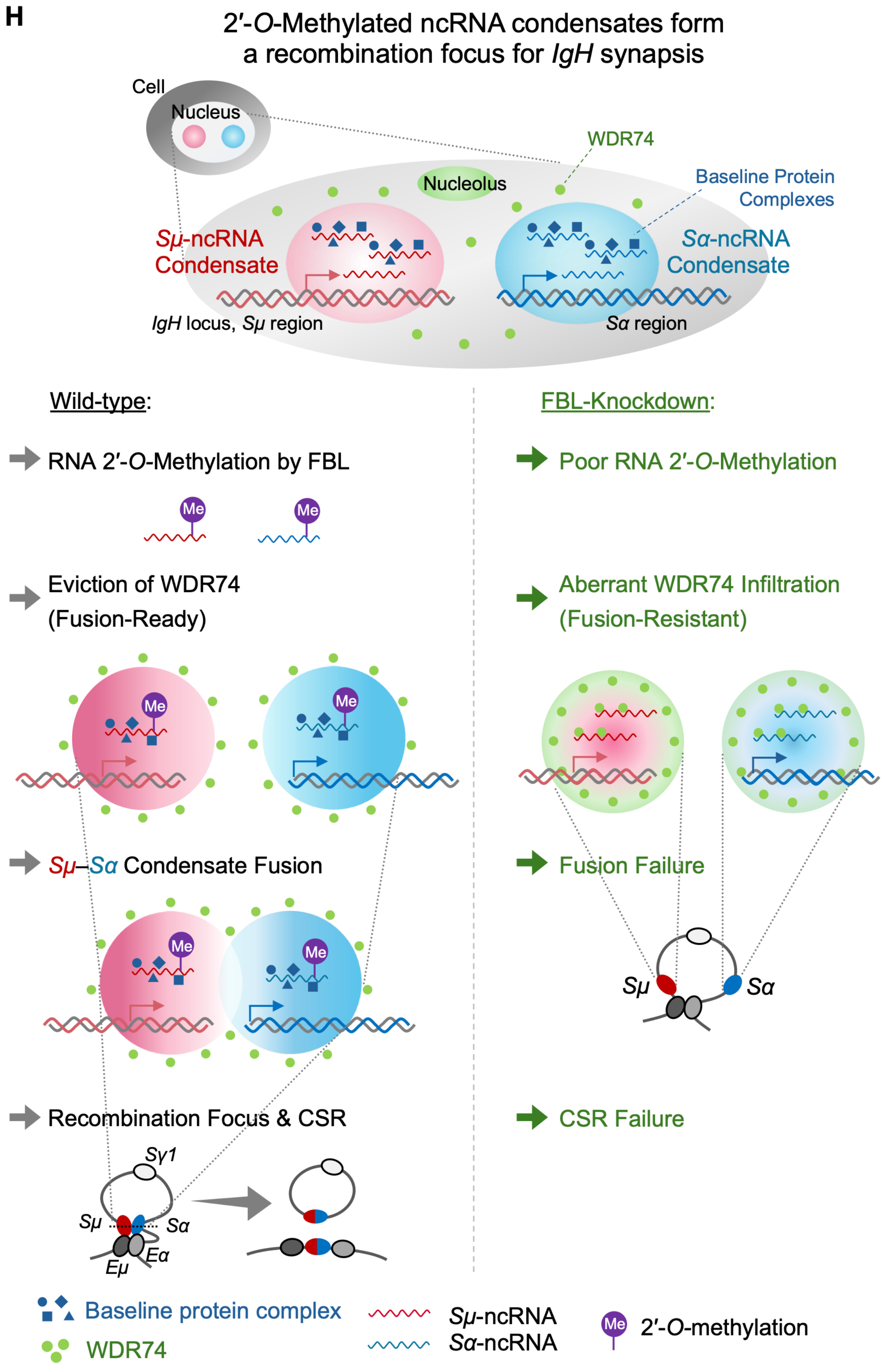
H
2′-O-Methylated ncRNA condensates form a recombination focus for IgH synapsis
Cell
Nucleus
WDR74
Nucleolus
Baseline Protein Complexes
Sμ-ncRNA Condensate
Sα-ncRNA Condensate
IgH locus, Sμ region
Sα region
Wild-type:
RNA 2′-O-Methylation by FBL
Me
Eviction of WDR74 (Fusion-Ready)
Sμ–Sα Condensate Fusion
Recombination Focus & CSR
Sγ1
Sμ
Sα
Eμ
Eα
FBL-Knockdown:
Poor RNA 2′-O-Methylation
Aberrant WDR74 Infiltration (Fusion-Resistant)
Fusion Failure
CSR Failure
Baseline protein complex
WDR74
Sμ-ncRNA
Sα-ncRNA
2′-O-methylation

**Figure S8. Functional characterization and validation of *Wdr74*-, *Lsm7*-, or *Nol7*-manipulated cells, Related to Figure 7**

(A and B) Cell proliferation assessed by CFSE fluorescence intensity in unstimulated *Wdr74*-, *Lsm7*-, *or Nol7*-knockdown cells at 24 h (A) and 48 h (B) in culture, normalized to control ($n$ = 3).

(C–F) Southern blot analysis from a second independent experiment evaluating *IgH*/*c-Myc* translocation in *Wdr74*-knockdown cells. See also Figures 7M and 7N.

(G) Western blot analysis validating the expression of FLAG-tagged WDR74, LSM7, and NOL7 overexpression constructs in *Sirius-8xMS2* knock-in cells. See also Figure 7S.

(H) Working model for FBL-dependent ncRNA 2′-*O*-methylation in regulating *Sμ–Sα* condensate dynamics and CSR. Top: Spatial representation of *Sμ*- and *Sα*-ncRNA condensates in the nucleus. Bottom: (Left) In wild-type cells, FBL-catalyzed 2′-*O*-methylation promotes WDR74 eviction, driving *Sμ–Sα* condensate fusion and productive synapsis. (Right) Upon FBL knockdown, reduced methylation leads to aberrant WDR74 infiltration, resulting in a fusion-resistant state, synapsis failure and CSR failure.

Data represent the mean ± SEM. Dashed horizontal lines indicate normalized control levels (1.0).
*$P_{adj}$ < 0.05, **$P_{adj}$ < 0.01, ***$P_{adj}$ < 0.001. See STAR Methods for statistical details.

**Tables S1–S4**

**Table S1. Mutation Numbers in the SHM analysis, Related to Figure 2F**

| treatment | cell | Total clones | Mutated clones | Mutation number | Total base number | Mutation frequency (x10$^{-4}$) | Del bp/ clone# | Ins bp/ clone# |
|---|---|---|---|---|---|---|---|---|
| no stim | Ctrl | 90 | 2 | 2 | 35,910 | 0.557 | 0 bp/0 | 0 bp/0 |
| CIT | Ctrl | 90 | 14 | 18 | 35,910 | 5.01 | 113 bp/2 | 0 bp/0 |
| | si*Fbl*#1 | 88 | 11 | 14 | 35,112 | 4 | 25 bp/1 | 0 bp/0 |
| | si*Fbl*#2 | 89 | 14 | 18 | 35,511 | 5.07 | 41 bp/3 | 0 bp/0 |

**Table S2. Mutation Patterns in the SHM analysis, Related to Figure 2F**

**Mutation pattern of SHM**

Ctrl-no stim

| From \ To | G | C | A | T | Sum |
|---|---|---|---|---|---|
| G | | 0 | 1 | 0 | 1 |
| C | 0 | | 0 | 0 | 0 |
| A | 1 | 0 | | 0 | 1 |
| T | 0 | 0 | 0 | | 0 |
| | | | | | 2 |

Ctrl-CIT

| From \ To | G | C | A | T | Sum |
|---|---|---|---|---|---|
| G | | 0 | 4 | 1 | 5 |
| C | 0 | | 0 | 10 | 10 |
| A | 1 | 0 | | 0 | 1 |
| T | 0 | 1 | 1 | | 2 |
| | | | | | 18 |

si*Fbl*#1-CIT

| From \ To | G | C | A | T | Sum |
|---|---|---|---|---|---|
| G | | 1 | 4 | 2 | 7 |
| C | 0 | | 0 | 6 | 6 |
| A | 0 | 1 | | 0 | 1 |
| T | 0 | 0 | 0 | | 0 |
| | | | | | 14 |

si*Fbl*#2-CIT

| From \ To | G | C | A | T | Sum |
|---|---|---|---|---|---|
| G | | 1 | 6 | 4 | 11 |
| C | 2 | | 0 | 3 | 5 |
| A | 0 | 0 | | 0 | 0 |
| T | 1 | 1 | 0 | | 2 |
| | | | | | 18 |

**Table S3. Highly expressed and significantly enhanced by MED19, Related to Figure 4N**

| Gene name | base Mean | Log$_2$(Fold Change) | Adjusted *p*-value | snoRNAs hosted by this gene |
|---|---|---|---|---|
| *2410006H16Rik* | 3265.83 | -2.81 | 8.38E-41 | Snord49b, Snord49a, Snord65 |
| *Chac1* | 5563.32 | -2.85 | 1.20E-67 | none |
| *Cox6a2* | 2299.21 | -4.85 | 4.19E-126 | none |
| *Cth* | 4562.98 | -3.23 | 9.00E-81 | none |
| *Ddit3* | 2873.74 | -2.18 | 2.71E-36 | none |
| *Fgf21* | 1690.60 | -3.43 | 7.95E-65 | none |
| *H2ac12* | 4561.20 | -2.13 | 4.23E-21 | none |
| *H2ac4* | 4821.73 | -2.10 | 1.01E-21 | none |
| *H2ac8* | 5720.18 | -2.10 | 1.28E-24 | none |
| *Ighg3* | 3589.37 | -2.77 | 2.06E-35 | none |
| *Iglc1* | 1682.90 | -3.68 | 1.95E-57 | none |
| *Nupr1* | 2399.27 | -2.44 | 5.56E-31 | none |
| *Pvt1* | 2888.06 | -2.10 | 1.81E-34 | none |
| *Reep6* | 1768.07 | -2.61 | 1.07E-34 | none |
| *Snhg1* | 1924.31 | -2.45 | 9.74E-26 | Snod22 |
| *Snhg16* | 1455.47 | -2.20 | 5.50E-23 | Snord1c, Snord1b, Snord1a |
| *Snhg5* | 2890.16 | -2.44 | 2.83E-40 | Snord50a, Snord50b |
| *Tmx2* | 2237.53 | -6.87 | 1.06E-13 | none |
| *Trib3* | 5452.14 | -3.02 | 6.40E-74 | none |
| *Tsc22d3* | 2359.15 | -2.33 | 5.03E-35 | none |
| *Wnt10a* | 1515.90 | -2.10 | 1.87E-15 | none |
| *Zfas1* | 1941.79 | -2.02 | 6.28E-16 | Snord12 |

Genes with a normalized expression level (baseMean) ≥ 1000 and $\log_2$[Fold Change (sg*Med19*/Ctrl)] < −2 are listed.

**Table S4. *Sα*-GLT binding proteins regulated by FBL, Related to Figure 6H**

| | | | | Enriched by si*Fbl* | | | Depleted by si*Fbl* | | | |
|---|---|---|---|---|---|---|---|---|---|---|
| Protein Name | | | | LSM7 | WDR74 | NOL7 | SF3B3 | SF3B1 | PHF5A | SF3B5 |
| *Sirius-8xMS2*-RIP-Proteome | Signal intensity of *Sα*-GLT binding | si*Fbl* | Set1 | 13.22 | 12.18 | 13.63 | 15.13 | 13.71 | 14.95 | 11.67 |
| | | | Set2 | 14.68 | 11.96 | 12.98 | 14.84 | 13.82 | 14.67 | 9.98 |
| | | | Set3 | 14.58 | 11.71 | 13.02 | 15.07 | 14.03 | 14.60 | 12.32 |
| | | Ctrl | Set1 | 10.72 | 9.51 | 10.95 | 15.76 | 14.11 | 15.38 | 11.68 |
| | | | Set2 | 10.06 | 10.02 | 10.43 | 15.43 | 14.97 | 15.17 | 12.07 |
| | | | Set3 | 14.00 | 9.37 | 11.78 | 15.52 | 14.27 | 15.35 | 12.13 |
| | | Mock | Set1 | 9.22 | 7.97 | 8.81 | 12.22 | 11.84 | 9.10 | 10.49 |
| | | | Set2 | 8.27 | 9.33 | 9.34 | 12.42 | 12.41 | 8.86 | 9.11 |
| | | | Set3 | 10.37 | 9.59 | 10.48 | 13.10 | 13.15 | 9.73 | 9.97 |
| | Welch's *t*-test (si*Fbl* vs. Ctrl) | $\log_2$ Difference | | 2.57 | 2.32 | 2.16 | -0.56 | -0.60 | -0.56 | -0.64 |
| | | $-\log_{10}$(P-value) | | 0.80 | 2.96 | 1.79 | 1.84 | 0.85 | 1.77 | 0.34 |
| | | FDR *q*-value | | 0.53 | 0.15 | 0.29 | 0.28 | 0.51 | 0.29 | 0.74 |
| Main Localization | | | | Nucleolus | | | U2snRNP | | | |
| References | | | | Kufel et al., 2003[81] | Hiraishi et al., 2018[80] | McCool et al., 2023[82] | Das et al., 1999[93] | | Will et al., 2002[94] | |
| FBL-interactome | FBL-CIT vs. Mock-CIT | Abundance Ratio | | 7.735 | not detected | 7.211 | 2.520 | 1.441 | 0.711 | not detected |
| | | Abundance Ratio *P* value | | 0.023 | | 0.031 | 0.013 | 0.124 | 0.631 | |
| | FBL-no stim vs. Mock-CIT | Abundance Ratio | | 7.734 | | 5.383 | 2.771 | 1.720 | 1.397 | |
| | | Abundance Ratio *P* value | | 0.023 | | 0.048 | 0.010 | 0.047 | 0.641 | |
| Knockdown effect on CSR | | | | This study | | | Decreased (Begum et al., 2021)[95] | | | |
| | | | | No change | Increased | Modestly decreased | | | | |

***Sirius-8xMS2*-RIP-Proteome**

- Signal intensity values of mass spectrometry data were $\log_2$-transformed and imputed from normal distribution
- Two-sample test (Welch's *t*-test, both sides, Benjamini-Hochberg FDR) comparing si*Fbl* and Control was performed ($n$ = 3).
- Imputed cells are highlighted in grey.

**FBL-interactome analysis**

- The co-immunoprecipitated protein fractions were obtained from three groups, CIT-stimulated 3xFLAG-tagged FBL cells (FBL-CIT), unstimulated 3xFLAG-tagged FBL cells (FBL-no stim), parental cells without FLAG protein (Mock-CIT) ($n$ = 2).
- Abundance Ratio and Abundance Ratio $P$ values are shown.
